# Predicting Small Molecule Ligand – RNA Binding Pocket Binding Modes Using Metadynamics

**DOI:** 10.1101/2023.10.04.560960

**Authors:** Zhixue Bai, Alan Chen

**Affiliations:** Department of Chemistry and the RNA Institute, University at Albany, State University of New York, Albany, NY, 12222, USA

**Keywords:** RNA binding pocket, small molecule ligand, molecular dynamics, metadynamics, upper wall restrain, free energy landscape, binding mode prediction

## Abstract

Understanding the structural dynamics of how small molecule ligand recognize its RNA binding pocket is always a crucial determinant in pharmaceutical research. Molecular dynamics (MD) simulation is often used to interpretate this process at atomic resolution. However, the insurmountable high energy barriers in the binding pathway results in the nonergodic dynamics for unbiased MD sampling. To address this limitation, we applied well-tempered metadynamics coupled with upper wall restrain in this work, therefore providing an novel modeling approach for sampling the multiple state transitions during this binding process and probing the most energy favorable binding modes through two-dimensional free energy landscape reconstructed by incorporating couple possible hydrogen binding interactions between small molecule ligand and its RNA binding pocket as collective variables (CVs). Our computational predictions of binding modes for all five cases studied are in quantitative agreement with structures solved by X-ray crystallography or NMR with RMSD less than 2.0 Å. In addition, we presented the first molecular dynamics binding pathway and binding mechanism for the three cases of in vitro selected RNA aptamer. Our study demonstrated that metadynamics can be applied to effectively sampling state transitions of ligand binding events. By coupling with upper wall restrain, we have enabled fast free energy profile calculation and binding mode prediction for small molecule-RNA binding process, facilitating RNA-ligand binding investigation. This method therefore could be much-needed in computer-aided drug design pipelines of RNA-targeted small molecule compounds.

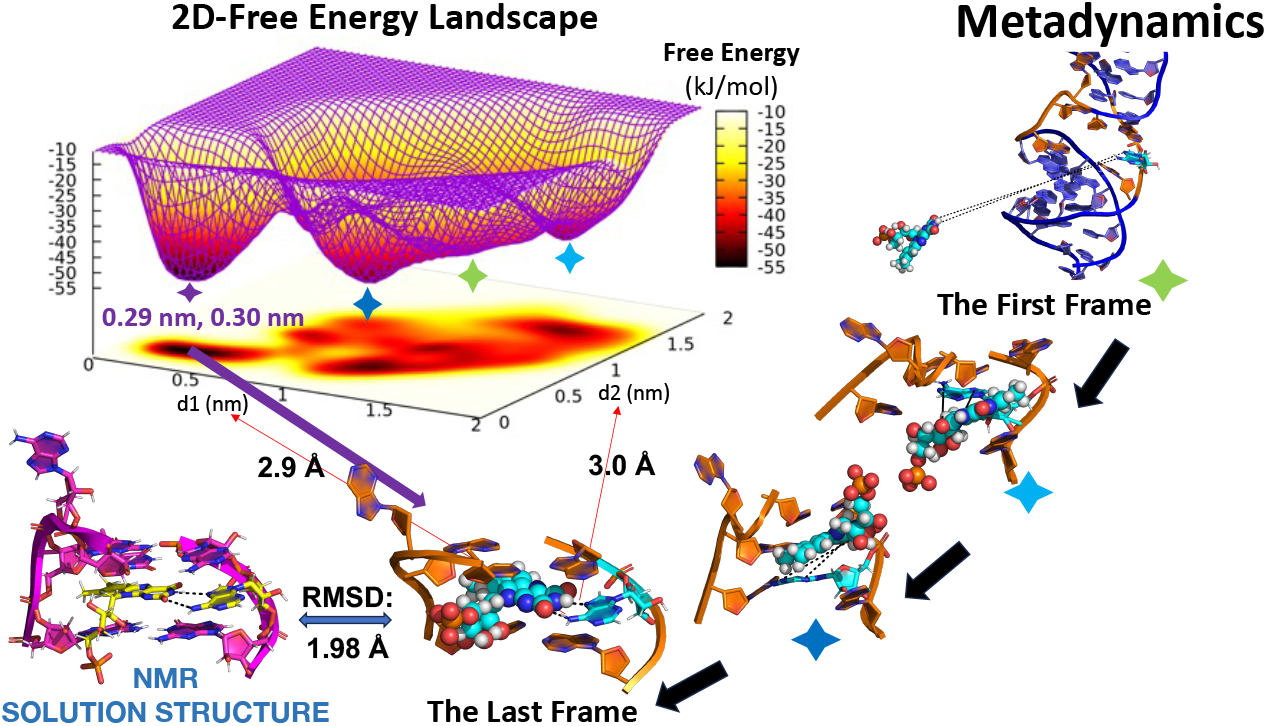

## INTRODUCTION

There are three types of well-known small molecule ligand-RNA binding pocket recognition cases. The high selectivity and high affinity (μM to nM) binding between RNA sequence identified by the techniques of in vitro selection and its small-molecule target including cofactors, antibiotics and therapeutic targets ^1–3^. The specific recognition between the high affinity aptamer domain of riboswitch located in the 5’ untranslated region of bacterial mRNAs and its small molecule ligand of metabolic intermediate or antimicrobial agent, which can lead to active or attenuated gene expression by altering accessibility to mRNA signals necessary for transcription or translation ^4–8^. The binding between small-molecule splicing modulator discovered as treatment for human genetic disease and single-stranded purine-rich RNA of pre-mRNA exon at 5’ splicing site, which can fix the weak base complementary pairing between exon and sequence of spliceosome thereby increasing the production of mature mRNA for relevant protein expression ^9–14^. Understanding the mechanism of the molecular recognition process between small molecule ligands and its RNA binding pocket is of great significance in drug discovery. Due to inherent flexibility and complex dynamics of structured RNAs, crystal structures of small molecule ligand-RNA complex are extremely difficult to solve through X-ray crystallography. While NMR is able to probe the structure of RNAs in solution, structures with long RNA sequence are often difficult to obtain due to the inherent technical problem of spectral overlapping ^15, 16^. In addition, both X-ray crystallography and NMR require considerable time and effort to solve a structure associated with RNA, and neither provided dynamic information related to RNA folding and molecular recognition.

Compared with experimental methods, computational molecular dynamics simulation is a powerful alternative method for elucidating structural dynamics and structure−function relationships of nucleic acid–small molecule ligand recognition at atomic resolution ^10, 17–20^, which determines the properties of system by integrating Newton’s equations of motion of atoms ^21^. There are many new developments for force fields of nucleic acid systems have been made in recent years including the correction of erroneously strong base stacking, base pairing, ionic interactions and parametrizations against experimental data ^7, 22–27^. However, the associated long time scale for simulating the real binding process is difficult to access via conventional unbiased MD simulations due to the high energy barriers needed to be overcome in the binding pathway ^28, 29^. In responds to this challenge, MD was used in connection with various of biasing algorithms to flatten free energy surface and accelerate sampling ^30^, including umbrella sampling ^31–33^, steered molecular dynamics ^17, 34, 35^, accelerated molecular dynamics ^10, 29, 36^, metadynamics ^19, 37–39^ and more. Compared to the other enhanced sampling methods, metadynamics is able to reconstruct free energy landscapes and accelerate state transition along multidimensional reaction coordinate with selectable collective variables. It also doesn’t require running multiple thermodynamics windows and laborious postprocessing like umbrella sampling. Metadynamics is biased from the history of experienced configurations by a sum of Gaussian contributions centered along the trajectory of selected CVs. This process of filling local minima with Gaussian potential discourages revisiting of experienced states and encourages exploring new conformational space, which brings the system cross the free energy barriers and realizing conformational transitions. Moreover, the well-tempered metadynamics is able to limit the extent of exploration of the free energy landscape via bias factor and avoid irreversible sampling ^40^. The features of metadynamics are especially useful for docking small molecule ligand on its flexible receptor ^7,39^.

Here we provide a new sampling strategy for predicting binding modes of small molecule ligand-RNA binding pocket system using well-tempered metadybamics coupled with upper wall restrain. This method can incorporate hydrogen binding information from experimental results to assess free energy landscape of the whole binding process of the recognition between small molecule ligand and its RNA binding pocket including barriers and intermediate minima and accurately probe the most energy favorable binding mode. To demonstrate the validity of the methodology, we studied five cases of high biological interests: (i) flavin mononucleotide (FMN) bound with 35-nucleotide RNA aptamer identified through in vitro selection ^3^, (ii) preQ1 riboswitch aptamer in the metabolite-bound state ^5^, (iii) ligand-bound adenine riboswitch aptamer domain ^6^, (iv) RNA aptamer complexed with citrulline and (v) RNA aptamer complexed with arginine ^11^ (Figure 1) (Figure 2). In each case, our simulation prediction of binding mode had quantitative agreement with experimental geometry solved by X-ray crystallography or NMR. We also presented the first molecular dynamics binding pathway and binding mechanism for (i), (iv) and (v). The good agreement between computational prediction and experimentally solved structure and the molecular dynamics insights provided in our study announced an important advance in RNA-ligand binding investigations and demonstrated that metadynamics can effectively sample state transitions in ligand binding process. This method could therefore be potentially valuable in computer-assisted drug design pipelines of RNA-targeted small molecule compounds.

**Figure 1.**
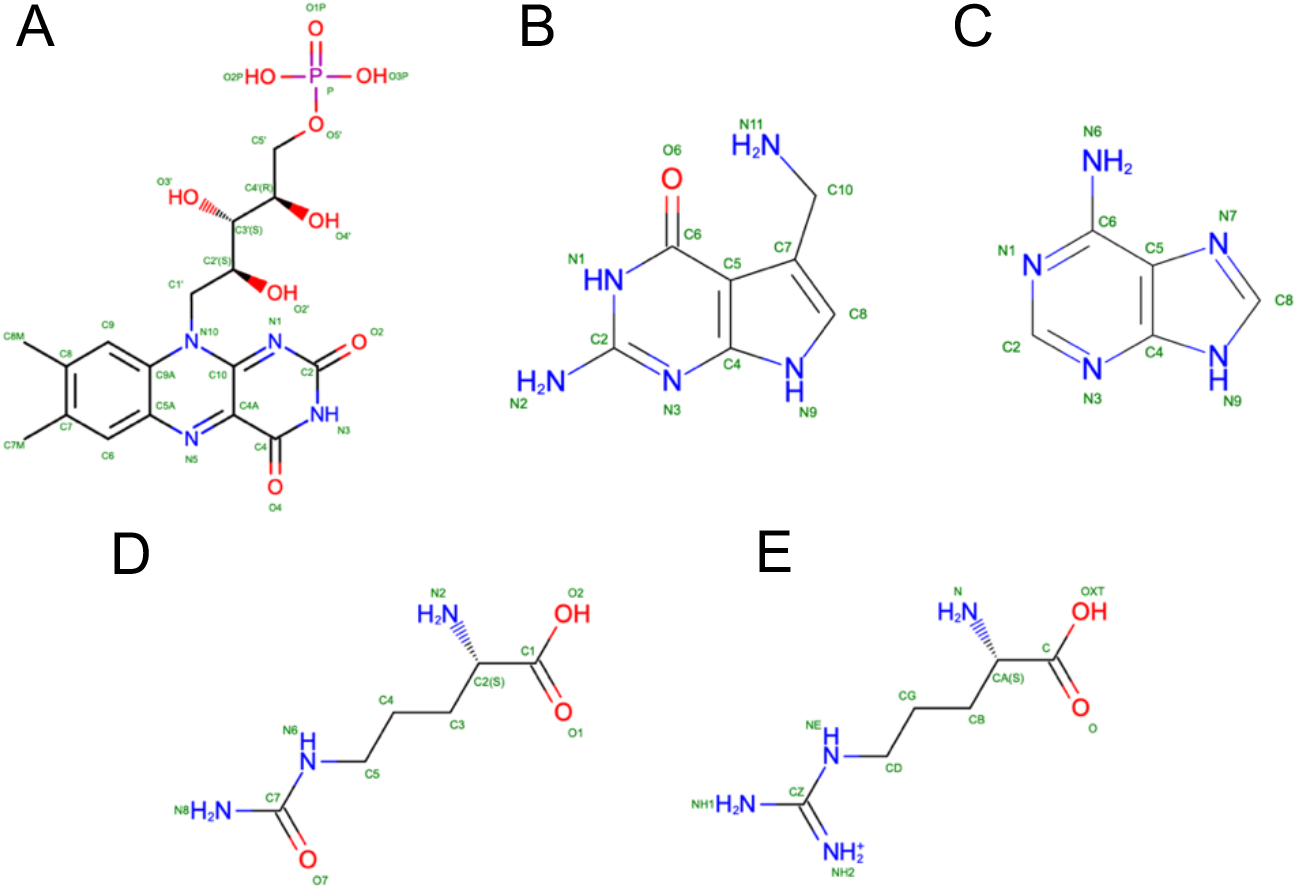
Small molecule ligand chemical structure of **(A)** flavin mononucleotide (FMN); **(B)** PreQ1; **(C)** adenine; **(D)** citrulline and **(E)** arginine.

**Figure 2.**
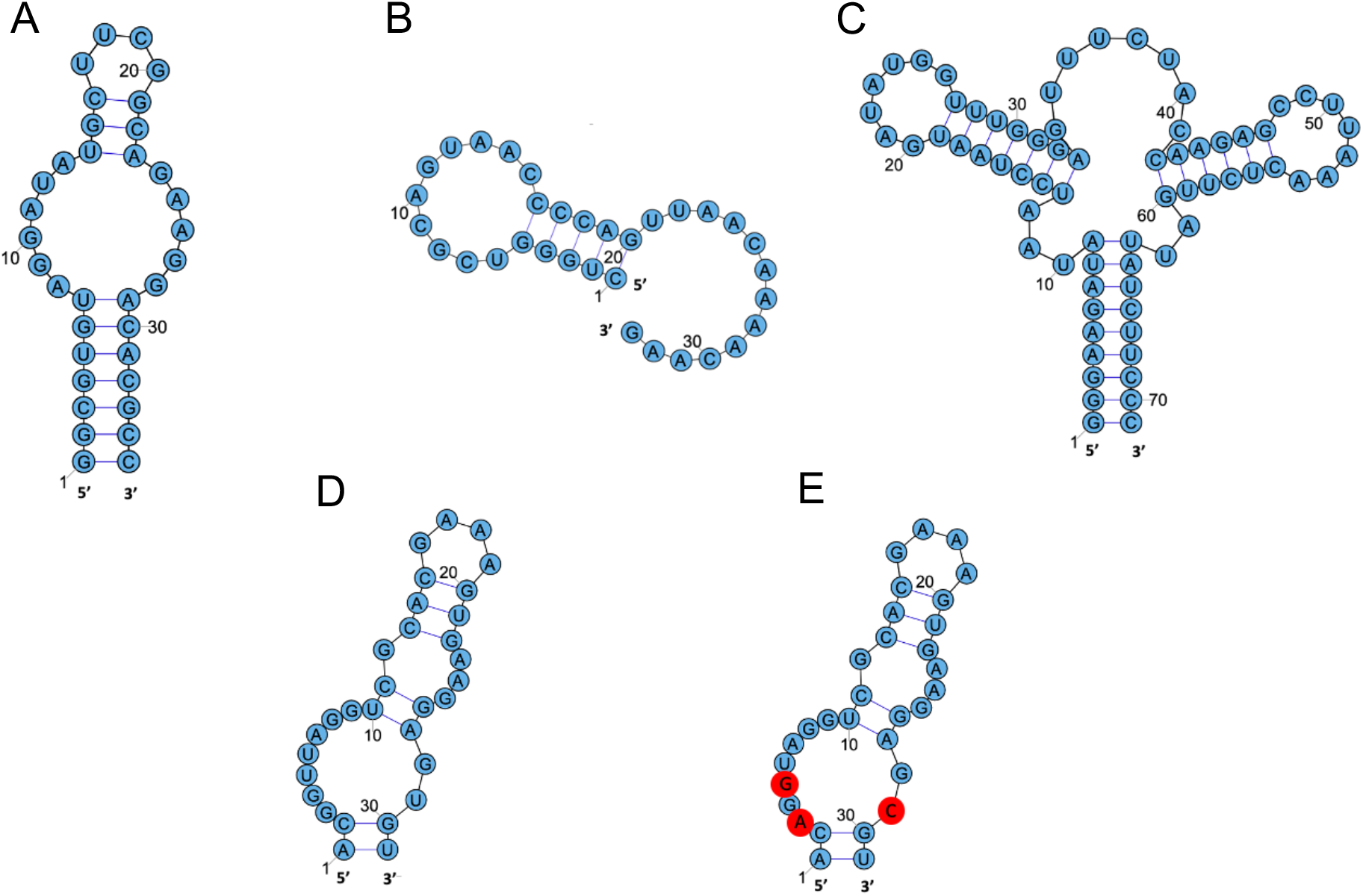
RNA secondary structure of **(A)** FMN aptamer; **(B)** preQ1 riboswitch aptamer; **(C)** adenine riboswitch aptamer; **(D)** citrulline aptamer and **(E)** arginine aptamer.

## MATERIAL AND METHODS

### System Setup

The starting coordinates of five small molecule ligand-RNA aptamer complex were derived from Protein Data Bank (PDB ID 1FMN, 3Q50, 5SWE, 1KOD and 1KOC). Five small molecule ligands including FMN, preQ1, adenine, citrulline and arginine were parameterized by Q-Chem and ACPYPE. Q-Chem 5.2 ^41^ was used to calculate optimized molecule geometry and RESP charges with method of Hartree-Fock and basis set of 6-31G(d) and cc-pVDZ (File S1). Topology files of small molecule ligands were produced by ACPYPE ^42^. The GROMACS 2019.6 software package ^28, 43–47^ compiled with PLUMED 2.7.0 ^40, 48–50^ was used for metadynamics simulations of five systems of small molecule ligand-RNA aptamer complex with water and ions in a box model. We chose the TIP4P explicit water model and revised forcefield for RNA by Chen and Garcia et al ^51^, which corrected the issue of hyper stacked structures of Amber-99 force field caused by the erroneously strong base-stacking propensity, the Steinbrecher and Case bioorganic phosphate parameters were used to prevent formation of spurious phosphate-base hydrogen bonds ^23^. For all the metadynamics simulations in this study, 150 mM excess KCl using the Cheatham-Young TIP4P-adapted ion parameters ^52^ were used, and 300 K was chosen as the simulation temperature to optimized chances of observing reversible RNA-ligand binding events ^32^. All of the simulations were performed in a cubic box in periodic boundary conditions applied to x, y and z directions. The distance between box edge and the complex of small molecule ligand-RNA aptamer was set to 1.0 nm.

All systems were minimized for 50000 steps maximum to remove unnatural contact ^53^. Subsequently, 1 ns of MD equilibration in NVT and NPT condition at 300 K and 1 bar were performed for each system. The long-range electrostatic interactions were calculated by the Particle Mesh Ewald ^54,55^ method. The cutoff for nonbonded interactions was set as 1.2 nm and the integration time step was set to 2 fs. The velocity-rescale algorithm ^56^ was employed for temperature coupling and the Parrinello-Rahman algorithm ^57^ for pressure coupling. After equilibration, well-tempered metadynamics simulations were carried out. VMD ^58^ and PyMOL (Schrödinger) were used for data analysis and visualization.

### Theory of Well-Tempered Metadynamics

We briefly review here the principles of well-tempered metadynamics method. A complete theoíy can be found in ^59, 60^. Metadynamics is biased by an external history-dependent potential built as a sum of Gaussians centered along the trajectory, which is constructed in the space of the predefined collective variables ^50^. The bias potential at time t is given by:

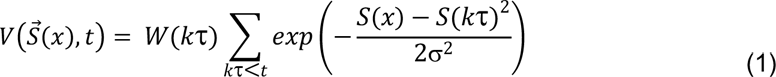

In eq (1), σ is the width of the Gaussians, W(kτ) is the height of the Gaussians, τ is the Gaussian deposition stride which means each Gaussian is deposited every τ MD timesteps and 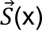 denotes the value of CVs ^61^. The bias potential of metadynamics can make the system escape local minimum to accelerate conformational transition against selected CVs. At long time, the value of bias potential converges to the projection of the free energy onto the CVs:

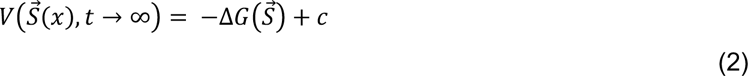

In direct metadynamics, Gaussian height was set to a constant value. As a result, the value of bias potential oscillates rather than converging ^62^. However, in well-tempered metadynamics, convergence of bias potential can be guaranteed by reducing Gaussian height in areas that are already strongly biased^40^:

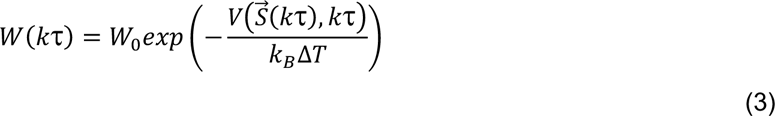

In eq (3), W_0_ is the initial Gaussian height, k_B_ is the Boltzmann constant, and ΔT is an additional hyperparameter with the dimension of temperature. With this rescaling of the Gaussian height, bias potential smoothly converges to a fraction of free energy:

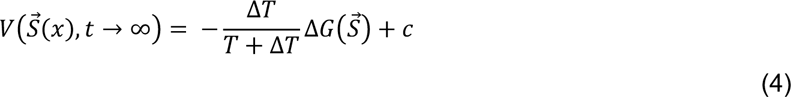

In eq (4), T is the temperature of system. (T+ΔT) is the effective temperature experienced along the CVs, which is higher than the system temperature T. Therefore, parameter of ΔT or “bias factor” 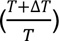 controls the extent of free energy exploration and strength of tempering: ΔT=0 corresponds to unbiased sampling. Conversely, ΔT→∞ corresponds to conventional metadynamics with constant Gaussian height.

In general, the bias of metadynamics constructed from the history of experienced configurations flattens the free energy landscape, which makes deep free energy basin regions accessible. Meanwhile, well-tempered metadynamics is able to limit the extent of exploration of the free energy landscape via bias factor and avoid irreversible sampling ^63^. In metadynamics, for mapping a free energy landscape from a molecular dynamics simulation, free energy is calculated from the sum of the Gaussians. Thus the energy of certain conformation with relevant value of CVs can be obtained, which provides a quantitative method for structural stability comparison. Moreover, since metadynamics can efficiently sample conformations of transition state along the trajectory, it is also possible to discover the reaction pathways of metastable states via metadynamics ^53^.

### Method Design and Workflow

#### Choice of Collective Variables

The selection of collective variables influences the effectiveness of metadynamics significantly. The ideal collective variable should be able to push the system to explore the high free energy states of your interests by taking different values in all relevant metastable states and the transition states between them, which are not accessible when the system is trapped in local minimum ^60, 63^.

In this study, we aim to simulate the whole induced-fit binding pathway of the recognition between small molecule ligand and its RNA aptamer binding pocket, and expect that the binding mode of small molecule ligand in the energy basin of free energy surface (FES) predicted by metadynamics has good agreement with the complexed structure solved by X-ray crystallography or NMR. In unbiased MD simulation, the binding/unbinding events of small molecule ligand-RNA system are not accessible in the timescale of hundreds of nanoseconds due to the high energy barriers in the binding/unbinding pathway. Therefore, to add extra potential for overcoming the energy barriers in the binding /unbinding pathway, the center-of-mass (COM) distance between small molecule ligand and its RNA binding pocket was selected as the first CVs (d of CVs). To enhance the motion and rotation of the groups of small molecule ligand within the RNA binding pocket and test on the importance of hydrogen bonds captured by X-ray crystallography or NMR, two of the hydrogen bond donor-to-acceptor distances was selected as the second and third CVs (d1, d2). Different value combinations of (d, d1, d2) correspond to different states in the binding /unbinding pathway or different binding modes of small molecule ligand within RNA binding pocket. The two-dimensional FES on (d1, d2) indicates whether the two hydrogen bonds represented by d1 and d2 are necessary for the energy basin binding modes. For the defining of binding pocket, we chose all of the nucleotides that interacted with small molecule ligand via hydrogen binding or base stacking captured by X-ray crystallography or NMR.

#### Optimization of metadynamics Parameters

The implementation of metadynamics simulation combining with restrains is widely applied to improve the efficiency of sampling ^64^. To prevent sampling from the unnecessary region out of the RNA aptamer binding pocket, the harmonic wall restrain, UPPER_WALLS, was used on d of CVs. The harmonic potential at value x of d is given by:

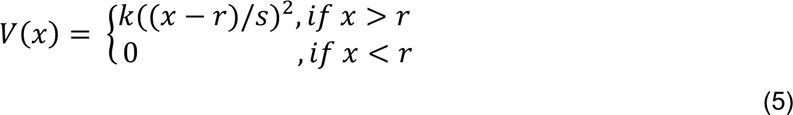

where k is the force constant, r is the upper wall threshold and s is a rescaling factor ^65^. The upper wall restrain placed on the COM distance between small molecule ligand and its RNA binding pocket can limit the range of motion of ligand to a spherical area with the COM of RNA binding pocket as the center. To prevent ligand from moving too far from the COM of RNA binding pocket and ensure effective interaction between ligand and RNA binding pocket along the trajectory of simulation, the radius r of the spherical region of upper wall restrain is determined by the size of RNA binding pocket depending on different ligand-RNA system. There is no bias of upper wall restrain within the spherical area, which only acts on d when ligand moves out of the spherical area representing the RNA binding pocket.

Since the fluctuation of CVs was limited by upper wall restrain, the width of Gaussians (σ) was determined by 1/3 of the radius of upper wall restrain. The value combination of initial Gaussian height (W_0_) and bias factor (γ) should ensure that ligand has sufficient potential to move and rotate frequently within the RNA binding pocket and the metadynamics simulation can converge in the end with the Gaussian height dropping to near zero. Direct metadynamics simulations with different value of Gaussian height in parallel were run to optimize W_0_, after which well-tempered metadynamics simulations with different value of bias factor in parallel were run to optimize γ. Gaussians were deposited every picosecond (500 steps). The optimized parameters of metadynamics for each system are summarized in (Table 1).

**Table 1.** Parameters for metadynamics simulations.

| System | Initial Gaussian height $W_0$ (kJ mol<sup>-1</sup>) | Bias factor $\gamma$ | Gaussian width $\sigma$ (nm) | Gaussian deposition stride $\tau$ (steps) | Simulation time $t$ (ns) | UPPER_WALLS restrain force constant $k$ (kJ mol<sup>-1</sup> nm<sup>-2</sup>) | UPPER_WALLS restrain threshold $r$ (nm) |
| --- | --- | --- | --- | --- | --- | --- | --- |
| 1FMN | 0.3 | 4 | 0.27 | 500 | 30 | 5000 | 0.8 |
| 3Q50 | 0.3 | 4 | 0.27 | 500 | 40 | 5000 | 0.8 |
| 5SWE | 0.8 | 4 | 0.27 | 500 | 20 | 5000 | 0.8 |
| 1KOD | 0.6 | 100 | 0.20 | 500 | 100 | 5000 | 0.6 |
| 1KOC | 0.6 | 6 | 0.23 | 500 | 60 | 5000 | 0.7 |

To evaluate the convergence of metadynamics predictions and establish unbiased free energy for reference, all of the simulations were reweighted for removing the effect of metadynamics bias on the free energy surface. The unbiased distribution of each CV was recovered using the reweighting algorithm introduced by Branduardi et al. ^66^

## RESULTS

This study simulated the molecular recognition process of five systems of small molecule ligand-RNA binding pocket complex via metadynamics. The binding modes at energy basin predicted by metadynamics simulation have good agreements with the solved structures in the Protein Data Bank. They are, NMR solution structure of the complex of flavin mononucleotide bound to a 35-nucleotide RNA aptamer identified through in vitro selection, crystal structure of preQ1 riboswitch aptamer in the metabolite-bound state, crystal structure of ligand-bound adenine riboswitch aptamer domain, NMR solution structure of RNA aptamer complexed with citrulline and NMR solution structure of RNA aptamer complexed with arginine.

### FMN/RNA Aptamer

The first small molecule ligand-RNA binding pocket system studied is the FMN/RNA aptamer complex. In the solved NMR solution structure in Protein Data Bank (PDB ID: 1FMN), the uracil like edge of isoalloxazine ring of FMN formed an A·U Reverse Hoogsteen pairing with the Hoogsteen edge of A26 by the two hydrogen bonds of A26:N6-H…FMN:O2 and FMN:N3-H…A26:N7. The isoalloxazine ring of FMN base stacked in between G·U·A triplet and G·G mismatch above and below (Figure 5B). Therefore, to simulate the whole binding process of FMN via metadynamics, we chose the center-of-mass distance between FMN and binding pocket as the first collective variables (d of CVs). The initial structure of simulation is the NMR solution structure with FMN moved 3.3 nm away from the center-of-mass of the binding pocket. The binding pocket is defined by the G10·U12·A25 triplet and G9·G27 mismatch above and below FMN binding nucleotide, A26. To test on the importance of these two hydrogen bonds for binding, we chose the hydrogen bond donor-to-acceptor distances of these two hydrogen bonds as the second and third CVs (d1 and d2).

In the simulation, FMN was pushed back into the binding pocket by upper wall restrain instantly, as shown in (Figure 3A) and (Movie S1), the center-of-mass distance between FMN and binding pocket dropped to 0.8 nm from 3.3 nm within the first couple frames. Afterwards the range of motion of FMN was limited by the upper wall restrain to a spherical area with the COM of binding pocket as the center and a radius of 0.8 nm, and FMN started trying various binding poses in this range. Under the effect of Gaussian potential of 0.3 kJ/mol, FMN obtained adequate energy for overcoming the energy barriers of state transition so that achieved movement and rotation as much as possible, which means the achievement of different binding modes of FMN in the RNA binding pocket as much as possible. Under an optimized value of bias factor of 4.0, height of Gaussian decreased whenever FMN ligand reaches energy favorable binding modes to keep FMN staying in this state. On the contrary, when FMN met an energy unfavorable pose, Gaussian height will be strengthened to force FMN leaving this state so that it will continue looking for more energy favorable binding modes. After repeating this process, when the most energy favorable binding mode was found, Gaussian height dropped to near zero without rebounding, as shown in (Figure 3B), which is a sign of the convergence of the metadynamics simulation. Compared with unbiased molecular dynamics simulation, the movement intensity of FMN ligand is significantly increased. The extra energy is from the Gaussian potential being added on the three collective variables (CVs), the COM distance between FMN and binding pocket (d of CVs), the distance between hydrogen bond donor A26:N6 and hydrogen bond acceptor FMN:O2 (d1 of CVs), the distance between hydrogen bond donor FMN:N3 and hydrogen bond acceptor A26:N7 (d2 of CVs). Gaussian potential being added on these three CVs forced their value constantly changing so that FMN ligand can realize the high-intensity motion which is unrealizable in unbiased MD simulation. Therefore, each value combination of these three CVs represents a specific binding mode. As shown in the final result of free energy surface calculation in (Figure 4), the deepest energy basin is located at d=0.14 nm, d1=0.29 nm and d2=0.30 nm in both biased and unbiased (reweighted) FES. The value of d1 lower than or equal to 0.35 nm means the hydrogen bond formation between FMN:O2 and A26:N6 and the value of d2 lower than or equal to 0.35 nm means the hydrogen bond formation between FMN:N3 and A26:N7. Thus, this energy basin conformation referred to the binding mode of FMN in the very end of this 30 ns-long metadynamics simulation when the two hydrogen bonds of d1 and d2 formed at the same time (Figure 5A). As shown in (Figure 5), the binding mode of FMN at the deepest energy basin of simulation is very close to the NMR solution structure (RMSD = 1.98 Å) with the same two hydrogen bonds between the uracil like edge of the isoalloxazine ring of FMN and Hoogsteen edge of A26, G10·U12·A25 triplet and G9·G27 mismatch above and below the binding site.

Moreover, the second deepest energy basin was located at d1=1.16 nm, d2=0.49 nm and d=0.44 nm, which referred to the binding mode as shown in (Figure 6). In this binding mode, instead of the two hydrogen bonds of d1 and d2, there are two other essential hydrogen bonds captured, G10:N2-H…FMN:O4, G10:N2-H…FMN:N5. The third deepest energy basin was located at d1=1.56 nm, d2=1.49 nm and d=0.77 nm, which referred to the binding mode with the isoalloxazine ring of FMN base stacking between G10·U12·A25 triplet and G27 (Figure 7). Therefore, we can see that this ‘docking-like’ metadynamics simulation strategy can accurately predict the most stable binding mode of FMN/RNA aptamer binding pocket system, meanwhile, provide other possible existing binding poses that are energy favorable via different essential interactions with the same RNA binding pocket. The metadynamics simulation also provided an potential dynamic pathway of binding for FMN/RNA aptamer complex. As shown in (Movie S1), before the isoalloxazine ring of FMN intercalated between G10·U12·A25 triplet and G9·G27 mismatch, A26 formed A26·G9 mismatch with G9, G10 partly stacked in between U12·A25 Hoogsteen base-pairing and A26·G9 mismatch, and G27 in bulge state. The intercalating of FMN pushed G10 to G10·U12·A25 triplet state. Afterwards, A26 recognized the uracil like edge of FMN, G27 moved upwards with A26 and formed G27·G9 mismatch with G9.

**Figure 3.**
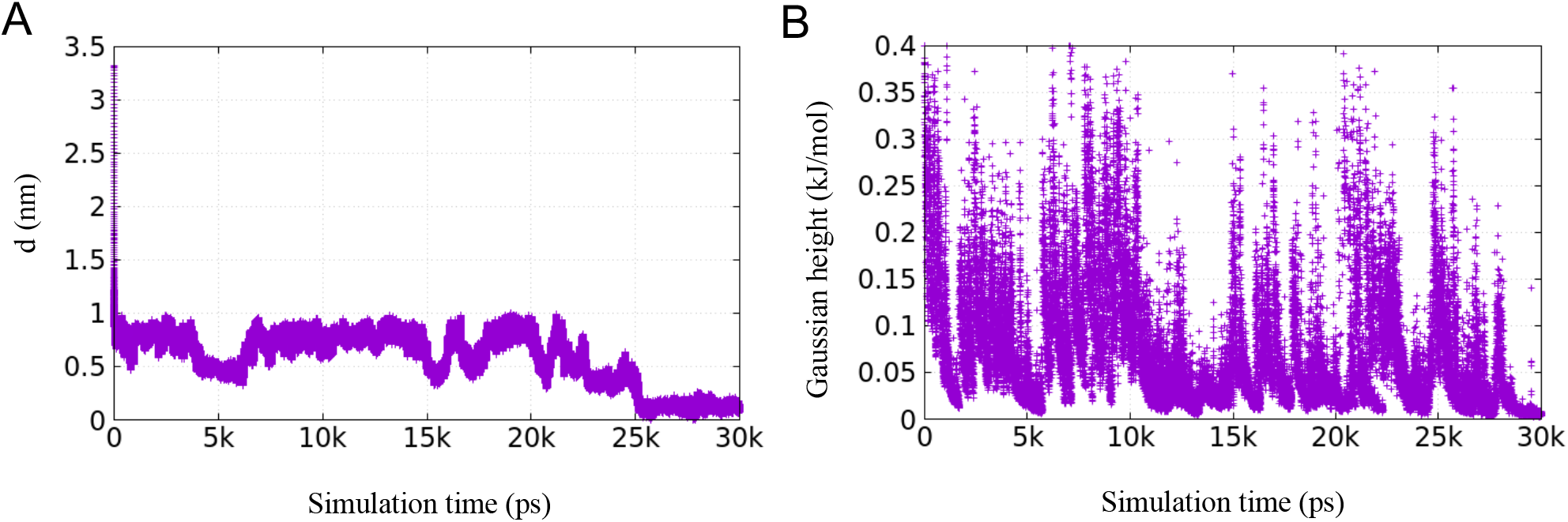
Time evolution of **A)** d of CVs (COM distance between FMN and RNA binding pocket defined by G10·U12·A25 triplet and G9·G27 mismatch above and below FMN binding site) and **B)** Gaussian height in a 30ns-long well-tempered docking metadynamics simulation of FMN/RNA aptamer system.

**Figure 4.**
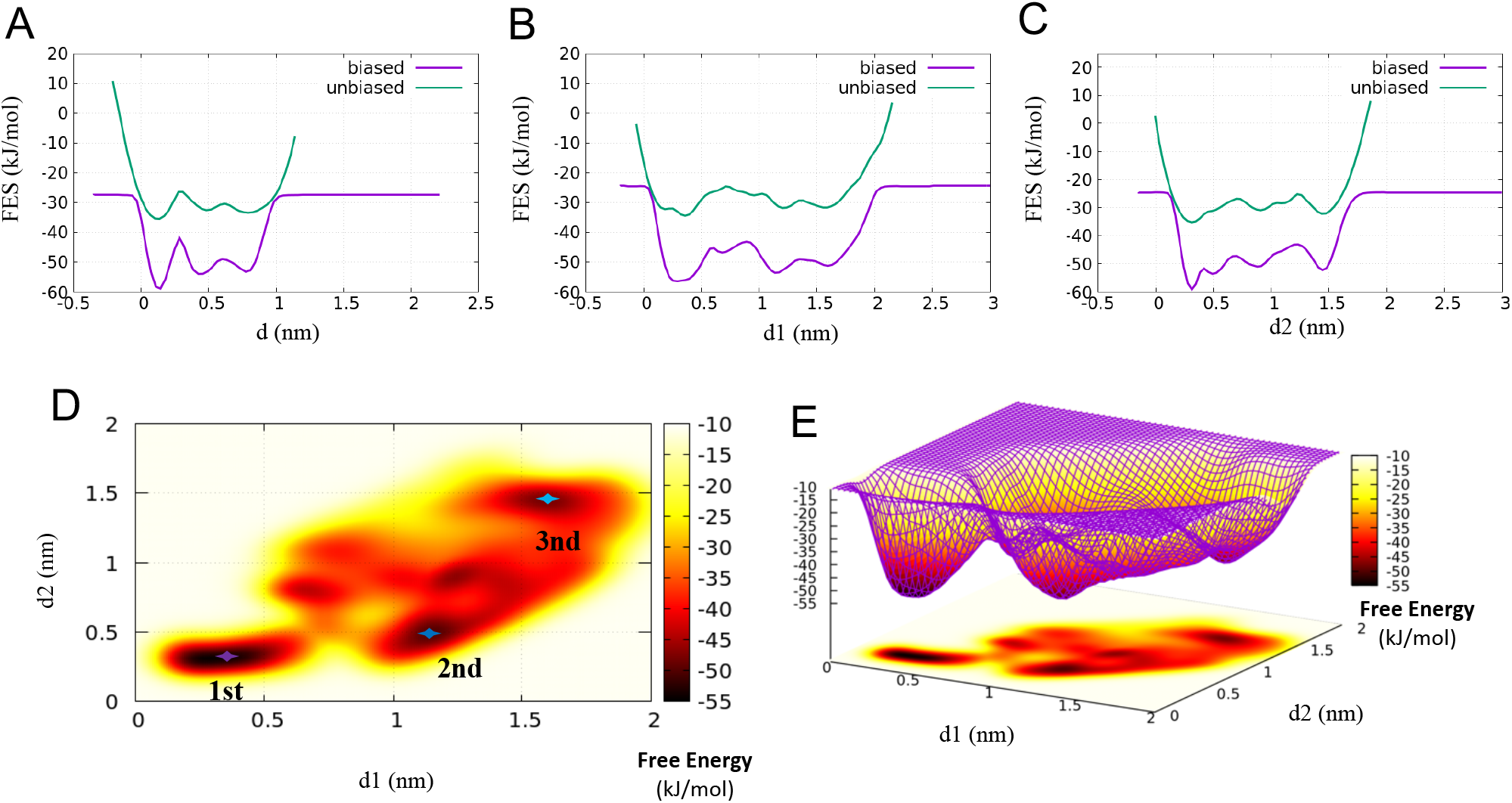
Comparison between the binding FES calculated from the metadynamics bias potential (biased, in purple) and by reweighting (unbiased, in green) as a function of **A)** d of CVs (COM distance between FMN and binding pocket); **B)** d1 of CVs (the distance between FMN:O2 and A26:N6); **C)** d2 of CVs (the distance between FMN:N3 and A26:N7). Two-dimensional free energy landscape projected onto d1 and d2 **D)** presented in 2D; **E)** presented in 3D

**Figure 5.**
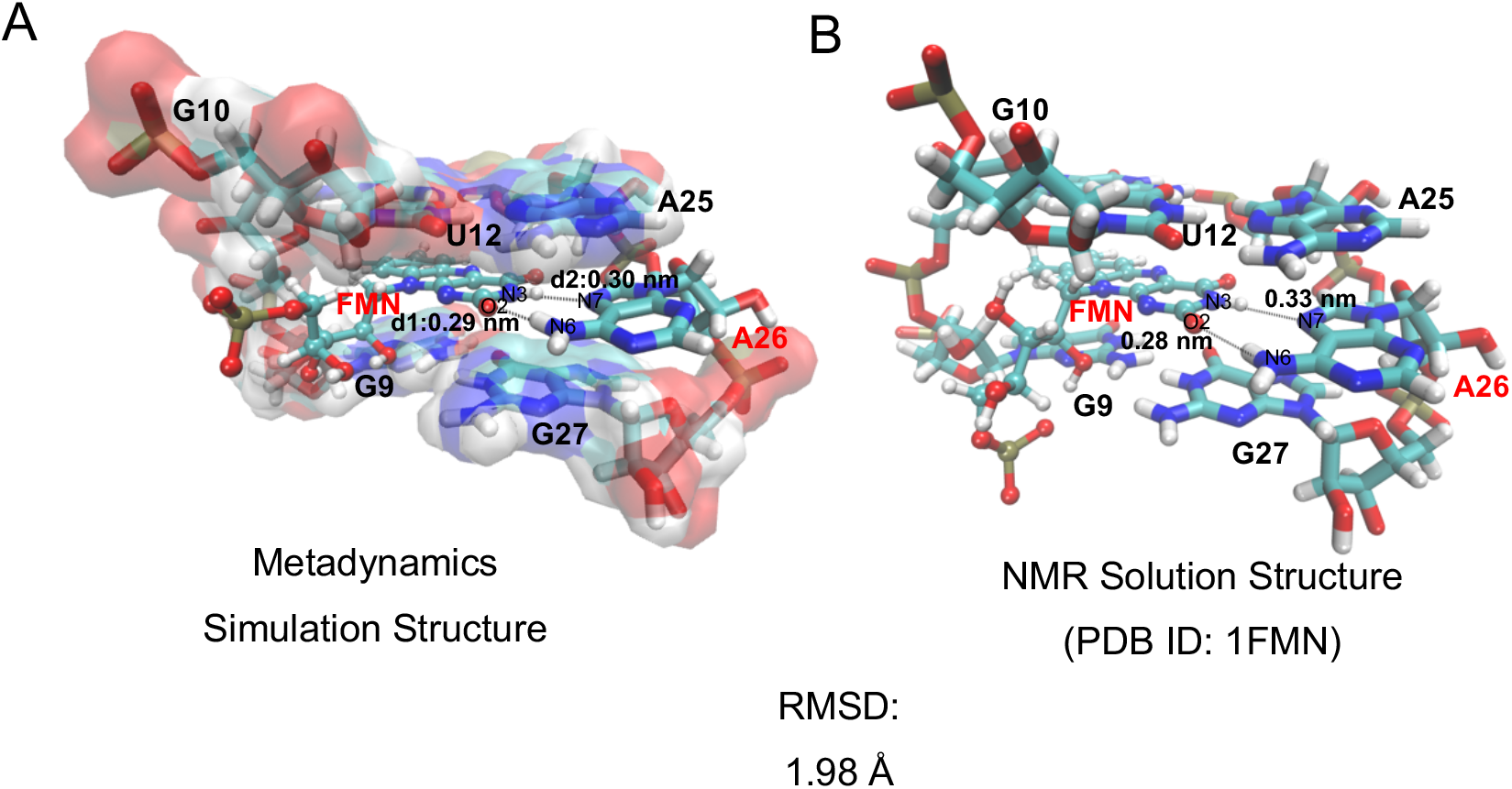
**A)** Snapshot of binding mode at energy basin of 2D free energy landscape onto d1 and d2 from metadynamics simulation **B)** NMR solution structure of FMN-FMN aptamer complex (PDB ID: 1FMN)

**Figure 6.**
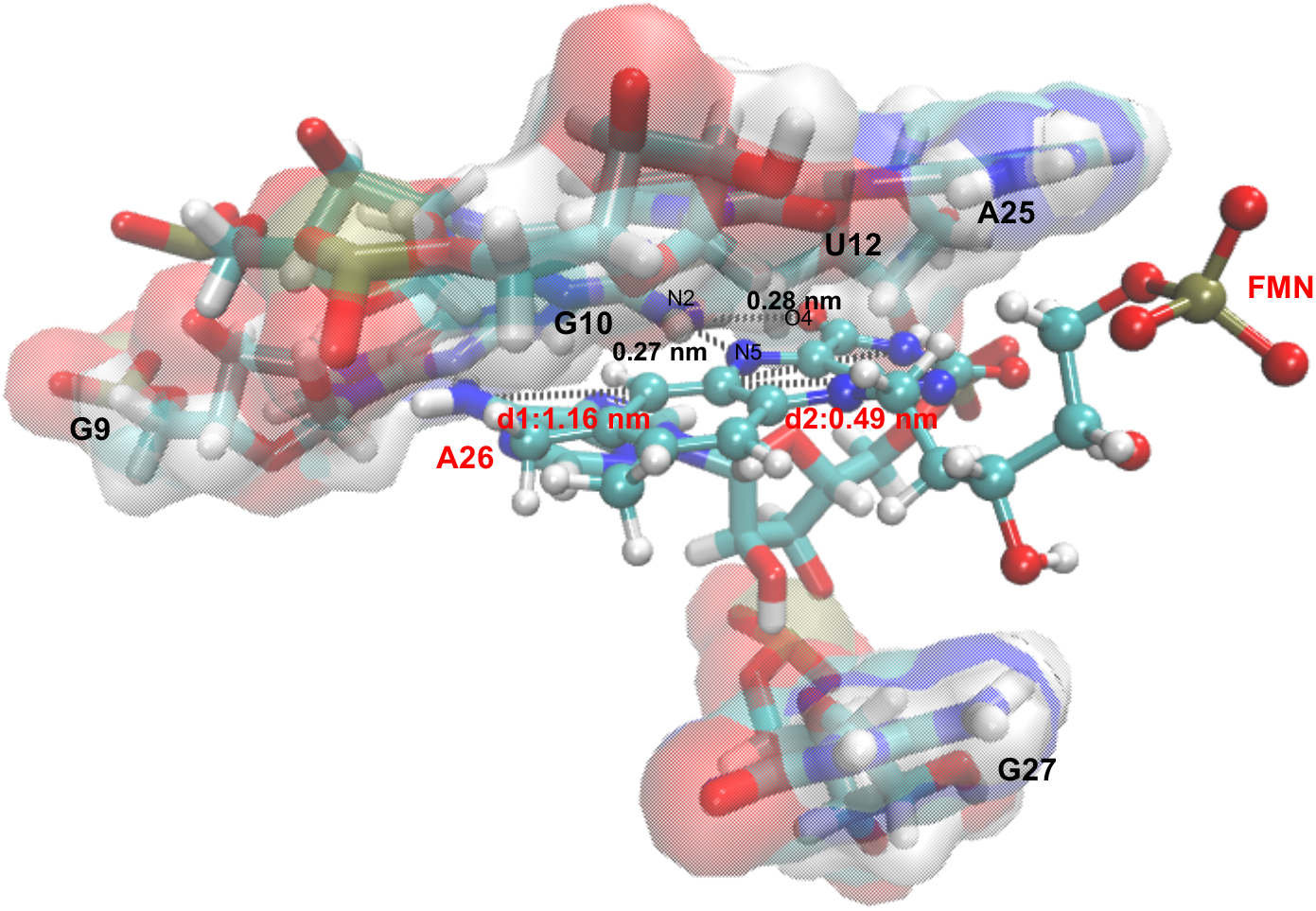
Snapshot of binding mode at the second energy basin of 2D free energy landscape onto d1 and d2 from metadynamics simulation.

**Figure 7.**
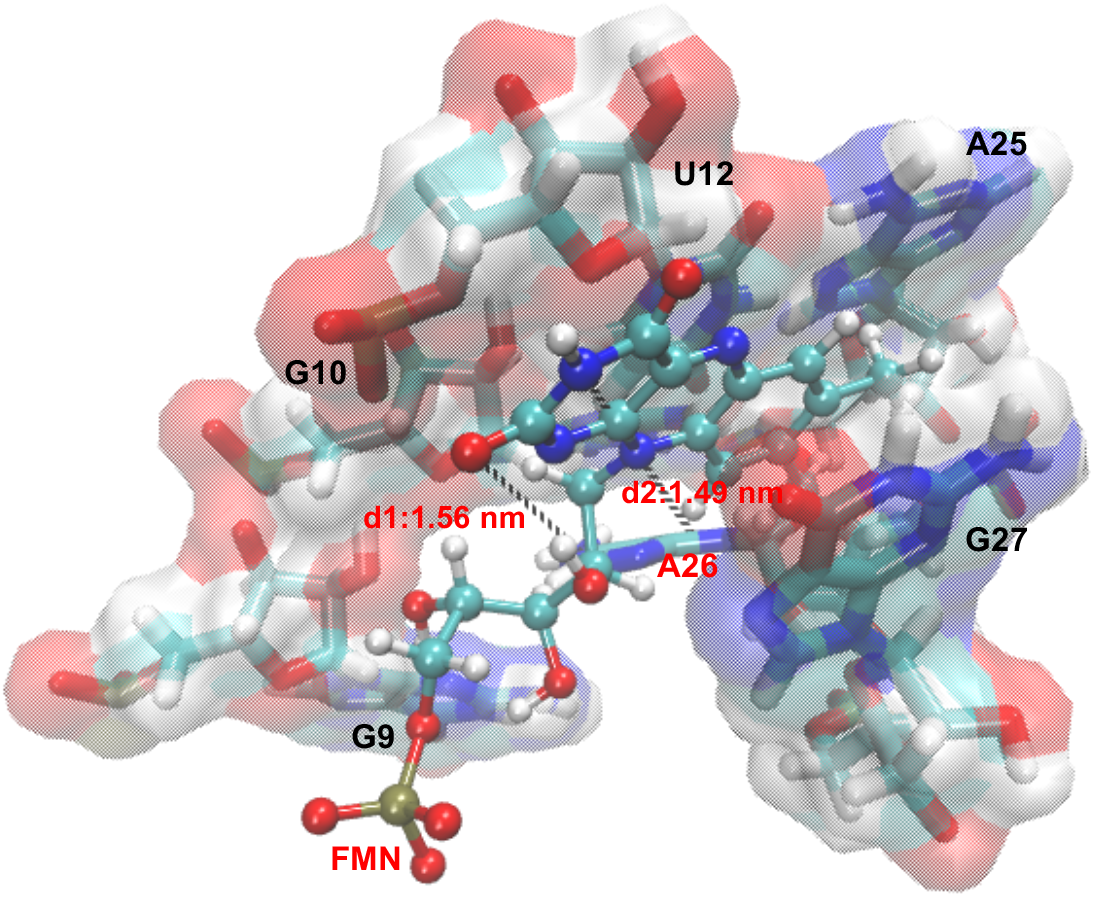
Snapshot of binding mode at the third energy basin of 2D free energy landscape onto d1 and d2 from metadynamics simulation.

### PreQ1 Riboswitch Aptamer

To test if this metadynamics simulation method is able to reproduce accurate binding mode prediction of small molecule ligand/RNA binding pocket complex, we chose to dock ligand of preQ1 on its riboswitch aptamer. In the solved crystal structure of preQ1 riboswitch aptamer in metabolite bound state (PDB ID: 3Q50), the Watson-Crick face of preQ1 formed G·C-like pairing with C15 by three hydrogen bonds of C15:N4-H…preQ1:O6, preQ1:N1-H…C15:N3 and preQ1:N2-H…C15:O2. The binding site of preQ1·C15 was stacked between G5·C16 base pair and A14·G11·C7 triplet, which have been referred to as the “floor” and “ceiling” of the binding pocket (Figure 10B). Based on the same strategy of metadynamics we applied on FMN/RNA aptamer system, the COM distance between preQ1 ligand and riboswitch binding pocket was chosen as the first CVs (d of CVs), the distance from C15:N4 to preQ1:O6 was chosen as the second CVs (d1 of CVs) to represent the first core hydrogen bond of preQ1·C15 pairing, the distance from preQ1:N1 to C15:N3 was chosen as the third CVs to represent the second core hydrogen bond of preQ1·C15 pairing. Binding pocket is defined by A14·G11·C7 triplet and G5·C16 base pair above and below the binding site.

As shown in (Figure 8A) and (Movie S2), preQ1 started from 1.8 nm away from the center-of-mass of binding pocket, and the guanine base of preQ1 intercalated between G5·C16 base pair and G11·C7 mismatch mainly driven by upper wall restrain on d of CVs with 0.8 nm as radius of sphere. Afterwards the minor groove of preQ1 was recognized by bases of U6 and A29 firstly, and preQ1 rotated in the plane of preQ1·C15 for adjusting its angle towards C15. Meanwhile, C15 kept swinging in and out of the major groove, switching between C15 bulge state and base stacking state until preQ1 ‘read out’ the Watson-Crick edge of C15 at the 8th ns. The preQ1·C15 pairing stabilized the ‘celling’ of A14·G11·C7 triplet via base stacking force, and stayed in this state for the rest 32 ns steady without obvious changing in its center-of-mass location (Figure 8A). This dynamic process can explain why Gaussian height pumped up and down between 0.3 and 0 kJ/mol in the first 8 ns and dropped to near zero afterwards without rebounding in the last 32 ns (Figure 8B). PreQ1 was powered by gaussian potential constructed on the three CVs to find an energy favorable binding mode within the binding pocket in the first 8 ns. When preQ1 formed G·C-like pairing with C15, gaussian height decreased to near zero to keep preQ1 staying in this energy favorable state. The free energy surface of metadynamics simulation shows an single energy basin at d1= 0.29 nm, d2= 0.29 nm and d= 0.10 nm from both biased and reweighted calculation (Figure 9). The binding mode at energy basin located by d1= 0.29 nm, d2= 0.29 nm has an RMSD of 0.87Å from the crystal structure with core hydrogen bonds of d1 and d2 for preQ1·C15 pairing, A14·G11·C7 as ceiling of binding pocket and C16·G5 as binding pocket floor (Figure 10). The FES result once again proved that this metadynamics simulation strategy can predict binding modes of small molecule ligand/RNA binding pocket complex with high accuracy. Moreover, the crystal structure of preQ1 riboswitch aptamer in metabolite free states shows that C15 swinging out of major groove and A14 shifting into the preQ1 binding site result in the ‘close’ access to preQ1 ligand. Our metadynamics simulation also captured the swinging of C15 during preQ1 recognition, which shows the potential of metadynamics simulation for describing dynamic mechanism of molecular recognition.

**Figure 8.**
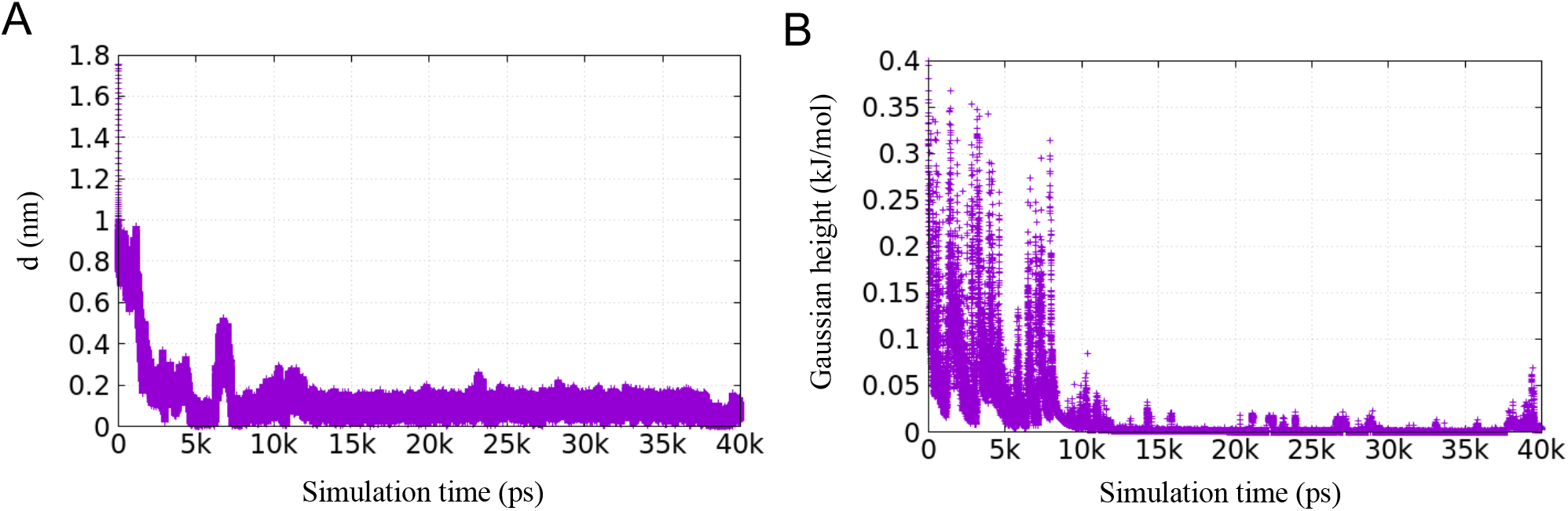
Time evolution of **A)** d of CVs (COM distance between preQ1 and RNA binding pocket defined by A14·G11·C7 triplet and C16·G5 base pair above and below preQ1 binding site) and **B)** Gaussian height in a 40 ns-long well-tempered docking metadynamics simulation of preQ1 riboswitch aptamer system.

**Figure 9.**
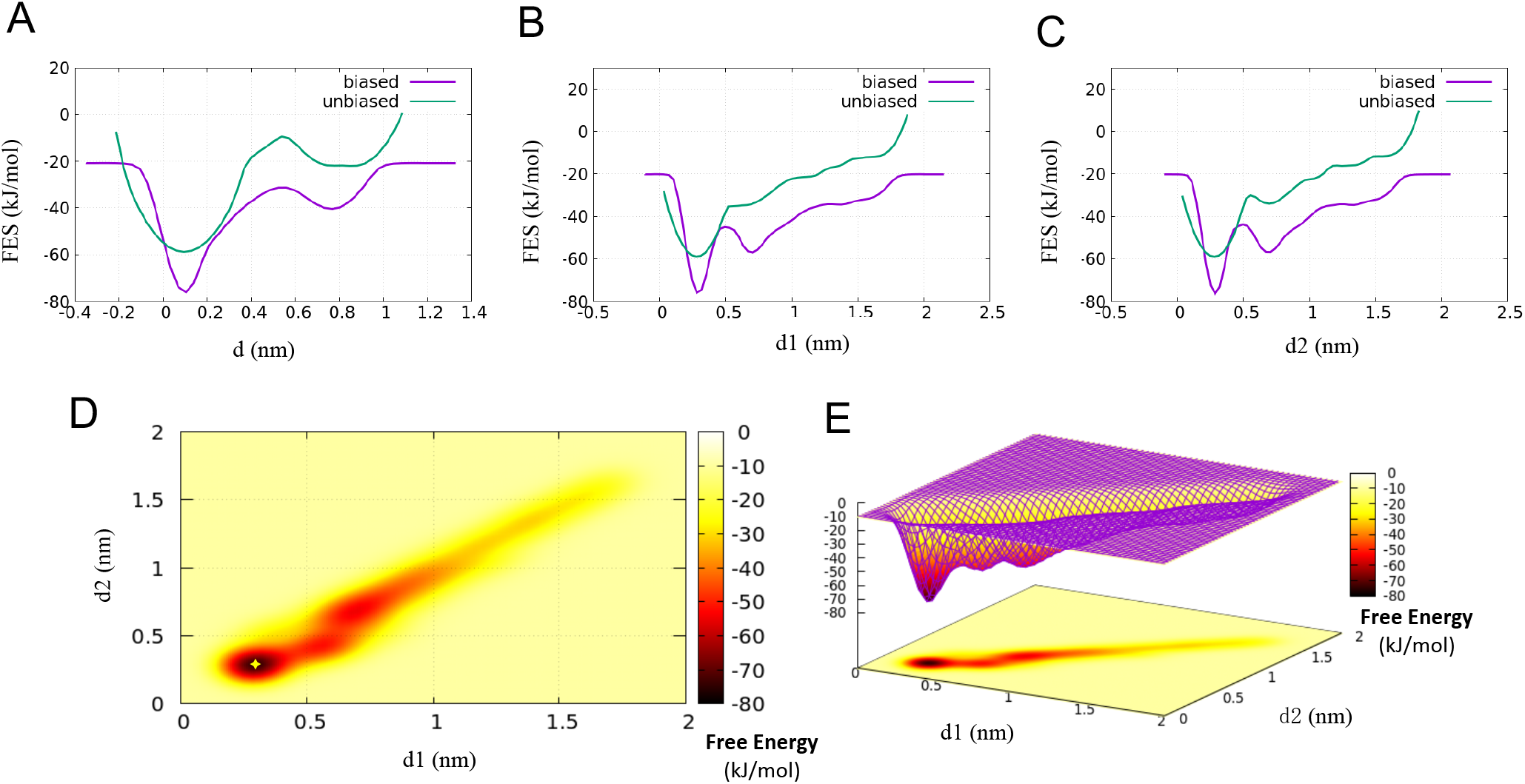
Comparison between the binding FES calculated from the metadynamics bias potential (biased, in purple) and by reweighting (unbiased, in green) as a function of **A)** d of CVs (COM distance between preQ1 and binding pocket); **B)** d1 of CVs (the distance between preQ1:O6 and C15:N4); **C)** d2 of CVs (the distance between preQ1:N1 and C15:N3). Two-dimensional free energy landscape projected onto d1 and d2 **D)** presented in 2D; **E)** presented in 3D

**Figure 10.**
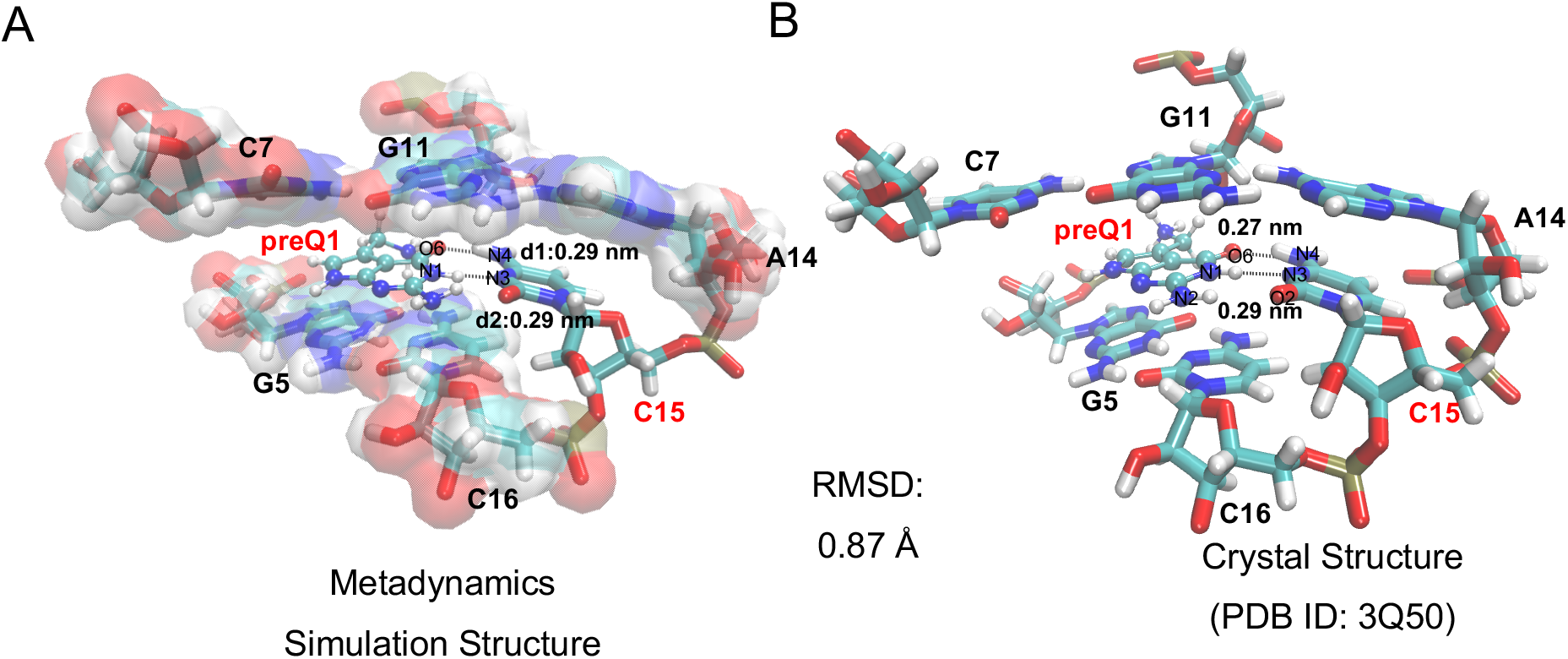
**A)** Snapshot of binding mode at energy basin of 2D free energy landscape onto d1 and d2 from metadynamics simulation; **B)** Crystal structure of preQ1 riboswitch aptamer complex (PDB ID: 3Q50)

### Adenine Riboswitch Aptamer Domain

Another case of riboswitch studied was adenine riboswitch aptamer domain, which is a 71-nucleotide RNA (rA71) of the translational control riboswitch from Vibrio vulnificus. The crystallography study showed that, in the final ligand-bound conformation, U74 as the ligand-recognition residue formed Watson-Crick base pairing with adenine ligand via two hydrogen bonds of ADE:N6-H…U74:O4 and U74:N3-H…ADE:N1. The binding site of ADE·U74 was base stacked between two base-triples (Figure 13B) (PDB ID:5SWE). Therefore, the binding pocket of our metadynamics simulation was defined by these two triplets of C50·U75·A21 and A73·A52·U22 above and below the binding site. Based on the same collective variable setting strategy, there are still three CVs, the COM distance between adenine ligand and the binding pocket (d of CVs), the distance between hydrogen bond donor of ADE:N6 and hydrogen bond acceptor of U74:O4 (d1 of CVs), the distance between hydrogen bond donor of U74:N3 and hydrogen bond acceptor of ADE:N1 (d2 of CVs).

The radius of the sphere of upper wall restrain acted on d of CV is 0.8 nm, which pushed adenine back into binding pocket from 1.9 nm away (Figure 11A). Due to the planar shape of adenine ligand, it intercalated between C50·U75·A21 and A73·A52·U22 easily. Afterwards, within the range of d less than 0.8 nm, adenine experienced different position relative to U74 which was powered by Gaussian potential on the three CVs. Under the effect of Gaussian height of 0.8 kJ/mol with bias factor of 4, adenine rotated and moved in the plane of adenine·U74 until forming A·U-like pairing with U74 in the 10th ns (Movie S3). After that, Gaussian height decreased to near zero (Figure 11B). The recognition between adenine ligand and U74 stabilized the whole binding pocket and was read as the most energy favorable conformation in the free energy surface. As shown in (Figure 12), the main energy basin is located at d1= 0.29 nm, d2= 0.30 nm and d= 0.23 nm in both biased and unbiased (reweighted) FES, which points to the state of adenine·U74 pairing as shown in (Figure 13A). This binding mode of adenine predicted by metadynamics has an RMSD of 0.87Å from the crystal structure (Figure 13B), which proved the accuracy of the metadynamics method again.

**Figure 11.**
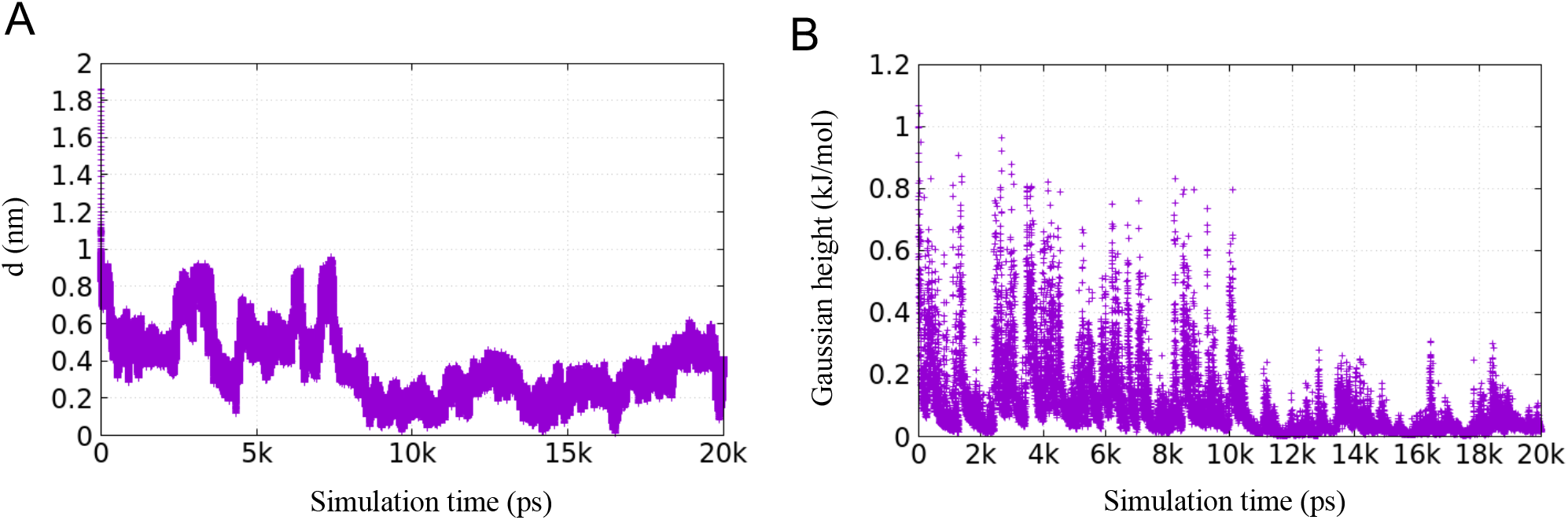
Time evolution of **A)** d of CVs (COM distance between adenine and RNA binding pocket defined by C50·U75·A21 and A73·A52·U22 triplet above and below adenine binding site) and **B)** Gaussian height in a 20ns-long well-tempered docking metadynamics simulation of adenine riboswitch aptamer domain system.

**Figure 12.**
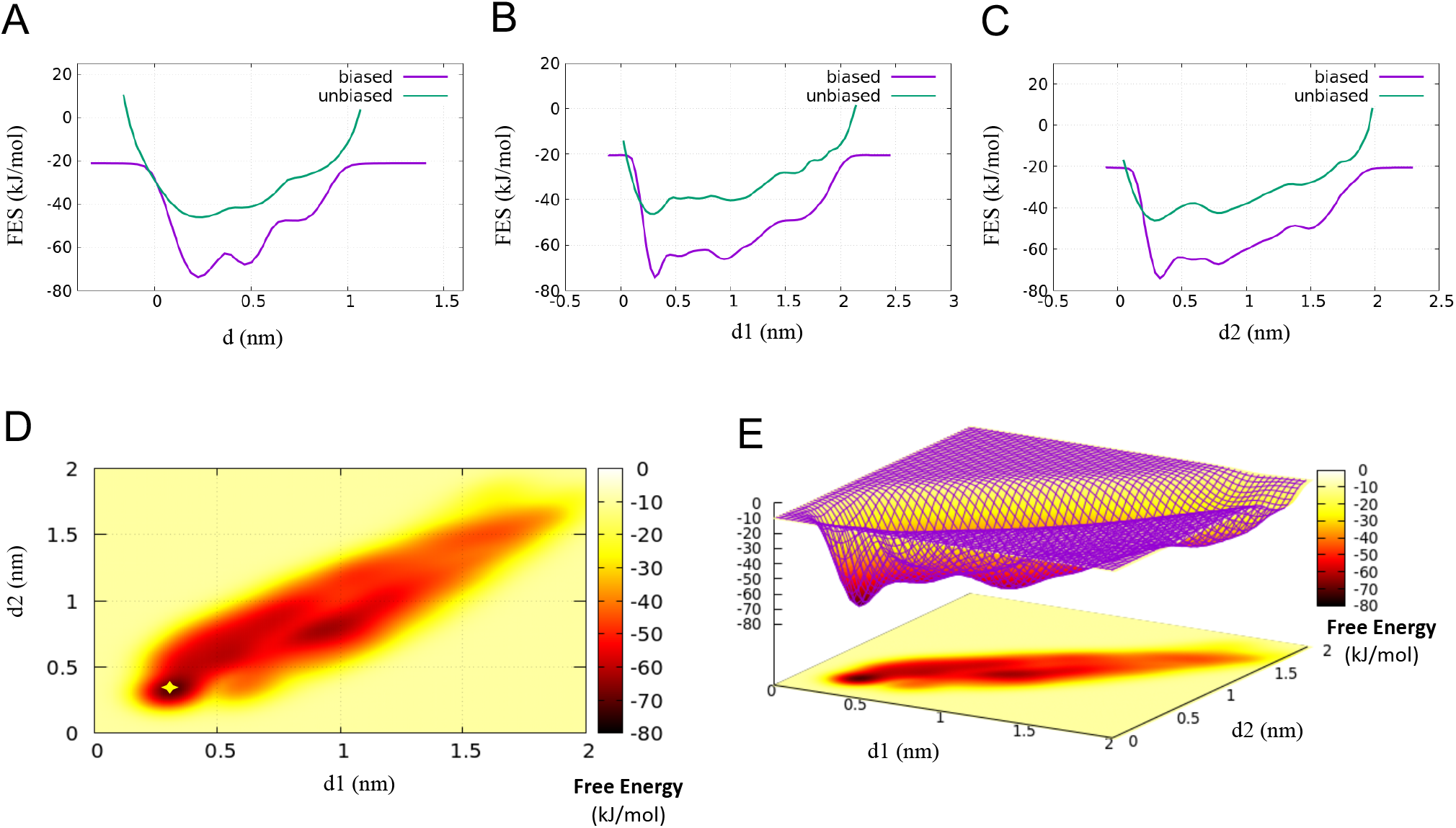
Comparison between the binding FES calculated from the metadynamics bias potential (biased, in purple) and by reweighting (unbiased, in green) as a function of **A)** d of CVs (COM distance between adenine and binding pocket); **B)** d1 of CVs (the distance between ADE:N6 and U74:O4); **C)** d2 of CVs (the distance between ADE:N1 and U74:N3). Two-dimensional free energy landscape projected onto d1 and d2 **D)** presented in 2D; **E)** presented in 3D

**Figure 13.**
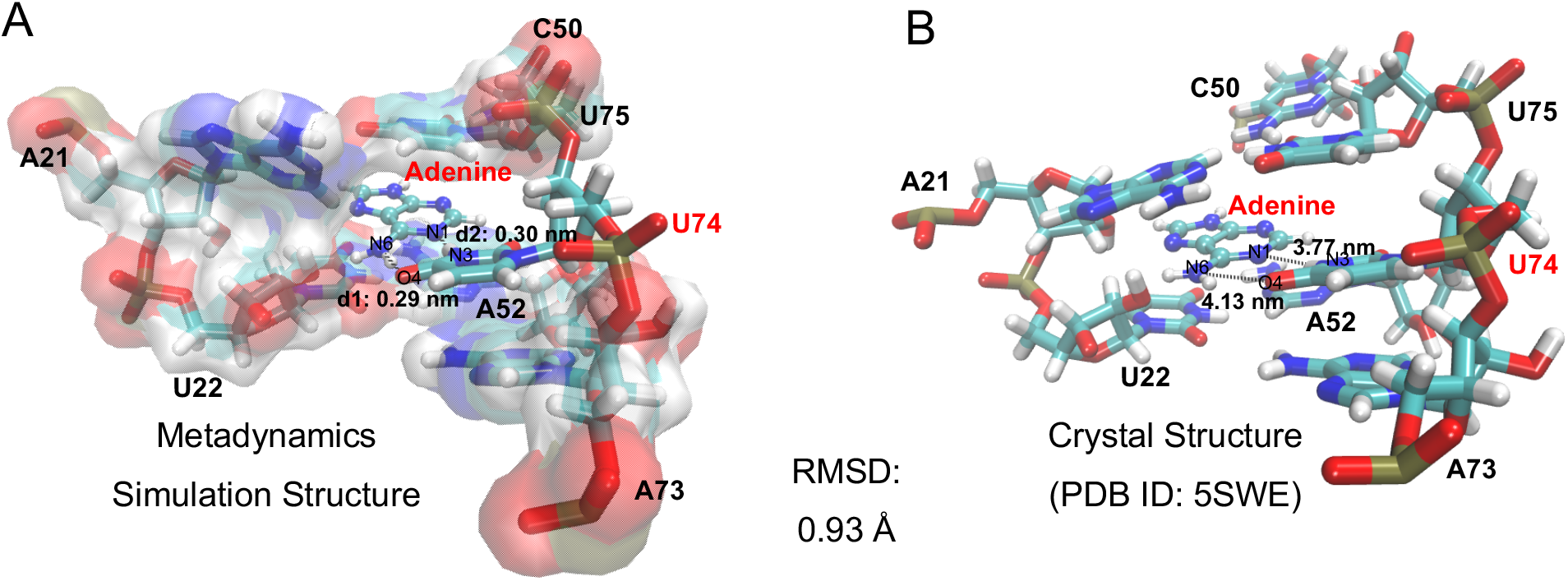
**A)** Snapshot of binding mode at energy basin of 2D free energy landscape onto d1 and d2 from metadynamics simulation; **B)** Crystal structure of adenine riboswitch aptamer domain complex (PDB ID: 5SWE)

### Citrulline/RNA Aptamer

The forth small molecule ligand-RNA system studied is citrulline/RNA aptamer complex obtained from in vitro selection. The initial structure of metadynamics is from NMR solution structure (PDB ID:1KOD). The binding pocket of metadynamics is defined same as the previous NMR spectroscopy study by G3-G4-U5-A7-G9-G25-G28-U29, G9-G28-G25 of which form an uneven plane of bottom, and G4-U5 nearly perpendicular to that surface (Figure 16B). The solved NMR solution structure showed that sequence-specific interactions occurred between the urea group of citrulline ligand and Watson-Crick edge of U29, Hoogsteen edge of G4 (Figure 16B). Therefore, we picked two core hydrogen bond distances of U29:N3-H…CIR:O7 and CIR:N8-H…G4:N7 as d1 of CVs and d2 of CVs respectively. d of CVs is still the COM distance between citrulline ligand and RNA binding pocket which functioned as the main driven force of binding/unbinding.

As shown in (Figure 14A) and (Movie S4), within the range of d less than 0.9 nm limited by upper wall restrain, citrulline experienced plenty of binding modes inside the RNA binding pocket in a 100 ns-long well-tempered docking metadynamics simulation. For this case, after optimization of bias factor, we found that under Gaussian height of 0.6 kJ/mol, a higher value of bias factor of 100 was needed to achieve the convergence of metadynamics simulation, which means 100 times of system temperature is needed to cross the relevant free-energy barriers efficiently in this simulation. Under the action of Gaussian height of 0.6 kJ/mol with bias factor of 100, citrulline realized relatively high-frequency state transitions, which effectively pushed citrulline away from energy-unfavorable binding modes until the energy basin conformation was met (Figure 14B). In this case, a relatively wide energy basin was found in the free energy surface. The most energy favorable binding mode of citrulline is located at d1= 0.26 nm, d2 = 0.27 nm and d= 0.26 nm in both biased and reweighted FES (Figure 15), which proved the importance of hydrogen bonding interactions between the urea group of citrulline and Watson-Crick edge of U29, Hoogsteen edge of G4 for the recognition between citrulline and its RNA binding pocket. As shown in (Figure 16), the binding mode of citrulline at energy basin of metadynamics has an RMSD of 0.86 Å from the NMR solution structure which is another successful case of small molecule-RNA binding mode prediction by our metadynamics Method.

**Figure 14.**
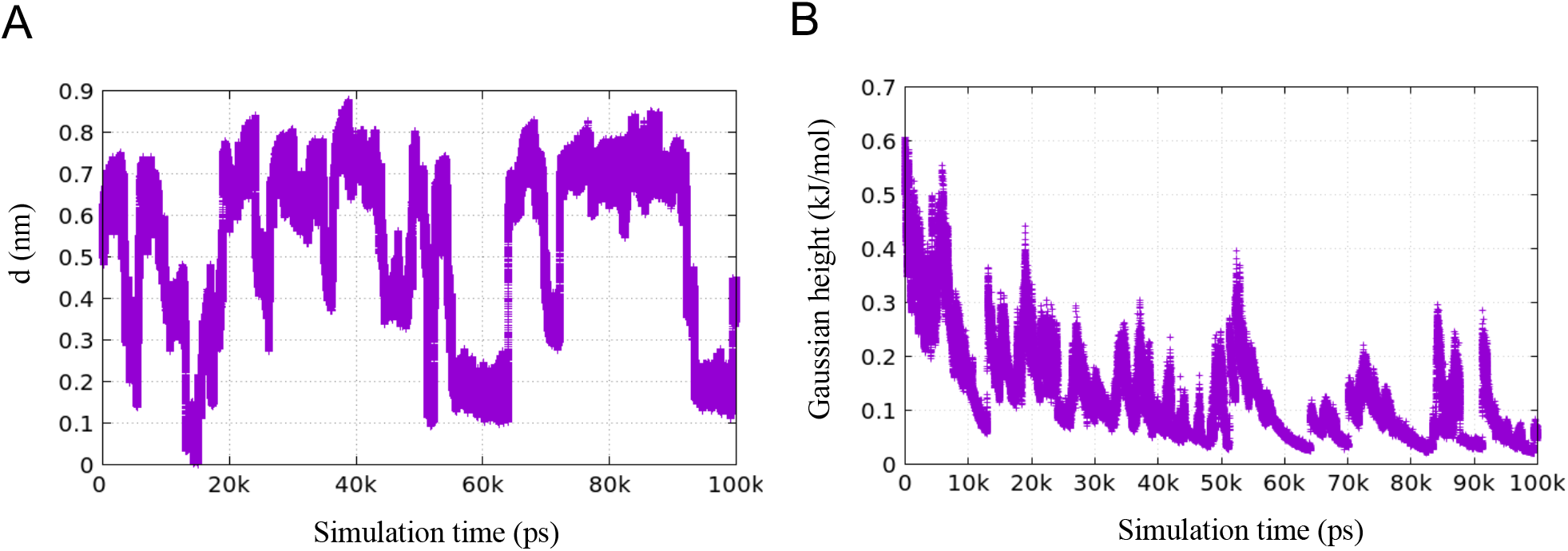
Time evolution of **A)** d of CVs (COM distance between citrulline and RNA binding pocket defined by G3-G4-U5-A7-G9-G25-G28-U29) and **B)** Gaussian height in a 100ns-long well-tempered docking metadynamics simulation of Citrulline RNA aptamer system.

**Figure 15.**
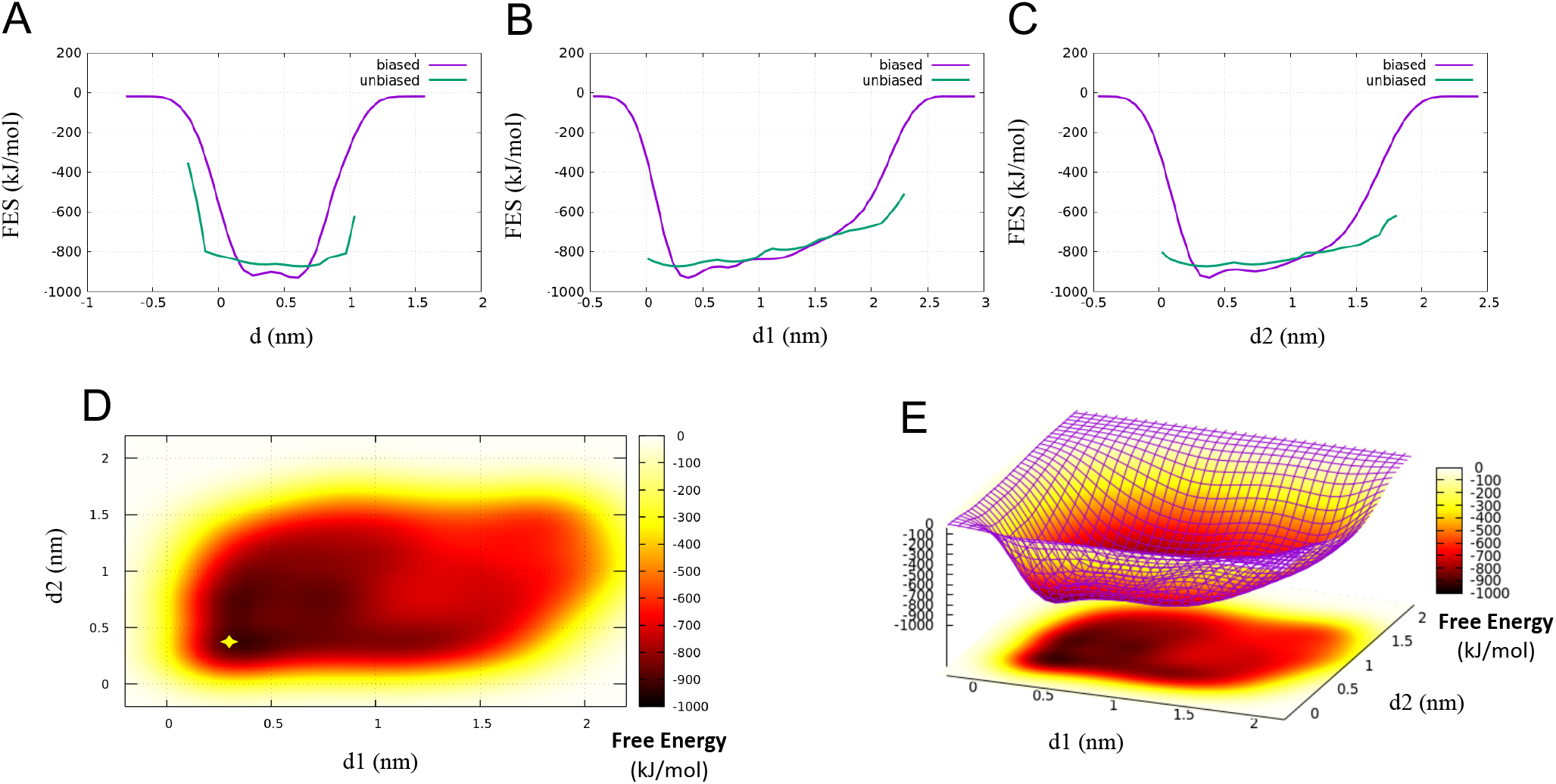
Comparison between the binding FES calculated from the metadynamics bias potential (biased, in purple) and by reweighting (unbiased, in green) as a function of **A)** d of CVs (COM distance between citrulline and RNA binding pocket); **B)** d1 of CVs (the distance between CIR:O7 and U29:N3); **C)** d2 of CVs (the distance between CIR:N8 and G4:N7). Two-dimensional free energy landscape onto d1 and d2 **D)** presented in 2D; **E)** presented in 3D

**Figure 16.**
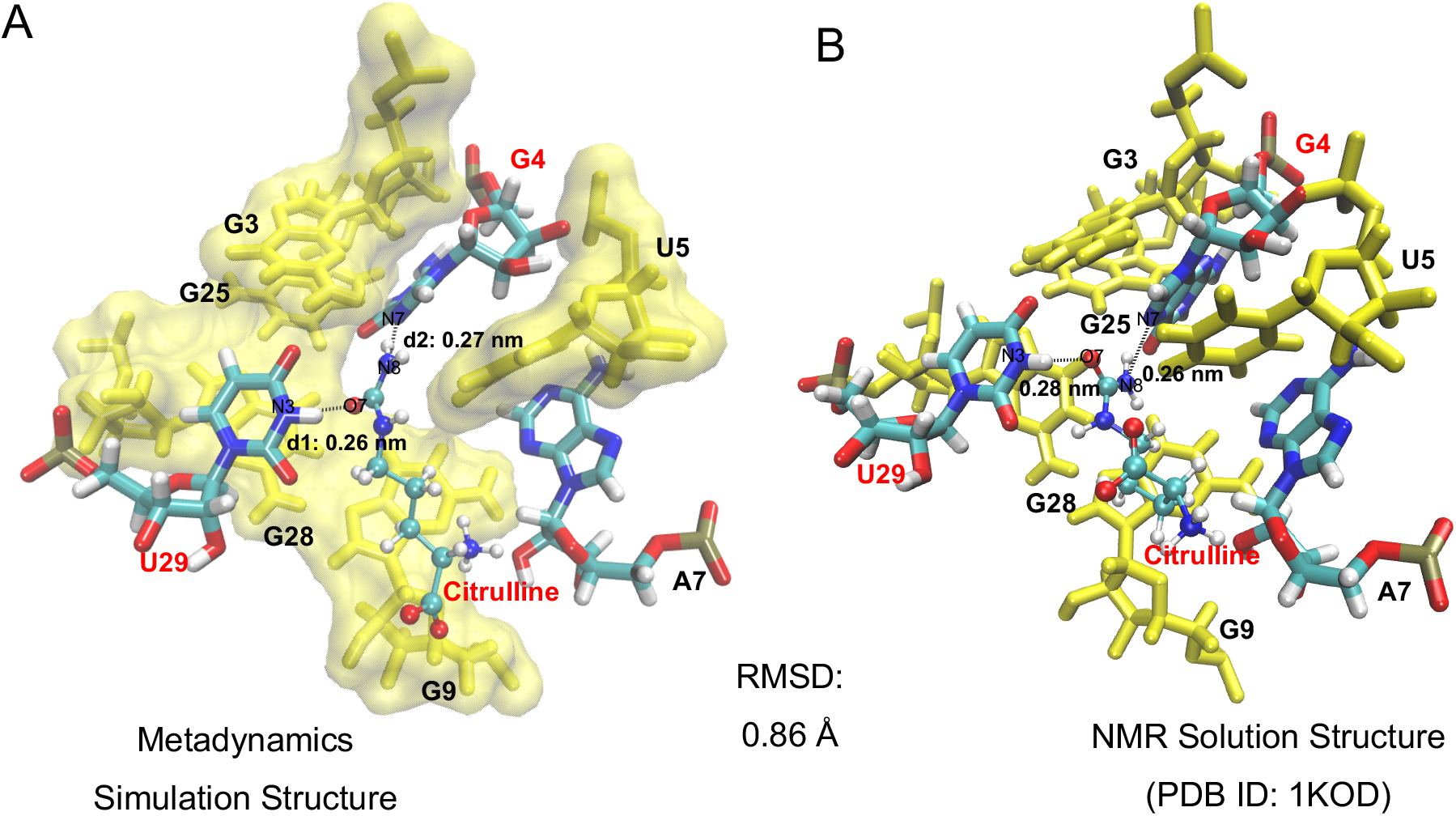
**A)** Snapshot of binding mode at energy basin of 2D free energy landscape onto d1 and d2 from metadynamics simulation; **B)** NMR solution structure of citrulline RNA aptamer complex (PDB ID: 1KOD)

### Arginine/RNA Aptamer

The last small molecule ligand-RNA system studied is arginine/RNA aptamer complex (PDB ID: 1KOC), which was evolved from the citrulline/RNA aptamer illustrated above. The arginine RNA aptamer differs from citrulline aptamer at only 3 nucleotides out of 44 as A3, G5 and C29. NMR solution structure showed that the conformation of arginine binding pocket is very common to citrulline RNA binding pocket with an uneven bottom plane of G9-G28-G25 and G4-G5 perpendicular to this surface. Therefore, we defined the arginine binding pocket of metadynamics simulation by the same positions of A3-G4-G5-A7-G9-G25-G28-C29. For cognate amino acid of arginine, the hydrogen binding interactions occurred between the guanidino group of arginine ligand and Watson-Crick edge of C29, Hoogsteen edge of G4 (Figure 19B). Thus, we chose the hydrogen bond donor-to-acceptor distance of ARG:NH2-H…C29:N3 as d1 of CVs, ARG:NH2-H…G4:O6 as d2 of CVs and the COM distance between arginine and RNA binding pocket as d of CVs.

In the metadynamics simulation, upper wall restrain forced arginine back into the binding pocket from 2.5 nm away. Within the range of d less that 0.7 nm, arginine has freedom of moving and rotating without being limited by restrains (Figure 17A). The Gaussian height of 0.6 kJ/mol with bias factor of 6.0 on the three CVs gave arginine sufficient energy for overcoming energy barriers of state transition so that arginine experienced various binding modes as many as possible within 60 ns (Movie S5). Whenever arginine approaching an energy favorable binding mode, height of Gaussian decreased to keep arginine staying in this state for longer. Whenever arginine approaching an energy unfavorable binding mode, height of Gaussian increased to push arginine leaving this state. Until the most energy favorable conformation was found, Gaussian height dropped to near zero without rebounding, as shown in (Figure 17B). According to the free energy surface of metadynamics (Figure 18), the deepest energy basin was located at d1=0.33 nm, d2=0.31 nm and d=6 nm from both biased and reweighted calculation, which referred to the conformation of arginine in the end of simulation when the two essential hydrogen bond of d1 and d2 formed at the same time (Figure 19A). As shown in (Figure 19), the most energy favorable binding mode of arginine has an RMSD of 0.64 Å from the NMR solution structure, which shows that metadynamics simulation prediction result is still consistent with experimental result. Moreover, the second energy favorable binding mode was found at d1=0.97 nm, d2=0.87 nm. In this binding mode (Figure 20), instead of the two hydrogen bonds of d1 and d2, we captured hydrogen bonding interactions between guanidino group of arginine and the phosphodiester backbond of RNA, amino group of arginine and Watson-Crick edge of C29.

**Figure 17.**
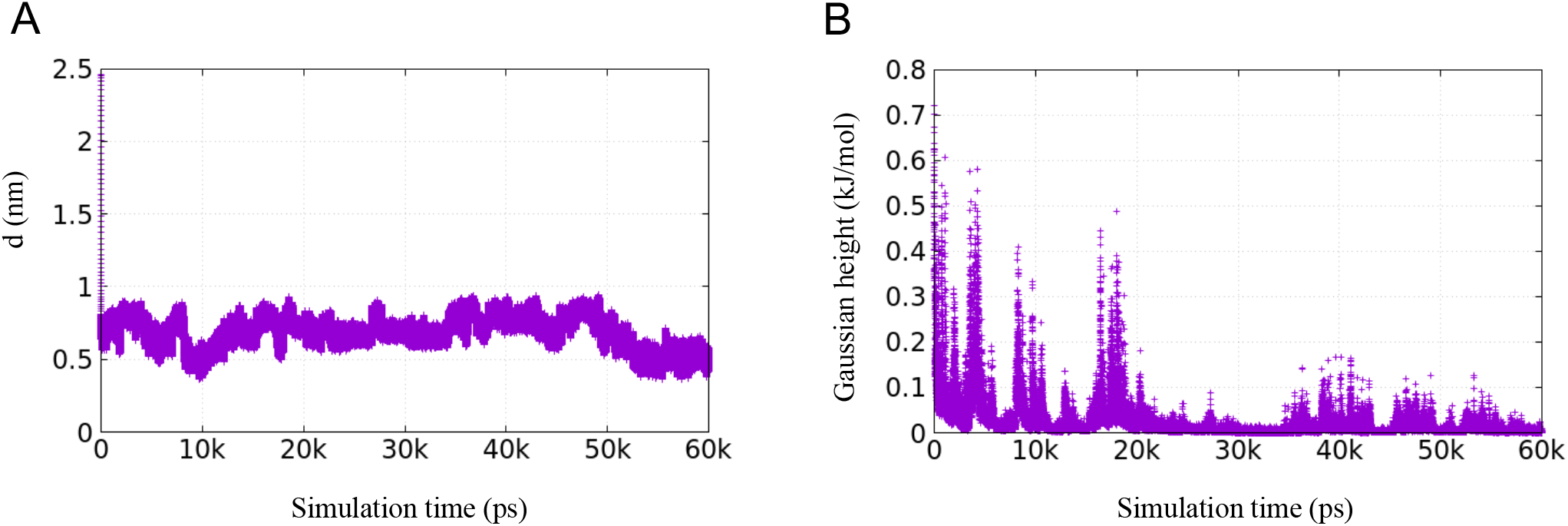
Time evolution of **A)** d of CVs (COM distance between arginine and RNA binding pocket defined by A3-G4-G5-A7-G9-G25-G28-C29) and **B)** Gaussian height in a 60ns-long well-tempered docking metadynamics simulation of arginine/RNA aptamer system.

**Figure 18.**
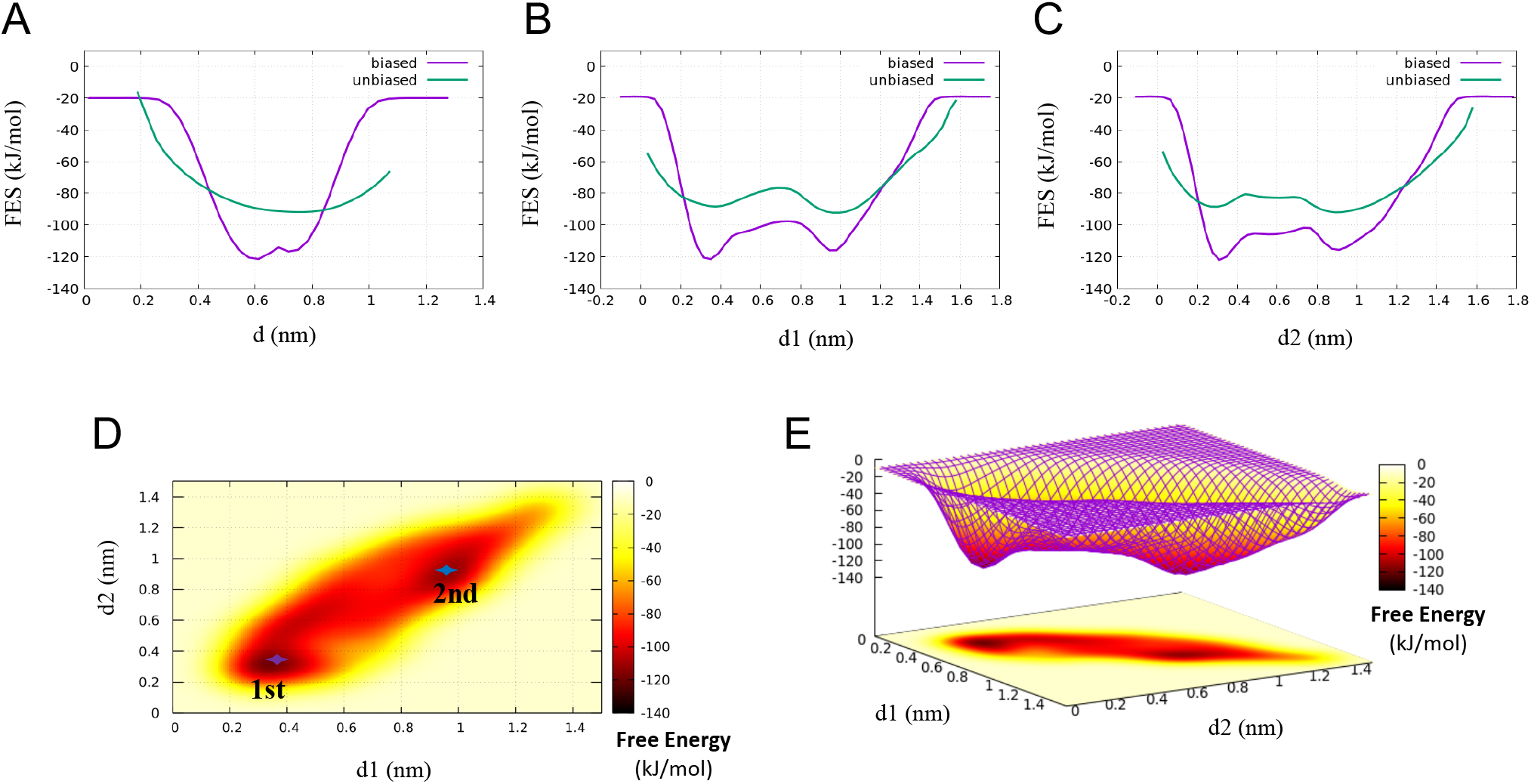
Comparison between the binding FES calculated from the metadynamics bias potential (biased, in purple) and by reweighting (unbiased, in green) as a function of **A)** d of CVs (COM distance between arginine and binding pocket); **B)** d1 of CVs (the distance between ARG:NH2 and C29:N3); **C)** d2 of CVs (the distance between ARG:NH2 and G4:O6). Two-dimensional free energy landscape onto d1 and d2 **D)** presented in 2D; **E)** presented in 3D

**Figure 19.**
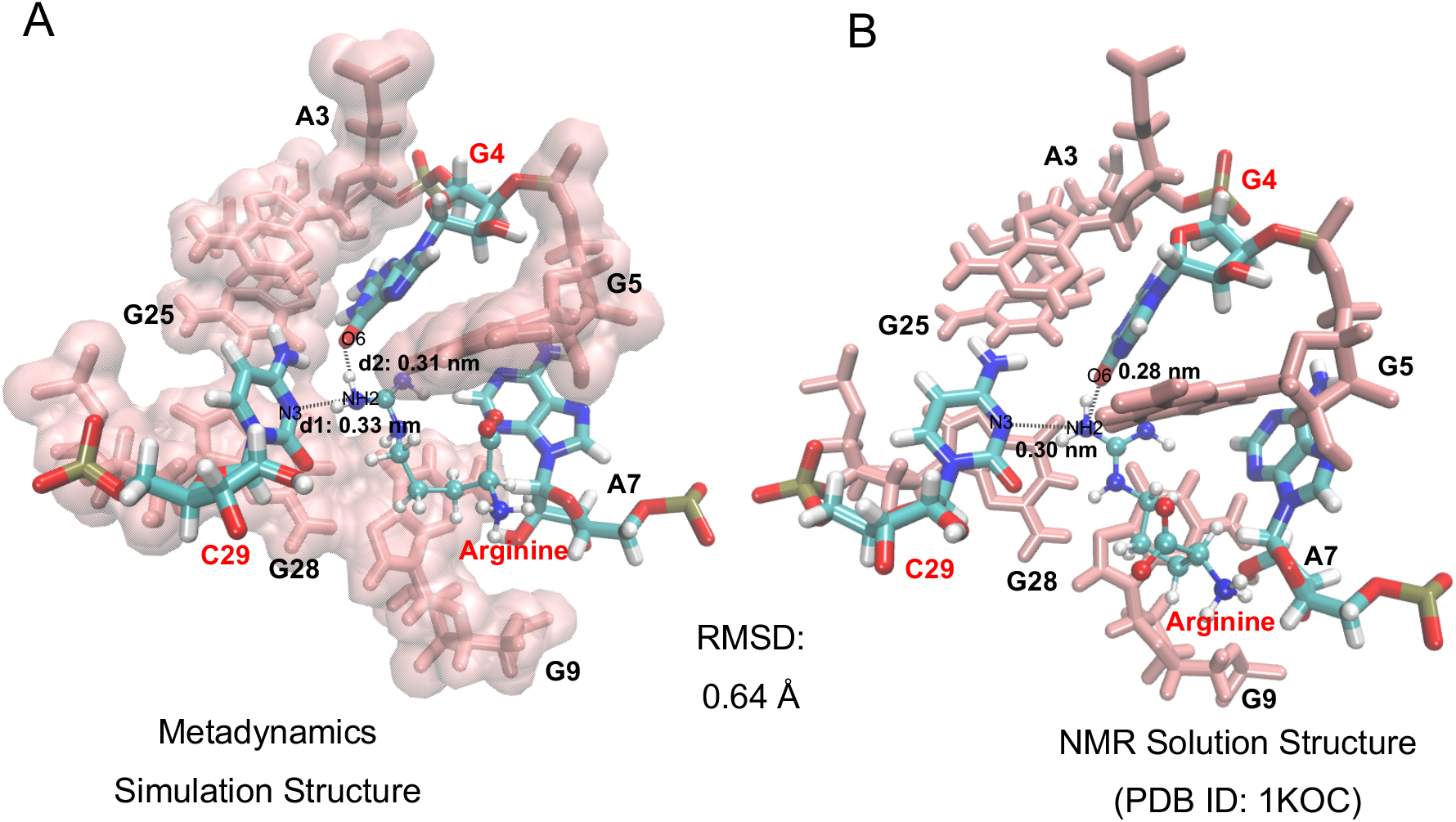
**A)** Snapshot of binding mode at energy basin of 2D free energy landscape onto d1 and d2 from metadynamics simulation; **B)** NMR solution structure of arginine-RNA aptamer complex (PDB ID: 1KOC)

**Figure 20.**
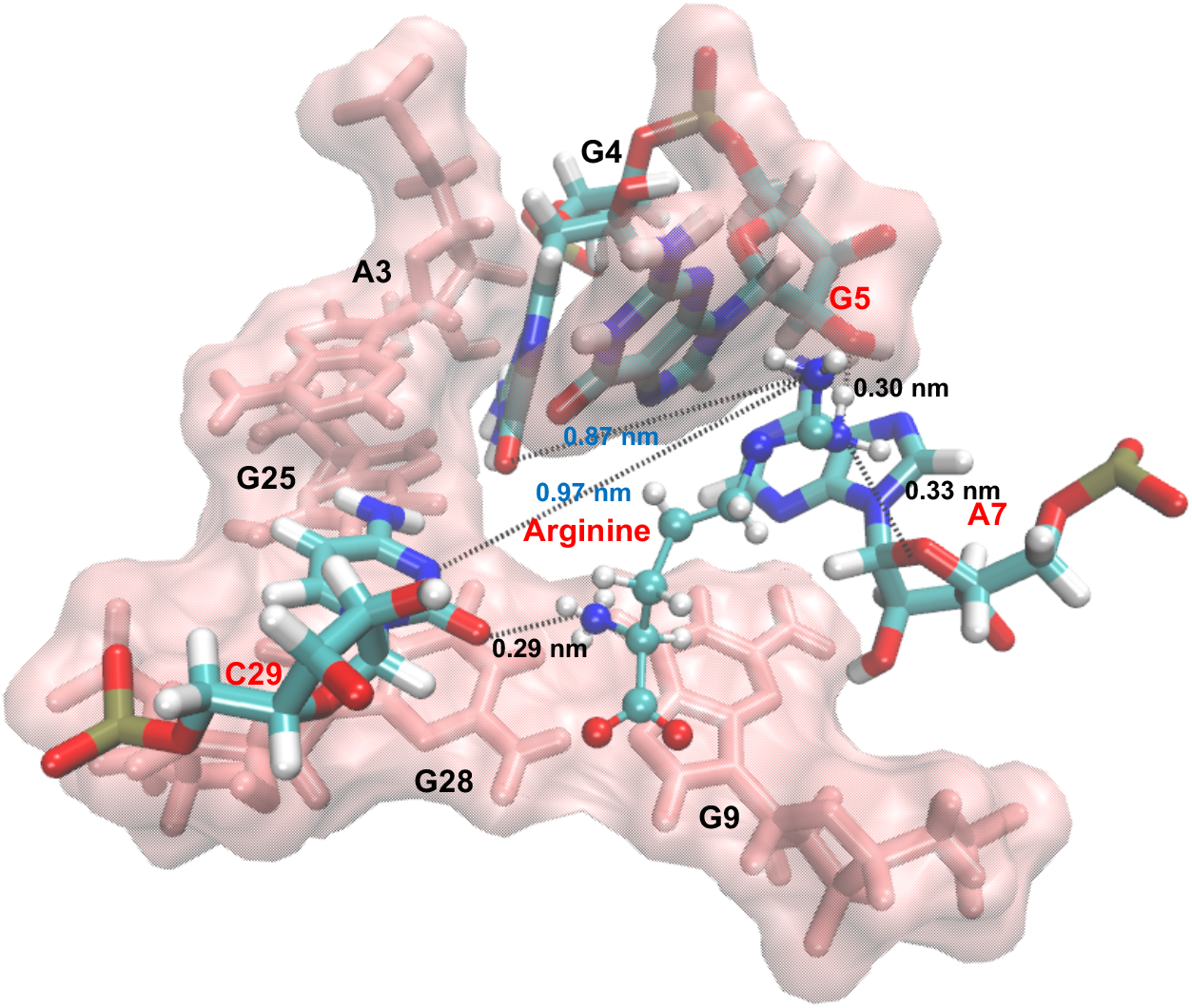
Snapshot of binding mode at the second energy basin of 2D free energy landscape onto d1 and d2 from metadynamics simulation.

## DISCUSSION

Elucidating the mechanism of molecular recognition process between small molecule ligands and its RNA binding pocket is always a key of concern in drug discovery. MD simulation has been an essential tool in interpretating structural dynamics of RNA and how they bind with small molecule ligands at atomic resolution. However, conventional unbiased MD simulations may miss many states with nonergodic dynamics due to the insurmountable high energy barriers in the binding pathway. To address this limitation, we applied well-tempered metadynamics coupled with upper wall restrain in this work, thus introducing an novel sampling approach for predicting binding modes of small molecule ligand recognizing RNA binding pocket. The two-dimensional free energy landscape of the whole binding process can be reconstructed in this method by incorporate couple possible hydrogen binding interactions between small molecule ligand and its RNA binding pocket, which can sample multiple transition states during the binding process and finally probe the most energy favorable binding modes accurately.

We calculate the free energy landscape as a function of three collective variables: the COM distance between small molecule ligand and its potential RNA binding pocket, the hydrogen bond donor-to-acceptor distances of two potential hydrogen bonds between small molecule ligand and its RNA binding pocket, which play the role of three coordinates on the free energy surface. The Gaussians on the three CVs gave the system enough potential for mimicking all possible dynamic motion and rotation of small molecule ligand during the binding process that cannot be reached by unbiased MD. The upper wall bias effectively ensured the ergodicity of metadynamics docking and the convergence of the free energy surface, avoiding invalid sampling. We studied five cases to exemplify the validity of the methodology: flavin mononucleotide (FMN) binding with 35-nucleotide RNA aptamer identified through in vitro selection, preQ1 riboswitch aptamer recognized by its metabolite, ligand binding to adenine riboswitch aptamer domain, RNA aptamer complexed with citrulline and RNA aptamer complexed with arginine. Our simulation prediction of binding mode in each case is in quantitative agreement with structure solved by X-ray crystallography or NMR. Additionally, we presented the first molecular dynamics binding pathway and binding mechanism for the three cases of in vitro selected RNA aptamer-small molecule ligand system.

The good agreement of our computational prediction with experimentally solved geometry and the molecular dynamics insights presented in this work demonstrates that metadynamics can be applied to effectively sampling of state transitions in ligand binding process and accurately predicting of binding modes via suitable selection of collective variables, which we have benchmarked with structures solved by X-ray crystallography or NMR. By coupling with upper wall restrain, we have enabled fast free energy profile calculation of small molecule binding process. In general, our simulation strategy will enable a faster and more detailed RNA-ligand binding investigation, facilitating a direct comparison of molecular modeling to experimental measurements. Moreover, our approach offers a valuable design and characterization tool for the growing community interested in applying in silico prediction to drug development of RNA-targeted small molecule compounds.

## SUPPORTING INFORMATION

Movie of metadynamics simulation for FMN/RNA aptamer complex (Movie S1);

Movie of metadynamics simulation for PreQ1 riboswitch aptamer complex (Movie S2);

Movie of metadynamics simulation for adenine riboswitch aptamer complex (Movie S3);

Movie of metadynamics simulation for citrulline/RNA aptamer complex (Movie S4);

Movie of metadynamics simulation for arginine/RNA aptamer complex (Movie S5);

Parameterization of five small molecule ligands (File S1).

## ACKNOWLEDGEMENTS

This work used computing resources provided by the Extreme Science and Engineering Discovery Environment (XSEDE) [allocation TG-MCB140273 to A.A.C], which is supported by National Science Foundation grant ACI-1548562.

