## Supplementary material for "Predicting Small Molecule Ligand – RNA Binding Pocket Binding Modes Using Metadynamics": File S1

### PARAMETERIZATION OF FIVE SMALL MOLECULE LIGANDS

#### FMN Parameterization

```
[ atomtypes ]
;name   bond_type     mass     charge   ptype   sigma         epsilon       Amb
 nc       nc          0.00000  0.00000   A     3.08750e-01   5.69031e-01 ; 1.82  0.1700
 c        c           0.00000  0.00000   A     3.22968e-01   2.87859e-01 ; 1.91  0.0860
 o        o           0.00000  0.00000   A     2.95992e-01   8.78640e-01 ; 1.66  0.2100
 or       o           0.00000  0.00000   A     2.91192e-01   7.02912e-01 ; 1.66  0.2100
 n        n           0.00000  0.00000   A     3.08750e-01   5.69031e-01 ; 1.82  0.1700
 hn       hn          0.00000  0.00000   A     1.06908e-01   6.56888e-02 ; 0.60  0.0157
 cd       cd          0.00000  0.00000   A     3.22968e-01   2.87859e-01 ; 1.91  0.0860
 ca       ca          0.00000  0.00000   A     3.22968e-01   2.87859e-01 ; 1.91  0.0860
 ha       ha          0.00000  0.00000   A     2.59964e-01   6.27600e-02 ; 1.46  0.0150
 c3       c3          0.00000  0.00000   A     3.39967e-01   4.57730e-01 ; 1.91  0.1094
 hc       hc          0.00000  0.00000   A     2.64953e-01   6.56888e-02 ; 1.49  0.0157
 na       na          0.00000  0.00000   A     3.08750e-01   5.69031e-01 ; 1.82  0.1700
 h1       h1          0.00000  0.00000   A     2.47135e-01   6.56888e-02 ; 1.39  0.0157
 oh       oh          0.00000  0.00000   A     3.06647e-01   8.80314e-01 ; 1.72  0.2104
 ho       ho          0.00000  0.00000   A     0.00000e+00   0.00000e+00 ; 0.00  0.0000
 os       os          0.00000  0.00000   A     3.00001e-01   7.11280e-01 ; 1.68  0.1700
 p5       p5          0.00000  0.00000   A     3.74177e-01   8.36800e-01 ; 2.10  0.2000

[ moleculetype ]
;name            nrexcl
 FMN              3

[ atoms ]
;   nr  type  resi  res  atom  cgnr     charge      mass       ; qtot   bond_type
     1   nc     1   FMN    N1    1    -0.629401     14.01000 ; qtot -0.629
     2    c     1   FMN    C2    2     0.728601     12.01000 ; qtot 0.099
     3   or     1   FMN    O2    3    -0.628901     16.00000 ; qtot -0.530
     4    n     1   FMN    N3    4    -0.446500     14.01000 ; qtot -0.976
     5   hn     1   FMN   HN3    5     0.302200      1.00800 ; qtot -0.674
     6    c     1   FMN    C4    6     0.455900     12.01000 ; qtot -0.218
     7   or     1   FMN    O4    7    -0.566301     16.00000 ; qtot -0.784
     8   cd     1   FMN   C4A    8     0.339500     12.01000 ; qtot -0.445
     9   nc     1   FMN    N5    9    -0.497400     14.01000 ; qtot -0.942
    10   ca     1   FMN   C5A   10     0.266000     12.01000 ; qtot -0.676
    11   ca     1   FMN    C6   11    -0.238800     12.01000 ; qtot -0.915
    12   ha     1   FMN    H6   12     0.148000      1.00800 ; qtot -0.767
    13   ca     1   FMN    C7   13     0.046000     12.01000 ; qtot -0.721
    14   c3     1   FMN   C7M   14    -0.203000     12.01000 ; qtot -0.924
    15   hc     1   FMN  HM71   15     0.055100      1.00800 ; qtot -0.869
    16   hc     1   FMN  HM72   16     0.055100      1.00800 ; qtot -0.814
    17   hc     1   FMN  HM73   17     0.055100      1.00800 ; qtot -0.759
    18   ca     1   FMN    C8   18     0.082400     12.01000 ; qtot -0.676
    19   c3     1   FMN   C8M   19    -0.022900     12.01000 ; qtot -0.699
    20   hc     1   FMN  HM81   20     0.039100      1.00800 ; qtot -0.660
    21   hc     1   FMN  HM82   21     0.039100      1.00800 ; qtot -0.621
    22   hc     1   FMN  HM83   22     0.039100      1.00800 ; qtot -0.582
    23   ca     1   FMN    C9   23    -0.247200     12.01000 ; qtot -0.829
    24   ha     1   FMN    H9   24     0.219200      1.00800 ; qtot -0.610
    25   ca     1   FMN   C9A   25    -0.011000     12.01000 ; qtot -0.621
    26   na     1   FMN   N10   26     0.020800     14.01000 ; qtot -0.600
    27   cd     1   FMN   C10   27     0.304800     12.01000 ; qtot -0.295
    28   c3     1   FMN   C1'   28    -0.066300     12.01000 ; qtot -0.362
    29   h1     1   FMN  H1'1   29     0.073800      1.00800 ; qtot -0.288
    30   h1     1   FMN  H1'2   30     0.073800      1.00800 ; qtot -0.214
    31   c3     1   FMN   C2'   31     0.148400     12.01000 ; qtot -0.066
    32   h1     1   FMN   H2'   32     0.078700      1.00800 ; qtot 0.013
    33   oh     1   FMN   O2'   33    -0.605401     16.00000 ; qtot -0.592
    34   ho     1   FMN  HO2'   34     0.412500      1.00800 ; qtot -0.180
    35   c3     1   FMN   C3'   35     0.123400     12.01000 ; qtot -0.057
    36   h1     1   FMN   H3'   36     0.055300      1.00800 ; qtot -0.001
    37   oh     1   FMN   O3'   37    -0.685901     16.00000 ; qtot -0.687
    38   ho     1   FMN  HO3'   38     0.456100      1.00800 ; qtot -0.231
    39   c3     1   FMN   C4'   39     0.072100     12.01000 ; qtot -0.159
    40   h1     1   FMN   H4'   40     0.103400      1.00800 ; qtot -0.056
```

```
    41   oh     1   FMN   O4'   41    -0.706901     16.00000 ; qtot -0.762
    42   ho     1   FMN  HO4'   42     0.466000      1.00800 ; qtot -0.296
    43   c3     1   FMN   C5'   43     0.075900     12.01000 ; qtot -0.221
    44   h1     1   FMN  H5'1   44     0.041600      1.00800 ; qtot -0.179
    45   h1     1   FMN  H5'2   45     0.041600      1.00800 ; qtot -0.137
    46   os     1   FMN   O5'   46    -0.476000     16.00000 ; qtot -0.613
    47   p5     1   FMN     P   47     1.260803     30.97000 ; qtot 0.648
    48    o     1   FMN   O1P   48    -0.882501     16.00000 ; qtot -0.235
    49    o     1   FMN   O2P   49    -0.882501     16.00000 ; qtot -1.117
    50    o     1   FMN   O3P   50    -0.882501     16.00000 ; qtot -2.000

[ bonds ]
;   ai     aj funct   r             k
     1      2  1    1.3784e-01    3.5840e+05 ;     N1 - C2
     1     27  1    1.3350e-01    4.1388e+05 ;     N1 - C10
     2      3  1    1.2140e-01    5.4225e+05 ;     C2 - O2
     2      4  1    1.3450e-01    4.0016e+05 ;     C2 - N3
     4      5  1    1.0090e-01    3.4326e+05 ;     N3 - HN3
     4      6  1    1.3450e-01    4.0016e+05 ;     N3 - C4
     6      7  1    1.2140e-01    5.4225e+05 ;     C4 - O4
     6      8  1    1.4620e-01    3.1581e+05 ;     C4 - C4A
     8      9  1    1.3350e-01    4.1388e+05 ;    C4A - N5
     8     27  1    1.4290e-01    3.5003e+05 ;    C4A - C10
     9     10  1    1.3360e-01    4.1246e+05 ;     N5 - C5A
    10     11  1    1.3870e-01    4.0033e+05 ;    C5A - C6
    10     25  1    1.3870e-01    4.0033e+05 ;    C5A - C9A
    11     12  1    1.0870e-01    2.8811e+05 ;     C6 - H6
    11     13  1    1.3870e-01    4.0033e+05 ;     C6 - C7
    13     14  1    1.5130e-01    2.7070e+05 ;     C7 - C7M
    13     18  1    1.3870e-01    4.0033e+05 ;     C7 - C8
    14     15  1    1.0920e-01    2.8225e+05 ;    C7M - HM71
    14     16  1    1.0920e-01    2.8225e+05 ;    C7M - HM72
    14     17  1    1.0920e-01    2.8225e+05 ;    C7M - HM73
    18     19  1    1.5130e-01    2.7070e+05 ;     C8 - C8M
    18     23  1    1.3870e-01    4.0033e+05 ;     C8 - C9
    19     20  1    1.0920e-01    2.8225e+05 ;    C8M - HM81
    19     21  1    1.0920e-01    2.8225e+05 ;    C8M - HM82
    19     22  1    1.0920e-01    2.8225e+05 ;    C8M - HM83
    23     24  1    1.0870e-01    2.8811e+05 ;     C9 - H9
    23     25  1    1.3870e-01    4.0033e+05 ;     C9 - C9A
    25     26  1    1.3500e-01    3.9355e+05 ;    C9A - N10
    26     27  1    1.3710e-01    3.6719e+05 ;    N10 - C10
    26     28  1    1.4560e-01    2.8008e+05 ;    N10 - C1'
    28     29  1    1.0930e-01    2.8108e+05 ;    C1' - H1'1
    28     30  1    1.0930e-01    2.8108e+05 ;    C1' - H1'2
    28     31  1    1.5350e-01    2.5363e+05 ;    C1' - C2'
    31     32  1    1.0930e-01    2.8108e+05 ;    C2' - H2'
    31     33  1    1.4260e-01    2.6284e+05 ;    C2' - O2'
    31     35  1    1.5350e-01    2.5363e+05 ;    C2' - C3'
    33     34  1    9.7400e-02    3.0928e+05 ;    O2' - HO2'
    35     36  1    1.0930e-01    2.8108e+05 ;    C3' - H3'
    35     37  1    1.4260e-01    2.6284e+05 ;    C3' - O3'
    35     39  1    1.5350e-01    2.5363e+05 ;    C3' - C4'
    37     38  1    9.7400e-02    3.0928e+05 ;    O3' - HO3'
    39     40  1    1.0930e-01    2.8108e+05 ;    C4' - H4'
    39     41  1    1.4260e-01    2.6284e+05 ;    C4' - O4'
    39     43  1    1.5350e-01    2.5363e+05 ;    C4' - C5'
    41     42  1    9.7400e-02    3.0928e+05 ;    O4' - HO4'
    43     44  1    1.0930e-01    2.8108e+05 ;    C5' - H5'1
    43     45  1    1.0930e-01    2.8108e+05 ;    C5' - H5'2
    43     46  1    1.4390e-01    2.5230e+05 ;    C5' - O5'
    46     47  1    1.6020e-01    2.8660e+05 ;    O5' - P
    47     48  1    1.4810e-01    4.0811e+05 ;      P - O1P
    47     49  1    1.4810e-01    4.0811e+05 ;      P - O2P
    47     50  1    1.4810e-01    4.0811e+05 ;      P - O3P

[ pairs ]
;   ai     aj    funct
     1      5      1 ;     N1 - HN3
     1      6      1 ;     N1 - C4
     1      9      1 ;     N1 - N5
     1     25      1 ;     N1 - C9A
```

```
   1     28      1 ;     N1 - C1'
   2      7      1 ;     C2 - O4
   2      8      1 ;     C2 - C4A
   2     26      1 ;     C2 - N10
   3      5      1 ;     O2 - HN3
   3      6      1 ;     O2 - C4
   4      9      1 ;     N3 - N5
   5      7      1 ;    HN3 - O4
   5      8      1 ;    HN3 - C4A
   6     10      1 ;     C4 - C5A
   6     26      1 ;     C4 - N10
   7      9      1 ;     O4 - N5
   7     27      1 ;     O4 - C10
   8     11      1 ;    C4A - C6
   8     25      1 ;    C4A - C9A
   8     28      1 ;    C4A - C1'
   9     12      1 ;     N5 - H6
   9     13      1 ;     N5 - C7
   9     23      1 ;     N5 - C9
   9     26      1 ;     N5 - N10
  10     14      1 ;    C5A - C7M
  10     18      1 ;    C5A - C8
  10     24      1 ;    C5A - H9
  10     27      1 ;    C5A - C10
  10     28      1 ;    C5A - C1'
  11     15      1 ;     C6 - HM71
  11     16      1 ;     C6 - HM72
  11     17      1 ;     C6 - HM73
  11     19      1 ;     C6 - C8M
  11     23      1 ;     C6 - C9
  11     26      1 ;     C6 - N10
  12     14      1 ;     H6 - C7M
  12     18      1 ;     H6 - C8
  12     25      1 ;     H6 - C9A
  13     20      1 ;     C7 - HM81
  13     21      1 ;     C7 - HM82
  13     22      1 ;     C7 - HM83
  13     24      1 ;     C7 - H9
  13     25      1 ;     C7 - C9A
  14     19      1 ;    C7M - C8M
  14     23      1 ;    C7M - C9
  15     18      1 ;   HM71 - C8
  16     18      1 ;   HM72 - C8
  17     18      1 ;   HM73 - C8
  18     26      1 ;     C8 - N10
  19     24      1 ;    C8M - H9
  19     25      1 ;    C8M - C9A
  20     23      1 ;   HM81 - C9
  21     23      1 ;   HM82 - C9
  22     23      1 ;   HM83 - C9
  23     27      1 ;     C9 - C10
  23     28      1 ;     C9 - C1'
  24     26      1 ;     H9 - N10
  25     29      1 ;    C9A - H1'1
  25     30      1 ;    C9A - H1'2
  25     31      1 ;    C9A - C2'
  26     32      1 ;    N10 - H2'
  26     33      1 ;    N10 - O2'
  26     35      1 ;    N10 - C3'
  27      3      1 ;    C10 - O2
  27      4      1 ;    C10 - N3
  27     29      1 ;    C10 - H1'1
  27     30      1 ;    C10 - H1'2
  27     31      1 ;    C10 - C2'
  28     34      1 ;    C1' - HO2'
  28     36      1 ;    C1' - H3'
  28     37      1 ;    C1' - O3'
  28     39      1 ;    C1' - C4'
  29     32      1 ;   H1'1 - H2'
  29     33      1 ;   H1'1 - O2'
  29     35      1 ;   H1'1 - C3'
```

```
    30     32      1 ;   H1'2 - H2'
    30     33      1 ;   H1'2 - O2'
    30     35      1 ;   H1'2 - C3'
    31     38      1 ;    C2' - HO3'
    31     40      1 ;    C2' - H4'
    31     41      1 ;    C2' - O4'
    31     43      1 ;    C2' - C5'
    32     34      1 ;    H2' - HO2'
    32     36      1 ;    H2' - H3'
    32     37      1 ;    H2' - O3'
    32     39      1 ;    H2' - C4'
    33     36      1 ;    O2' - H3'
    33     37      1 ;    O2' - O3'
    33     39      1 ;    O2' - C4'
    34     35      1 ;   HO2' - C3'
    35     42      1 ;    C3' - HO4'
    35     44      1 ;    C3' - H5'1
    35     45      1 ;    C3' - H5'2
    35     46      1 ;    C3' - O5'
    36     38      1 ;    H3' - HO3'
    36     40      1 ;    H3' - H4'
    36     41      1 ;    H3' - O4'
    36     43      1 ;    H3' - C5'
    37     40      1 ;    O3' - H4'
    37     41      1 ;    O3' - O4'
    37     43      1 ;    O3' - C5'
    38     39      1 ;   HO3' - C4'
    39     47      1 ;    C4' - P
    40     42      1 ;    H4' - HO4'
    40     44      1 ;    H4' - H5'1
    40     45      1 ;    H4' - H5'2
    40     46      1 ;    H4' - O5'
    41     44      1 ;    O4' - H5'1
    41     45      1 ;    O4' - H5'2
    41     46      1 ;    O4' - O5'
    42     43      1 ;   HO4' - C5'
    43     48      1 ;    C5' - O1P
    43     49      1 ;    C5' - O2P
    43     50      1 ;    C5' - O3P
    44     47      1 ;   H5'1 - P
    45     47      1 ;   H5'2 - P

[ angles ]
;   ai     aj     ak    funct   theta         cth
     1      2      3      1    1.2398e+02    6.1965e+02 ;     N1 - C2     - O2
     1      2      4      1    1.1705e+02    6.0827e+02 ;     N1 - C2     - N3
     1     27      8      1    1.1256e+02    5.9538e+02 ;     N1 - C10    - C4A
     1     27     26      1    1.1202e+02    6.2576e+02 ;     N1 - C10    - N10
     2      1     27      1    1.2032e+02    5.5689e+02 ;     C2 - N1     - C10
     2      4      5      1    1.1846e+02    4.1179e+02 ;     C2 - N3     - HN3
     2      4      6      1    1.2714e+02    5.4668e+02 ;     C2 - N3     - C4
     3      2      4      1    1.2203e+02    6.3455e+02 ;     O2 - C2     - N3
     4      6      7      1    1.2203e+02    6.3455e+02 ;     N3 - C4     - O4
     4      6      8      1    1.1186e+02    5.8735e+02 ;     N3 - C4     - C4A
     5      4      6      1    1.1846e+02    4.1179e+02 ;    HN3 - N3     - C4
     6      8      9      1    1.2186e+02    5.6442e+02 ;     C4 - C4A    - N5
     6      8     27      1    1.2269e+02    5.3321e+02 ;     C4 - C4A    - C10
     7      6      8      1    1.2571e+02    5.7664e+02 ;     O4 - C4     - C4A
     8      9     10      1    1.0424e+02    6.0810e+02 ;    C4A - N5     - C5A
     8     27     26      1    1.0680e+02    6.0425e+02 ;    C4A - C10    - N10
     9      8     27      1    1.1256e+02    5.9538e+02 ;     N5 - C4A    - C10
     9     10     11      1    1.1972e+02    5.8693e+02 ;     N5 - C5A    - C6
     9     10     25      1    1.1972e+02    5.8693e+02 ;     N5 - C5A    - C9A
    10     11     12      1    1.2001e+02    4.0551e+02 ;    C5A - C6     - H6
    10     11     13      1    1.1997e+02    5.6216e+02 ;    C5A - C6     - C7
    10     25     23      1    1.1997e+02    5.6216e+02 ;    C5A - C9A    - C9
    10     25     26      1    1.1834e+02    5.8752e+02 ;    C5A - C9A    - N10
    11     10     25      1    1.1997e+02    5.6216e+02 ;     C6 - C5A    - C9A
    11     13     14      1    1.2063e+02    5.3421e+02 ;     C6 - C7     - C7M
    11     13     18      1    1.1997e+02    5.6216e+02 ;     C6 - C7     - C8
    12     11     13      1    1.2001e+02    4.0551e+02 ;     H6 - C6     - C7
```

```
   13      14      15       1    1.1015e+02    3.9296e+02 ;      C7 - C7M    - HM71
   13      14      16       1    1.1015e+02    3.9296e+02 ;      C7 - C7M    - HM72
   13      14      17       1    1.1015e+02    3.9296e+02 ;      C7 - C7M    - HM73
   13      18      19       1    1.2063e+02    5.3421e+02 ;      C7 - C8     - C8M
   13      18      23       1    1.1997e+02    5.6216e+02 ;      C7 - C8     - C9
   14      13      18       1    1.2063e+02    5.3421e+02 ;     C7M - C7     - C8
   15      14      16       1    1.0835e+02    3.2995e+02 ;    HM71 - C7M    - HM72
   15      14      17       1    1.0835e+02    3.2995e+02 ;    HM71 - C7M    - HM73
   16      14      17       1    1.0835e+02    3.2995e+02 ;    HM72 - C7M    - HM73
   18      19      20       1    1.1015e+02    3.9296e+02 ;      C8 - C8M    - HM81
   18      19      21       1    1.1015e+02    3.9296e+02 ;      C8 - C8M    - HM82
   18      19      22       1    1.1015e+02    3.9296e+02 ;      C8 - C8M    - HM83
   18      23      24       1    1.2001e+02    4.0551e+02 ;      C8 - C9     - H9
   18      23      25       1    1.1997e+02    5.6216e+02 ;      C8 - C9     - C9A
   19      18      23       1    1.2063e+02    5.3421e+02 ;     C8M - C8     - C9
   20      19      21       1    1.0835e+02    3.2995e+02 ;    HM81 - C8M    - HM82
   20      19      22       1    1.0835e+02    3.2995e+02 ;    HM81 - C8M    - HM83
   21      19      22       1    1.0835e+02    3.2995e+02 ;    HM82 - C8M    - HM83
   23      25      26       1    1.1834e+02    5.8752e+02 ;      C9 - C9A    - N10
   24      23      25       1    1.2001e+02    4.0551e+02 ;      H9 - C9     - C9A
   25      26      27       1    1.1315e+02    5.7287e+02 ;     C9A - N10    - C10
   25      26      28       1    1.2436e+02    5.2844e+02 ;     C9A - N10    - C1'
   26      28      29       1    1.0945e+02    4.1756e+02 ;     N10 - C1'    - H1'1
   26      28      30       1    1.0945e+02    4.1756e+02 ;     N10 - C1'    - H1'2
   26      28      31       1    1.1281e+02    5.5003e+02 ;     N10 - C1'    - C2'
   27      26      28       1    1.2509e+02    5.2350e+02 ;     C10 - N10    - C1'
   28      31      32       1    1.1007e+02    3.8794e+02 ;     C1' - C2'    - H2'
   28      31      33       1    1.0943e+02    5.6668e+02 ;     C1' - C2'    - O2'
   28      31      35       1    1.1063e+02    5.2894e+02 ;     C1' - C2'    - C3'
   29      28      30       1    1.0955e+02    3.2786e+02 ;    H1'1 - C1'    - H1'2
   29      28      31       1    1.1007e+02    3.8794e+02 ;    H1'1 - C1'    - C2'
   30      28      31       1    1.1007e+02    3.8794e+02 ;    H1'2 - C1'    - C2'
   31      33      34       1    1.0816e+02    3.9405e+02 ;     C2' - O2'    - HO2'
   31      35      36       1    1.1007e+02    3.8794e+02 ;     C2' - C3'    - H3'
   31      35      37       1    1.0943e+02    5.6668e+02 ;     C2' - C3'    - O3'
   31      35      39       1    1.1063e+02    5.2894e+02 ;     C2' - C3'    - C4'
   32      31      33       1    1.0988e+02    4.2652e+02 ;     H2' - C2'    - O2'
   32      31      35       1    1.1007e+02    3.8794e+02 ;     H2' - C2'    - C3'
   33      31      35       1    1.0943e+02    5.6668e+02 ;     O2' - C2'    - C3'
   35      37      38       1    1.0816e+02    3.9405e+02 ;     C3' - O3'    - HO3'
   35      39      40       1    1.1007e+02    3.8794e+02 ;     C3' - C4'    - H4'
   35      39      41       1    1.0943e+02    5.6668e+02 ;     C3' - C4'    - O4'
   35      39      43       1    1.1063e+02    5.2894e+02 ;     C3' - C4'    - C5'
   36      35      37       1    1.0988e+02    4.2652e+02 ;     H3' - C3'    - O3'
   36      35      39       1    1.1007e+02    3.8794e+02 ;     H3' - C3'    - C4'
   37      35      39       1    1.0943e+02    5.6668e+02 ;     O3' - C3'    - C4'
   39      41      42       1    1.0816e+02    3.9405e+02 ;     C4' - O4'    - HO4'
   39      43      44       1    1.1007e+02    3.8794e+02 ;     C4' - C5'    - H5'1
   39      43      45       1    1.1007e+02    3.8794e+02 ;     C4' - C5'    - H5'2
   39      43      46       1    1.0842e+02    5.6718e+02 ;     C4' - C5'    - O5'
   40      39      41       1    1.0988e+02    4.2652e+02 ;     H4' - C4'    - O4'
   40      39      43       1    1.1007e+02    3.8794e+02 ;     H4' - C4'    - C5'
   41      39      43       1    1.0943e+02    5.6668e+02 ;     O4' - C4'    - C5'
   43      46      47       1    1.1800e+02    5.1882e+02 ;     C5' - O5'    - P
   44      43      45       1    1.0955e+02    3.2786e+02 ;    H5'1 - C5'    - H5'2
   44      43      46       1    1.0882e+02    4.2543e+02 ;    H5'1 - C5'    - O5'
   45      43      46       1    1.0882e+02    4.2543e+02 ;    H5'2 - C5'    - O5'
   46      47      48       1    1.1609e+02    5.8861e+02 ;     O5' - P      - O1P
   46      47      49       1    1.1609e+02    5.8861e+02 ;     O5' - P      - O2P
   46      47      50       1    1.1609e+02    5.8861e+02 ;     O5' - P      - O3P
   48      47      49       1    1.1580e+02    6.1530e+02 ;     O1P - P      - O2P
   48      47      50       1    1.1580e+02    6.1530e+02 ;     O1P - P      - O3P
   49      47      50       1    1.1580e+02    6.1530e+02 ;     O2P - P      - O3P

[ dihedrals ] ; propers
; treated as RBs in GROMACS to use combine multiple AMBER torsions per quartet
;    i      j      k      l   func     C0           C1          C2           C3           C4           C5
     1      2      4      5      3   20.92000     0.00000  -20.92000     0.00000     0.00000     0.00000 ;     N1-    C2-    N3-    HN3
     1      2      4      6      3   20.92000     0.00000  -20.92000     0.00000     0.00000     0.00000 ;     N1-    C2-    N3-     C4
     1     27      8      6      3   33.47200     0.00000  -33.47200     0.00000     0.00000     0.00000 ;     N1-   C10-   C4A-     C4
     1     27      8      9      3   33.47200     0.00000  -33.47200     0.00000     0.00000     0.00000 ;     N1-   C10-   C4A-     N5
     1     27     26     25      3   14.22560     0.00000  -14.22560     0.00000     0.00000     0.00000 ;     N1-   C10-   N10-    C9A
     1     27     26     28      3   14.22560     0.00000  -14.22560     0.00000     0.00000     0.00000 ;     N1-   C10-   N10-    C1'
     2      1     27      8      3   39.74800     0.00000  -39.74800     0.00000     0.00000     0.00000 ;     C2-    N1-   C10-    C4A
```

| | | | | | | | | | | | | | | | |
|---|---|---|---|---|---|---|---|---|---|---|---|---|---|---|---|
| 2 | 1 | 27 | 26 | 3 | 39.74800 | 0.00000 | -39.74800 | 0.00000 | 0.00000 | 0.00000 | ; | C2- | N1- | C10- | N10 |
| 2 | 4 | 6 | 7 | 3 | 20.92000 | 0.00000 | -20.92000 | 0.00000 | 0.00000 | 0.00000 | ; | C2- | N3- | C4- | O4 |
| 2 | 4 | 6 | 8 | 3 | 20.92000 | 0.00000 | -20.92000 | 0.00000 | 0.00000 | 0.00000 | ; | C2- | N3- | C4- | C4A |
| 3 | 2 | 4 | 5 | 3 | 29.28800 | -8.36800 | -20.92000 | 0.00000 | 0.00000 | 0.00000 | ; | O2- | C2- | N3- | HN3 |
| 3 | 2 | 4 | 6 | 3 | 20.92000 | 0.00000 | -20.92000 | 0.00000 | 0.00000 | 0.00000 | ; | O2- | C2- | N3- | C4 |
| 4 | 6 | 8 | 9 | 3 | 24.05800 | 0.00000 | -24.05800 | 0.00000 | 0.00000 | 0.00000 | ; | N3- | C4- | C4A- | N5 |
| 4 | 6 | 8 | 27 | 3 | 24.05800 | 0.00000 | -24.05800 | 0.00000 | 0.00000 | 0.00000 | ; | N3- | C4- | C4A- | C10 |
| 5 | 4 | 6 | 7 | 3 | 29.28800 | -8.36800 | -20.92000 | 0.00000 | 0.00000 | 0.00000 | ; | HN3- | N3- | C4- | O4 |
| 5 | 4 | 6 | 8 | 3 | 20.92000 | 0.00000 | -20.92000 | 0.00000 | 0.00000 | 0.00000 | ; | HN3- | N3- | C4- | C4A |
| 6 | 8 | 9 | 10 | 3 | 39.74800 | 0.00000 | -39.74800 | 0.00000 | 0.00000 | 0.00000 | ; | C4- | C4A- | N5- | C5A |
| 6 | 8 | 27 | 26 | 3 | 33.47200 | 0.00000 | -33.47200 | 0.00000 | 0.00000 | 0.00000 | ; | C4- | C4A- | C10- | N10 |
| 7 | 6 | 8 | 9 | 3 | 24.05800 | 0.00000 | -24.05800 | 0.00000 | 0.00000 | 0.00000 | ; | O4- | C4- | C4A- | N5 |
| 7 | 6 | 8 | 27 | 3 | 24.05800 | 0.00000 | -24.05800 | 0.00000 | 0.00000 | 0.00000 | ; | O4- | C4- | C4A- | C10 |
| 8 | 9 | 10 | 11 | 3 | 40.16640 | 0.00000 | -40.16640 | 0.00000 | 0.00000 | 0.00000 | ; | C4A- | N5- | C5A- | C6 |
| 8 | 9 | 10 | 25 | 3 | 40.16640 | 0.00000 | -40.16640 | 0.00000 | 0.00000 | 0.00000 | ; | C4A- | N5- | C5A- | C9A |
| 8 | 27 | 26 | 25 | 3 | 14.22560 | 0.00000 | -14.22560 | 0.00000 | 0.00000 | 0.00000 | ; | C4A- | C10- | N10- | C9A |
| 8 | 27 | 26 | 28 | 3 | 14.22560 | 0.00000 | -14.22560 | 0.00000 | 0.00000 | 0.00000 | ; | C4A- | C10- | N10- | C1' |
| 9 | 8 | 27 | 26 | 3 | 33.47200 | 0.00000 | -33.47200 | 0.00000 | 0.00000 | 0.00000 | ; | N5- | C4A- | C10- | N10 |
| 9 | 10 | 11 | 12 | 3 | 30.33400 | 0.00000 | -30.33400 | 0.00000 | 0.00000 | 0.00000 | ; | N5- | C5A- | C6- | H6 |
| 9 | 10 | 11 | 13 | 3 | 30.33400 | 0.00000 | -30.33400 | 0.00000 | 0.00000 | 0.00000 | ; | N5- | C5A- | C6- | C7 |
| 9 | 10 | 25 | 23 | 3 | 30.33400 | 0.00000 | -30.33400 | 0.00000 | 0.00000 | 0.00000 | ; | N5- | C5A- | C9A- | C9 |
| 9 | 10 | 25 | 26 | 3 | 30.33400 | 0.00000 | -30.33400 | 0.00000 | 0.00000 | 0.00000 | ; | N5- | C5A- | C9A- | N10 |
| 10 | 9 | 8 | 27 | 3 | 39.74800 | 0.00000 | -39.74800 | 0.00000 | 0.00000 | 0.00000 | ; | C5A- | N5- | C4A- | C10 |
| 10 | 11 | 13 | 14 | 3 | 30.33400 | 0.00000 | -30.33400 | 0.00000 | 0.00000 | 0.00000 | ; | C5A- | C6- | C7- | C7M |
| 10 | 11 | 13 | 18 | 3 | 30.33400 | 0.00000 | -30.33400 | 0.00000 | 0.00000 | 0.00000 | ; | C5A- | C6- | C7- | C8 |
| 10 | 25 | 23 | 18 | 3 | 30.33400 | 0.00000 | -30.33400 | 0.00000 | 0.00000 | 0.00000 | ; | C5A- | C9A- | C9- | C8 |
| 10 | 25 | 23 | 24 | 3 | 30.33400 | 0.00000 | -30.33400 | 0.00000 | 0.00000 | 0.00000 | ; | C5A- | C9A- | C9- | H9 |
| 10 | 25 | 26 | 27 | 3 | 2.51040 | 0.00000 | -2.51040 | 0.00000 | 0.00000 | 0.00000 | ; | C5A- | C9A- | N10- | C10 |
| 10 | 25 | 26 | 28 | 3 | 2.51040 | 0.00000 | -2.51040 | 0.00000 | 0.00000 | 0.00000 | ; | C5A- | C9A- | N10- | C1' |
| 11 | 10 | 25 | 23 | 3 | 30.33400 | 0.00000 | -30.33400 | 0.00000 | 0.00000 | 0.00000 | ; | C6- | C5A- | C9A- | C9 |
| 11 | 10 | 25 | 26 | 3 | 30.33400 | 0.00000 | -30.33400 | 0.00000 | 0.00000 | 0.00000 | ; | C6- | C5A- | C9A- | N10 |
| 11 | 13 | 14 | 15 | 3 | 0.00000 | 0.00000 | 0.00000 | 0.00000 | 0.00000 | 0.00000 | ; | C6- | C7- | C7M- | HM71 |
| 11 | 13 | 14 | 16 | 3 | 0.00000 | 0.00000 | 0.00000 | 0.00000 | 0.00000 | 0.00000 | ; | C6- | C7- | C7M- | HM72 |
| 11 | 13 | 14 | 17 | 3 | 0.00000 | 0.00000 | 0.00000 | 0.00000 | 0.00000 | 0.00000 | ; | C6- | C7- | C7M- | HM73 |
| 11 | 13 | 18 | 19 | 3 | 30.33400 | 0.00000 | -30.33400 | 0.00000 | 0.00000 | 0.00000 | ; | C6- | C7- | C8- | C8M |
| 11 | 13 | 18 | 23 | 3 | 30.33400 | 0.00000 | -30.33400 | 0.00000 | 0.00000 | 0.00000 | ; | C6- | C7- | C8- | C9 |
| 12 | 11 | 10 | 25 | 3 | 30.33400 | 0.00000 | -30.33400 | 0.00000 | 0.00000 | 0.00000 | ; | H6- | C6- | C5A- | C9A |
| 12 | 11 | 13 | 14 | 3 | 30.33400 | 0.00000 | -30.33400 | 0.00000 | 0.00000 | 0.00000 | ; | H6- | C6- | C7- | C7M |
| 12 | 11 | 13 | 18 | 3 | 30.33400 | 0.00000 | -30.33400 | 0.00000 | 0.00000 | 0.00000 | ; | H6- | C6- | C7- | C8 |
| 13 | 11 | 10 | 25 | 3 | 30.33400 | 0.00000 | -30.33400 | 0.00000 | 0.00000 | 0.00000 | ; | C7- | C6- | C5A- | C9A |
| 13 | 18 | 19 | 20 | 3 | 0.00000 | 0.00000 | 0.00000 | 0.00000 | 0.00000 | 0.00000 | ; | C7- | C8- | C8M- | HM81 |
| 13 | 18 | 19 | 21 | 3 | 0.00000 | 0.00000 | 0.00000 | 0.00000 | 0.00000 | 0.00000 | ; | C7- | C8- | C8M- | HM82 |
| 13 | 18 | 19 | 22 | 3 | 0.00000 | 0.00000 | 0.00000 | 0.00000 | 0.00000 | 0.00000 | ; | C7- | C8- | C8M- | HM83 |
| 13 | 18 | 23 | 24 | 3 | 30.33400 | 0.00000 | -30.33400 | 0.00000 | 0.00000 | 0.00000 | ; | C7- | C8- | C9- | H9 |
| 13 | 18 | 23 | 25 | 3 | 30.33400 | 0.00000 | -30.33400 | 0.00000 | 0.00000 | 0.00000 | ; | C7- | C8- | C9- | C9A |
| 14 | 13 | 18 | 19 | 3 | 30.33400 | 0.00000 | -30.33400 | 0.00000 | 0.00000 | 0.00000 | ; | C7M- | C7- | C8- | C8M |
| 14 | 13 | 18 | 23 | 3 | 30.33400 | 0.00000 | -30.33400 | 0.00000 | 0.00000 | 0.00000 | ; | C7M- | C7- | C8- | C9 |
| 15 | 14 | 13 | 18 | 3 | 0.00000 | 0.00000 | 0.00000 | 0.00000 | 0.00000 | 0.00000 | ; | HM71- | C7M- | C7- | C8 |
| 16 | 14 | 13 | 18 | 3 | 0.00000 | 0.00000 | 0.00000 | 0.00000 | 0.00000 | 0.00000 | ; | HM72- | C7M- | C7- | C8 |
| 17 | 14 | 13 | 18 | 3 | 0.00000 | 0.00000 | 0.00000 | 0.00000 | 0.00000 | 0.00000 | ; | HM73- | C7M- | C7- | C8 |
| 18 | 23 | 25 | 26 | 3 | 30.33400 | 0.00000 | -30.33400 | 0.00000 | 0.00000 | 0.00000 | ; | C8- | C9- | C9A- | N10 |
| 19 | 18 | 23 | 24 | 3 | 30.33400 | 0.00000 | -30.33400 | 0.00000 | 0.00000 | 0.00000 | ; | C8M- | C8- | C9- | H9 |
| 19 | 18 | 23 | 25 | 3 | 30.33400 | 0.00000 | -30.33400 | 0.00000 | 0.00000 | 0.00000 | ; | C8M- | C8- | C9- | C9A |
| 20 | 19 | 18 | 23 | 3 | 0.00000 | 0.00000 | 0.00000 | 0.00000 | 0.00000 | 0.00000 | ; | HM81- | C8M- | C8- | C9 |
| 21 | 19 | 18 | 23 | 3 | 0.00000 | 0.00000 | 0.00000 | 0.00000 | 0.00000 | 0.00000 | ; | HM82- | C8M- | C8- | C9 |
| 22 | 19 | 18 | 23 | 3 | 0.00000 | 0.00000 | 0.00000 | 0.00000 | 0.00000 | 0.00000 | ; | HM83- | C8M- | C8- | C9 |
| 23 | 25 | 26 | 27 | 3 | 2.51040 | 0.00000 | -2.51040 | 0.00000 | 0.00000 | 0.00000 | ; | C9- | C9A- | N10- | C10 |
| 23 | 25 | 26 | 28 | 3 | 2.51040 | 0.00000 | -2.51040 | 0.00000 | 0.00000 | 0.00000 | ; | C9- | C9A- | N10- | C1' |
| 24 | 23 | 25 | 26 | 3 | 30.33400 | 0.00000 | -30.33400 | 0.00000 | 0.00000 | 0.00000 | ; | H9- | C9- | C9A- | N10 |
| 25 | 26 | 28 | 29 | 3 | 0.00000 | 0.00000 | 0.00000 | 0.00000 | 0.00000 | 0.00000 | ; | C9A- | N10- | C1'- | H1'1 |
| 25 | 26 | 28 | 30 | 3 | 0.00000 | 0.00000 | 0.00000 | 0.00000 | 0.00000 | 0.00000 | ; | C9A- | N10- | C1'- | H1'2 |
| 25 | 26 | 28 | 31 | 3 | 0.00000 | 0.00000 | 0.00000 | 0.00000 | 0.00000 | 0.00000 | ; | C9A- | N10- | C1'- | C2' |
| 26 | 28 | 31 | 32 | 3 | 0.65084 | 1.95253 | 0.00000 | -2.60338 | 0.00000 | 0.00000 | ; | N10- | C1'- | C2'- | H2' |
| 26 | 28 | 31 | 33 | 3 | 0.65084 | 1.95253 | 0.00000 | -2.60338 | 0.00000 | 0.00000 | ; | N10- | C1'- | C2'- | O2' |
| 26 | 28 | 31 | 35 | 3 | 0.65084 | 1.95253 | 0.00000 | -2.60338 | 0.00000 | 0.00000 | ; | N10- | C1'- | C2'- | C3' |
| 27 | 1 | 2 | 3 | 3 | 33.47200 | 0.00000 | -33.47200 | 0.00000 | 0.00000 | 0.00000 | ; | C10- | N1- | C2- | O2 |
| 27 | 1 | 2 | 4 | 3 | 33.47200 | 0.00000 | -33.47200 | 0.00000 | 0.00000 | 0.00000 | ; | C10- | N1- | C2- | N3 |
| 27 | 26 | 28 | 29 | 3 | 0.00000 | 0.00000 | 0.00000 | 0.00000 | 0.00000 | 0.00000 | ; | C10- | N10- | C1'- | H1'1 |
| 27 | 26 | 28 | 30 | 3 | 0.00000 | 0.00000 | 0.00000 | 0.00000 | 0.00000 | 0.00000 | ; | C10- | N10- | C1'- | H1'2 |
| 27 | 26 | 28 | 31 | 3 | 0.00000 | 0.00000 | 0.00000 | 0.00000 | 0.00000 | 0.00000 | ; | C10- | N10- | C1'- | C2' |
| 28 | 31 | 33 | 34 | 3 | 1.71544 | 0.96232 | 0.00000 | -2.67776 | 0.00000 | 0.00000 | ; | C1'- | C2'- | O2'- | HO2' |
| 28 | 31 | 35 | 36 | 3 | 0.65084 | 1.95253 | 0.00000 | -2.60338 | 0.00000 | 0.00000 | ; | C1'- | C2'- | C3'- | H3' |
| 28 | 31 | 35 | 37 | 3 | 0.65084 | 1.95253 | 0.00000 | -2.60338 | 0.00000 | 0.00000 | ; | C1'- | C2'- | C3'- | O3' |
| 28 | 31 | 35 | 39 | 3 | 3.68192 | 3.09616 | -2.09200 | -3.01248 | 0.00000 | 0.00000 | ; | C1'- | C2'- | C3'- | C4' |
| 29 | 28 | 31 | 32 | 3 | 0.65084 | 1.95253 | 0.00000 | -2.60338 | 0.00000 | 0.00000 | ; | H1'1- | C1'- | C2'- | H2' |
| 29 | 28 | 31 | 33 | 3 | 1.04600 | -1.04600 | 0.00000 | 0.00000 | 0.00000 | 0.00000 | ; | H1'1- | C1'- | C2'- | O2' |
| 29 | 28 | 31 | 35 | 3 | 0.65084 | 1.95253 | 0.00000 | -2.60338 | 0.00000 | 0.00000 | ; | H1'1- | C1'- | C2'- | C3' |
| 30 | 28 | 31 | 32 | 3 | 0.65084 | 1.95253 | 0.00000 | -2.60338 | 0.00000 | 0.00000 | ; | H1'2- | C1'- | C2'- | H2' |
| 30 | 28 | 31 | 33 | 3 | 1.04600 | -1.04600 | 0.00000 | 0.00000 | 0.00000 | 0.00000 | ; | H1'2- | C1'- | C2'- | O2' |
| 30 | 28 | 31 | 35 | 3 | 0.65084 | 1.95253 | 0.00000 | -2.60338 | 0.00000 | 0.00000 | ; | H1'2- | C1'- | C2'- | C3' |
| 31 | 35 | 37 | 38 | 3 | 1.71544 | 0.96232 | 0.00000 | -2.67776 | 0.00000 | 0.00000 | ; | C2'- | C3'- | O3'- | HO3' |
| 31 | 35 | 39 | 40 | 3 | 0.65084 | 1.95253 | 0.00000 | -2.60338 | 0.00000 | 0.00000 | ; | C2'- | C3'- | C4'- | H4' |
| 31 | 35 | 39 | 41 | 3 | 0.65084 | 1.95253 | 0.00000 | -2.60338 | 0.00000 | 0.00000 | ; | C2'- | C3'- | C4'- | O4' |
| 31 | 35 | 39 | 43 | 3 | 3.68192 | 3.09616 | -2.09200 | -3.01248 | 0.00000 | 0.00000 | ; | C2'- | C3'- | C4'- | C5' |
| 32 | 31 | 33 | 34 | 3 | 0.69733 | 2.09200 | 0.00000 | -2.78933 | 0.00000 | 0.00000 | ; | H2'- | C2'- | O2'- | HO2' |
| 32 | 31 | 35 | 36 | 3 | 0.65084 | 1.95253 | 0.00000 | -2.60338 | 0.00000 | 0.00000 | ; | H2'- | C2'- | C3'- | H3' |
| 32 | 31 | 35 | 37 | 3 | 1.04600 | -1.04600 | 0.00000 | 0.00000 | 0.00000 | 0.00000 | ; | H2'- | C2'- | C3'- | O3' |
| 32 | 31 | 35 | 39 | 3 | 0.65084 | 1.95253 | 0.00000 | -2.60338 | 0.00000 | 0.00000 | ; | H2'- | C2'- | C3'- | C4' |
| 33 | 31 | 35 | 36 | 3 | 1.04600 | -1.04600 | 0.00000 | 0.00000 | 0.00000 | 0.00000 | ; | O2'- | C2'- | C3'- | H3' |
| 33 | 31 | 35 | 37 | 3 | 0.60250 | 1.80749 | 9.83240 | -2.40998 | 0.00000 | 0.00000 | ; | O2'- | C2'- | C3'- | O3' |
| 33 | 31 | 35 | 39 | 3 | 0.65084 | 1.95253 | 0.00000 | -2.60338 | 0.00000 | 0.00000 | ; | O2'- | C2'- | C3'- | C4' |
| 34 | 33 | 31 | 35 | 3 | 1.71544 | 0.96232 | 0.00000 | -2.67776 | 0.00000 | 0.00000 | ; | HO2'- | O2'- | C2'- | C3' |
| 35 | 39 | 41 | 42 | 3 | 1.71544 | 0.96232 | 0.00000 | -2.67776 | 0.00000 | 0.00000 | ; | C3'- | C4'- | O4'- | HO4' |
| 35 | 39 | 43 | 44 | 3 | 0.65084 | 1.95253 | 0.00000 | -2.60338 | 0.00000 | 0.00000 | ; | C3'- | C4'- | C5'- | H5'1 |
| 35 | 39 | 43 | 45 | 3 | 0.65084 | 1.95253 | 0.00000 | -2.60338 | 0.00000 | 0.00000 | ; | C3'- | C4'- | C5'- | H5'2 |
| 35 | 39 | 43 | 46 | 3 | 0.65084 | 1.95253 | 0.00000 | -2.60338 | 0.00000 | 0.00000 | ; | C3'- | C4'- | C5'- | O5' |
| 36 | 35 | 37 | 38 | 3 | 0.69733 | 2.09200 | 0.00000 | -2.78933 | 0.00000 | 0.00000 | ; | H3'- | C3'- | O3'- | HO3' |
| 36 | 35 | 39 | 40 | 3 | 0.65084 | 1.95253 | 0.00000 | -2.60338 | 0.00000 | 0.00000 | ; | H3'- | C3'- | C4'- | H4' |
| 36 | 35 | 39 | 41 | 3 | 1.04600 | -1.04600 | 0.00000 | 0.00000 | 0.00000 | 0.00000 | ; | H3'- | C3'- | C4'- | O4' |
| 36 | 35 | 39 | 43 | 3 | 0.65084 | 1.95253 | 0.00000 | -2.60338 | 0.00000 | 0.00000 | ; | H3'- | C3'- | C4'- | C5' |
| 37 | 35 | 39 | 40 | 3 | 1.04600 | -1.04600 | 0.00000 | 0.00000 | 0.00000 | 0.00000 | ; | O3'- | C3'- | C4'- | H4' |
| 37 | 35 | 39 | 41 | 3 | 0.60250 | 1.80749 | 9.83240 | -2.40998 | 0.00000 | 0.00000 | ; | O3'- | C3'- | C4'- | O4' |
| 37 | 35 | 39 | 43 | 3 | 0.65084 | 1.95253 | 0.00000 | -2.60338 | 0.00000 | 0.00000 | ; | O3'- | C3'- | C4'- | C5' |
| 38 | 37 | 35 | 39 | 3 | 1.71544 | 0.96232 | 0.00000 | -2.67776 | 0.00000 | 0.00000 | ; | HO3'- | O3'- | C3'- | C4' |

```
    39      43      46      47       3    1.60387    4.81160    0.00000   -6.41547    0.00000    0.00000 ;    C4'-   C5'-   O5'-      P
    40      39      41      42       3    0.69733    2.09200    0.00000   -2.78933    0.00000    0.00000 ;    H4'-   C4'-   O4'-   HO4'
    40      39      43      44       3    0.65084    1.95253    0.00000   -2.60338    0.00000    0.00000 ;    H4'-   C4'-   C5'-   H5'1
    40      39      43      45       3    0.65084    1.95253    0.00000   -2.60338    0.00000    0.00000 ;    H4'-   C4'-   C5'-   H5'2
    40      39      43      46       3    1.04600   -1.04600    0.00000    0.00000    0.00000    0.00000 ;    H4'-   C4'-   C5'-    O5'
    41      39      43      44       3    1.04600   -1.04600    0.00000    0.00000    0.00000    0.00000 ;    O4'-   C4'-   C5'-   H5'1
    41      39      43      45       3    1.04600   -1.04600    0.00000    0.00000    0.00000    0.00000 ;    O4'-   C4'-   C5'-   H5'2
    41      39      43      46       3    0.60250    1.80749    9.83240   -2.40998    0.00000    0.00000 ;    O4'-   C4'-   C5'-    O5'
    42      41      39      43       3    1.71544    0.96232    0.00000   -2.67776    0.00000    0.00000 ;   HO4'-   O4'-   C4'-    C5'
    43      46      47      48       3    0.00000    0.00000    6.69440    0.00000    0.00000    0.00000 ;    C5'-   O5'-     P-    O1P
    43      46      47      49       3    0.00000    0.00000    6.69440    0.00000    0.00000    0.00000 ;    C5'-   O5'-     P-    O2P
    43      46      47      50       3    0.00000    0.00000    6.69440    0.00000    0.00000    0.00000 ;    C5'-   O5'-     P-    O3P
    44      43      46      47       3    1.60387    4.81160    0.00000   -6.41547    0.00000    0.00000 ;   H5'1-   C5'-   O5'-      P
    45      43      46      47       3    1.60387    4.81160    0.00000   -6.41547    0.00000    0.00000 ;   H5'2-   C5'-   O5'-      P

[ dihedrals ] ; impropers
; treated as propers in GROMACS to use correct AMBER analytical function
;    i      j      k      l   func   phase     kd      pn
     1     27     26      8      1   180.00   4.60240   2 ;     N1-   C10-   N10-   C4A
     2      6      4      5      1   180.00   4.60240   2 ;     C2-    C4-    N3-   HN3
     4      1      2      3      1   180.00  43.93200   2 ;     N3-    N1-    C2-    O2
     6     27      8      9      1   180.00   4.60240   2 ;     C4-   C10-   C4A-    N5
     8      4      6      7      1   180.00  43.93200   2 ;    C4A-    N3-    C4-    O4
    10     13     11     12      1   180.00   4.60240   2 ;    C5A-    C7-    C6-    H6
    10     23     25     26      1   180.00   4.60240   2 ;    C5A-    C9-   C9A-   N10
    11     18     13     14      1   180.00   4.60240   2 ;     C6-    C8-    C7-   C7M
    11     25     10      9      1   180.00   4.60240   2 ;     C6-   C9A-   C5A-    N5
    18     25     23     24      1   180.00   4.60240   2 ;     C8-   C9A-    C9-    H9
    19     13     18     23      1   180.00   4.60240   2 ;    C8M-    C7-    C8-    C9
    28     25     26     27      1   180.00   4.60240   2 ;    C1'-   C9A-   N10-   C10
```

#### PreQ1 Parameterization

```
[ atomtypes ]
;name   bond_type     mass     charge   ptype   sigma         epsilon       Amb
 n        n          0.00000  0.00000   A     3.25000e-01   7.11280e-01 ; 1.82  0.1700
 cc       cc         0.00000  0.00000   A     3.39967e-01   3.59824e-01 ; 1.91  0.0860
 nd       nd         0.00000  0.00000   A     3.25000e-01   7.11280e-01 ; 1.82  0.1700
 cd       cd         0.00000  0.00000   A     3.39967e-01   3.59824e-01 ; 1.91  0.0860
 c        c          0.00000  0.00000   A     3.39967e-01   3.59824e-01 ; 1.91  0.0860
 o        o          0.00000  0.00000   A     2.95992e-01   8.78640e-01 ; 1.66  0.2100
 c3       c3         0.00000  0.00000   A     3.39967e-01   4.57730e-01 ; 1.91  0.1094
 n4       n4         0.00000  0.00000   A     3.25000e-01   7.11280e-01 ; 1.82  0.1700
 na       na         0.00000  0.00000   A     3.25000e-01   7.11280e-01 ; 1.82  0.1700
 nh       nh         0.00000  0.00000   A     3.25000e-01   7.11280e-01 ; 1.82  0.1700
 hn       hn         0.00000  0.00000   A     1.06908e-01   6.56888e-02 ; 0.60  0.0157
 hx       hx         0.00000  0.00000   A     1.95998e-01   6.56888e-02 ; 1.10  0.0157
 h4       h4         0.00000  0.00000   A     2.51055e-01   6.27600e-02 ; 1.41  0.0150

[ moleculetype ]
;name            nrexcl
 PRF              3

[ atoms ]
;   nr  type  resi  res  atom  cgnr     charge      mass       ; qtot   bond_type
     1    n     1   PRF    N1    1    -0.812223     14.01000 ; qtot -0.812
     2   cc     1   PRF    C2    2     1.113785     12.01000 ; qtot 0.302
     3   nd     1   PRF    N3    3    -0.779199     14.01000 ; qtot -0.478
     4   cd     1   PRF    C4    4     0.507226     12.01000 ; qtot 0.030
     5   cc     1   PRF    C5    5    -0.310980     12.01000 ; qtot -0.281
     6    c     1   PRF    C6    6     0.771100     12.01000 ; qtot 0.490
     7    o     1   PRF    O6    7    -0.655883     16.00000 ; qtot -0.166
     8   cc     1   PRF    C7    8    -0.088930     12.01000 ; qtot -0.255
     9   c3     1   PRF   C10    9     0.165983     12.01000 ; qtot -0.089
    10   n4     1   PRF   N11   10    -0.619529     14.01000 ; qtot -0.709
    11   cd     1   PRF    C8   11    -0.277226     12.01000 ; qtot -0.986
    12   na     1   PRF    N9   12    -0.289640     14.01000 ; qtot -1.276
    13   nh     1   PRF    N2   13    -1.136486     14.01000 ; qtot -2.412
    14   hn     1   PRF    H1   14     0.451407      1.00800 ; qtot -1.961
    15   hx     1   PRF  H101   15     0.092254      1.00800 ; qtot -1.868
    16   hx     1   PRF  H102   16     0.092254      1.00800 ; qtot -1.776
    17   hn     1   PRF  H111   17     0.359643      1.00800 ; qtot -1.416
    18   hn     1   PRF  H112   18     0.359643      1.00800 ; qtot -1.057
    19   hn     1   PRF  H113   19     0.415401      1.00800 ; qtot -0.641
    20   h4     1   PRF    H8   20     0.253938      1.00800 ; qtot -0.387
    21   hn     1   PRF    H9   21     0.389385      1.00800 ; qtot 0.002
    22   hn     1   PRF   H21   22     0.499037      1.00800 ; qtot 0.501
    23   hn     1   PRF   H22   23     0.499037      1.00800 ; qtot 1.000

[ bonds ]
;   ai     aj funct   r             k
     1      2   1    1.3807e-01    3.5572e+05 ;     N1 - C2
     1      6   1    1.3789e-01    3.5782e+05 ;     N1 - C6
     1     14   1    1.0129e-01    3.3740e+05 ;     N1 - H1
     2      3   1    1.3172e-01    4.3965e+05 ;     C2 - N3
     2     13   1    1.3735e-01    3.6418e+05 ;     C2 - N2
     3      4   1    1.3694e-01    3.6911e+05 ;     N3 - C4
     4      5   1    1.3729e-01    4.1915e+05 ;     C4 - C5
     4     12   1    1.3802e-01    3.5631e+05 ;     C4 - N9
     5      6   1    1.4676e-01    3.1045e+05 ;     C5 - C6
     5      8   1    1.4278e-01    3.5129e+05 ;     C5 - C7
     6      7   1    1.2183e-01    5.3363e+05 ;     C6 - O6
     8      9   1    1.5015e-01    2.8016e+05 ;     C7 - C10
     8     11   1    1.3729e-01    4.1915e+05 ;     C7 - C8
     9     10   1    1.5110e-01    2.3707e+05 ;    C10 - N11
     9     15   1    1.0910e-01    2.8342e+05 ;    C10 - H101
     9     16   1    1.0910e-01    2.8342e+05 ;    C10 - H102
    10     17   1    1.0304e-01    3.1229e+05 ;    N11 - H111
    10     18   1    1.0304e-01    3.1229e+05 ;    N11 - H112
    10     19   1    1.0304e-01    3.1229e+05 ;    N11 - H113
    11     12   1    1.3802e-01    3.5631e+05 ;     C8 - N9
    11     20   1    1.0817e-01    2.9455e+05 ;     C8 - H8
```

```
    12     21   1    1.0100e-01    3.4175e+05 ;     N9 - H9
    13     22   1    1.0121e-01    3.3857e+05 ;     N2 - H21
    13     23   1    1.0121e-01    3.3857e+05 ;     N2 - H22

[ pairs ]
;   ai     aj    funct
     1      4      1 ;     N1 - C4
     1      8      1 ;     N1 - C7
     1     22      1 ;     N1 - H21
     1     23      1 ;     N1 - H22
     2      5      1 ;     C2 - C5
     2      7      1 ;     C2 - O6
     2     12      1 ;     C2 - N9
     3      8      1 ;     N3 - C7
     3     11      1 ;     N3 - C8
     3     21      1 ;     N3 - H9
     3     22      1 ;     N3 - H21
     3     23      1 ;     N3 - H22
     4      7      1 ;     C4 - O6
     4      9      1 ;     C4 - C10
     4     13      1 ;     C4 - N2
     4     20      1 ;     C4 - H8
     5     10      1 ;     C5 - N11
     5     15      1 ;     C5 - H101
     5     16      1 ;     C5 - H102
     5     20      1 ;     C5 - H8
     5     21      1 ;     C5 - H9
     6      3      1 ;     C6 - N3
     6      9      1 ;     C6 - C10
     6     11      1 ;     C6 - C8
     6     12      1 ;     C6 - N9
     6     13      1 ;     C6 - N2
     7      8      1 ;     O6 - C7
     8     17      1 ;     C7 - H111
     8     18      1 ;     C7 - H112
     8     19      1 ;     C7 - H113
     8     21      1 ;     C7 - H9
     9     12      1 ;    C10 - N9
     9     20      1 ;    C10 - H8
    10     11      1 ;    N11 - C8
    11     15      1 ;     C8 - H101
    11     16      1 ;     C8 - H102
    14      3      1 ;     H1 - N3
    14      5      1 ;     H1 - C5
    14      7      1 ;     H1 - O6
    14     13      1 ;     H1 - N2
    15     17      1 ;   H101 - H111
    15     18      1 ;   H101 - H112
    15     19      1 ;   H101 - H113
    16     17      1 ;   H102 - H111
    16     18      1 ;   H102 - H112
    16     19      1 ;   H102 - H113
    20     21      1 ;     H8 - H9

[ angles ]
;   ai     aj     ak    funct   theta         cth
     1      2      3      1    1.2300e+02    5.9831e+02 ;     N1 - C2     - N3
     1      2     13      1    1.1694e+02    6.0166e+02 ;     N1 - C2     - N2
     1      6      5      1    1.1270e+02    5.7823e+02 ;     N1 - C6     - C5
     1      6      7      1    1.2305e+02    6.2091e+02 ;     N1 - C6     - O6
     2      1      6      1    1.2327e+02    5.4141e+02 ;     C2 - N1     - C6
     2      1     14      1    1.1926e+02    4.0083e+02 ;     C2 - N1     - H1
     2      3      4      1    1.0549e+02    6.0082e+02 ;     C2 - N3     - C4
     2     13     22      1    1.1563e+02    4.0920e+02 ;     C2 - N2     - H21
     2     13     23      1    1.1563e+02    4.0920e+02 ;     C2 - N2     - H22
     3      2     13      1    1.2065e+02    6.0584e+02 ;     N3 - C2     - N2
     3      4      5      1    1.1165e+02    6.0417e+02 ;     N3 - C4     - C5
     3      4     12      1    1.2195e+02    5.9078e+02 ;     N3 - C4     - N9
     4      5      6      1    1.2135e+02    5.4476e+02 ;     C4 - C5     - C6
     4      5      8      1    1.1419e+02    5.7070e+02 ;     C4 - C5     - C7
     4     12     11      1    1.0990e+02    5.7321e+02 ;     C4 - N9     - C8
```

```
     4     12     21      1    1.2550e+02    3.9162e+02 ;     C4 - N9     - H9
     5      4     12      1    1.0699e+02    6.1421e+02 ;     C5 - C4     - N9
     5      6      7      1    1.2393e+02    5.7823e+02 ;     C5 - C6     - O6
     5      8      9      1    1.1597e+02    5.4057e+02 ;     C5 - C7     - C10
     5      8     11      1    1.1419e+02    5.7070e+02 ;     C5 - C7     - C8
     6      1     14      1    1.1755e+02    4.0417e+02 ;     C6 - N1     - H1
     6      5      8      1    1.2269e+02    5.3220e+02 ;     C6 - C5     - C7
     8      9     10      1    1.1558e+02    5.4057e+02 ;     C7 - C10    - N11
     8      9     15      1    1.1101e+02    3.9413e+02 ;     C7 - C10    - H101
     8      9     16      1    1.1101e+02    3.9413e+02 ;     C7 - C10    - H102
     8     11     12      1    1.0699e+02    6.1421e+02 ;     C7 - C8     - N9
     8     11     20      1    1.2848e+02    3.9581e+02 ;     C7 - C8     - H8
     9      8     11      1    1.1945e+02    5.4141e+02 ;    C10 - C7     - C8
     9     10     17      1    1.1011e+02    3.8409e+02 ;    C10 - N11    - H111
     9     10     18      1    1.1011e+02    3.8409e+02 ;    C10 - N11    - H112
     9     10     19      1    1.1011e+02    3.8409e+02 ;    C10 - N11    - H113
    10      9     15      1    1.0801e+02    4.0668e+02 ;    N11 - C10    - H101
    10      9     16      1    1.0801e+02    4.0668e+02 ;    N11 - C10    - H102
    11     12     21      1    1.2550e+02    3.9162e+02 ;     C8 - N9     - H9
    12     11     20      1    1.2053e+02    4.1673e+02 ;     N9 - C8     - H8
    15      9     16      1    1.0975e+02    3.2803e+02 ;   H101 - C10    - H102
    17     10     18      1    1.0830e+02    3.3974e+02 ;   H111 - N11    - H112
    17     10     19      1    1.0830e+02    3.3974e+02 ;   H111 - N11    - H113
    18     10     19      1    1.0830e+02    3.3974e+02 ;   H112 - N11    - H113
    22     13     23      1    1.1512e+02    3.3556e+02 ;    H21 - N2     - H22

[ dihedrals ] ; propers
; treated as RBs in GROMACS to use combine multiple AMBER torsions per quartet
;    i      j      k      l   func     C0         C1         C2         C3         C4         C5
     1      2      3      4      3   39.74800    0.00000  -39.74800    0.00000    0.00000    0.00000 ;     N1-    C2-    N3-    C4
     1      2     13     22      3    8.78640    0.00000   -8.78640    0.00000    0.00000    0.00000 ;     N1-    C2-    N2-   H21
     1      2     13     23      3    8.78640    0.00000   -8.78640    0.00000    0.00000    0.00000 ;     N1-    C2-    N2-   H22
     1      6      5      4      3   24.05800    0.00000  -24.05800    0.00000    0.00000    0.00000 ;     N1-    C6-    C5-    C4
     1      6      5      8      3   24.05800    0.00000  -24.05800    0.00000    0.00000    0.00000 ;     N1-    C6-    C5-    C7
     2      1      6      5      3   20.92000    0.00000  -20.92000    0.00000    0.00000    0.00000 ;     C2-    N1-    C6-    C5
     2      1      6      7      3   20.92000    0.00000  -20.92000    0.00000    0.00000    0.00000 ;     C2-    N1-    C6-    O6
     2      3      4      5      3   39.74800    0.00000  -39.74800    0.00000    0.00000    0.00000 ;     C2-    N3-    C4-    C5
     2      3      4     12      3   39.74800    0.00000  -39.74800    0.00000    0.00000    0.00000 ;     C2-    N3-    C4-    N9
     3      2     13     22      3    8.78640    0.00000   -8.78640    0.00000    0.00000    0.00000 ;     N3-    C2-    N2-   H21
     3      2     13     23      3    8.78640    0.00000   -8.78640    0.00000    0.00000    0.00000 ;     N3-    C2-    N2-   H22
     3      4      5      6      3   33.47200    0.00000  -33.47200    0.00000    0.00000    0.00000 ;     N3-    C4-    C5-    C6
     3      4      5      8      3   33.47200    0.00000  -33.47200    0.00000    0.00000    0.00000 ;     N3-    C4-    C5-    C7
     3      4     12     11      3   14.22560    0.00000  -14.22560    0.00000    0.00000    0.00000 ;     N3-    C4-    N9-    C8
     3      4     12     21      3   14.22560    0.00000  -14.22560    0.00000    0.00000    0.00000 ;     N3-    C4-    N9-    H9
     4      3      2     13      3   39.74800    0.00000  -39.74800    0.00000    0.00000    0.00000 ;     C4-    N3-    C2-    N2
     4      5      6      7      3   24.05800    0.00000  -24.05800    0.00000    0.00000    0.00000 ;     C4-    C5-    C6-    O6
     4      5      8      9      3   33.47200    0.00000  -33.47200    0.00000    0.00000    0.00000 ;     C4-    C5-    C7-   C10
     4      5      8     11      3   33.47200    0.00000  -33.47200    0.00000    0.00000    0.00000 ;     C4-    C5-    C7-    C8
     4     12     11      8      3   14.22560    0.00000  -14.22560    0.00000    0.00000    0.00000 ;     C4-    N9-    C8-    C7
     4     12     11     20      3   14.22560    0.00000  -14.22560    0.00000    0.00000    0.00000 ;     C4-    N9-    C8-    H8
     5      4     12     11      3   14.22560    0.00000  -14.22560    0.00000    0.00000    0.00000 ;     C5-    C4-    N9-    C8
     5      4     12     21      3   14.22560    0.00000  -14.22560    0.00000    0.00000    0.00000 ;     C5-    C4-    N9-    H9
     5      8      9     10      3    0.00000    0.00000    0.00000    0.00000    0.00000    0.00000 ;     C5-    C7-   C10-   N11
     5      8      9     15      3    0.00000    0.00000    0.00000    0.00000    0.00000    0.00000 ;     C5-    C7-   C10-  H101
     5      8      9     16      3    0.00000    0.00000    0.00000    0.00000    0.00000    0.00000 ;     C5-    C7-   C10-  H102
     5      8     11     12      3   33.47200    0.00000  -33.47200    0.00000    0.00000    0.00000 ;     C5-    C7-    C8-    N9
     5      8     11     20      3   33.47200    0.00000  -33.47200    0.00000    0.00000    0.00000 ;     C5-    C7-    C8-    H8
     6      1      2      3      3   13.80720    0.00000  -13.80720    0.00000    0.00000    0.00000 ;     C6-    N1-    C2-    N3
     6      1      2     13      3   13.80720    0.00000  -13.80720    0.00000    0.00000    0.00000 ;     C6-    N1-    C2-    N2
     6      5      4     12      3   33.47200    0.00000  -33.47200    0.00000    0.00000    0.00000 ;     C6-    C5-    C4-    N9
     6      5      8      9      3   33.47200    0.00000  -33.47200    0.00000    0.00000    0.00000 ;     C6-    C5-    C7-   C10
     6      5      8     11      3   33.47200    0.00000  -33.47200    0.00000    0.00000    0.00000 ;     C6-    C5-    C7-    C8
     7      6      5      8      3   24.05800    0.00000  -24.05800    0.00000    0.00000    0.00000 ;     O6-    C6-    C5-    C7
     8      5      4     12      3   33.47200    0.00000  -33.47200    0.00000    0.00000    0.00000 ;     C7-    C5-    C4-    N9
     8      9     10     17      3    0.65084    1.95253    0.00000   -2.60338    0.00000    0.00000 ;     C7-   C10-   N11-  H111
     8      9     10     18      3    0.65084    1.95253    0.00000   -2.60338    0.00000    0.00000 ;     C7-   C10-   N11-  H112
     8      9     10     19      3    0.65084    1.95253    0.00000   -2.60338    0.00000    0.00000 ;     C7-   C10-   N11-  H113
     8     11     12     21      3   14.22560    0.00000  -14.22560    0.00000    0.00000    0.00000 ;     C7-    C8-    N9-    H9
     9      8     11     12      3   33.47200    0.00000  -33.47200    0.00000    0.00000    0.00000 ;    C10-    C7-    C8-    N9
     9      8     11     20      3   33.47200    0.00000  -33.47200    0.00000    0.00000    0.00000 ;    C10-    C7-    C8-    H8
    10      9      8     11      3    0.00000    0.00000    0.00000    0.00000    0.00000    0.00000 ;    N11-   C10-    C7-    C8
    11      8      9     15      3    0.00000    0.00000    0.00000    0.00000    0.00000    0.00000 ;     C8-    C7-   C10-  H101
    11      8      9     16      3    0.00000    0.00000    0.00000    0.00000    0.00000    0.00000 ;     C8-    C7-   C10-  H102
    14      1      2      3      3   13.80720    0.00000  -13.80720    0.00000    0.00000    0.00000 ;     H1-    N1-    C2-    N3
    14      1      2     13      3   13.80720    0.00000  -13.80720    0.00000    0.00000    0.00000 ;     H1-    N1-    C2-    N2
    14      1      6      5      3   20.92000    0.00000  -20.92000    0.00000    0.00000    0.00000 ;     H1-    N1-    C6-    C5
    14      1      6      7      3   29.28800   -8.36800  -20.92000    0.00000    0.00000    0.00000 ;     H1-    N1-    C6-    O6
    15      9     10     17      3    0.65084    1.95253    0.00000   -2.60338    0.00000    0.00000 ;   H101-   C10-   N11-  H111
    15      9     10     18      3    0.65084    1.95253    0.00000   -2.60338    0.00000    0.00000 ;   H101-   C10-   N11-  H112
    15      9     10     19      3    0.65084    1.95253    0.00000   -2.60338    0.00000    0.00000 ;   H101-   C10-   N11-  H113
    16      9     10     17      3    0.65084    1.95253    0.00000   -2.60338    0.00000    0.00000 ;   H102-   C10-   N11-  H111
    16      9     10     18      3    0.65084    1.95253    0.00000   -2.60338    0.00000    0.00000 ;   H102-   C10-   N11-  H112
    16      9     10     19      3    0.65084    1.95253    0.00000   -2.60338    0.00000    0.00000 ;   H102-   C10-   N11-  H113
    20     11     12     21      3   14.22560    0.00000  -14.22560    0.00000    0.00000    0.00000 ;     H8-    C8-    N9-    H9

[ dihedrals ] ; impropers
; treated as propers in GROMACS to use correct AMBER analytical function
;    i      j      k      l   func   phase     kd      pn
     1      3      2     13      1   180.00   4.60240   2 ;     N1-    N3-    C2-    N2
```

| | | | | | | | | | | | | |
|---|---|---|---|---|---|---|---|---|---|---|---|---|
| 2 | 22 | 13 | 23 | 1 | 180.00 | 4.60240 | 2 | ; | C2- | H21- | N2- | H22 |
| 4 | 11 | 12 | 21 | 1 | 180.00 | 4.60240 | 2 | ; | C4- | C8- | N9- | H9 |
| 5 | 1 | 6 | 7 | 1 | 180.00 | 43.93200 | 2 | ; | C5- | N1- | C6- | O6 |
| 5 | 12 | 4 | 3 | 1 | 180.00 | 4.60240 | 2 | ; | C5- | N9- | C4- | N3 |
| 6 | 8 | 5 | 4 | 1 | 180.00 | 4.60240 | 2 | ; | C6- | C7- | C5- | C4 |
| 8 | 20 | 11 | 12 | 1 | 180.00 | 4.60240 | 2 | ; | C7- | H8- | C8- | N9 |
| 9 | 5 | 8 | 11 | 1 | 180.00 | 4.60240 | 2 | ; | C10- | C5- | C7- | C8 |
| 14 | 1 | 2 | 6 | 1 | 180.00 | 4.60240 | 2 | ; | H1- | N1- | C2- | C6 |

### Adenine Parameterization

```
[ atomtypes ]
;name   bond_type     mass     charge   ptype   sigma         epsilon       Amb
 na       na         0.00000  0.00000   A     3.25000e-01   7.11280e-01 ; 1.82  0.1700
 cc       cc         0.00000  0.00000   A     3.39967e-01   3.59824e-01 ; 1.91  0.0860
 nd       nd         0.00000  0.00000   A     3.25000e-01   7.11280e-01 ; 1.82  0.1700
 ca       ca         0.00000  0.00000   A     3.39967e-01   3.59824e-01 ; 1.91  0.0860
 nh       nh         0.00000  0.00000   A     3.25000e-01   7.11280e-01 ; 1.82  0.1700
 nb       nb         0.00000  0.00000   A     3.25000e-01   7.11280e-01 ; 1.82  0.1700
 hn       hn         0.00000  0.00000   A     1.06908e-01   6.56888e-02 ; 0.60  0.0157
 h5       h5         0.00000  0.00000   A     2.42146e-01   6.27600e-02 ; 1.36  0.0150

[ moleculetype ]
;name            nrexcl
 ADE              3

[ atoms ]
;   nr  type  resi  res  atom  cgnr     charge      mass       ; qtot   bond_type
     1   na     1   ADE    N9    1    -0.680395     14.01000 ; qtot -0.680
     2   cc     1   ADE    C8    2     0.320807     12.01000 ; qtot -0.360
     3   nd     1   ADE    N7    3    -0.601205     14.01000 ; qtot -0.961
     4   ca     1   ADE    C5    4    -0.072394     12.01000 ; qtot -1.033
     5   ca     1   ADE    C6    5     0.858578     12.01000 ; qtot -0.175
     6   nh     1   ADE    N6    6    -1.070396     14.01000 ; qtot -1.245
     7   nb     1   ADE    N1    7    -0.805178     14.01000 ; qtot -2.050
     8   ca     1   ADE    C2    8     0.506323     12.01000 ; qtot -1.544
     9   nb     1   ADE    N3    9    -0.788317     14.01000 ; qtot -2.332
    10   ca     1   ADE    C4   10     0.731612     12.01000 ; qtot -1.601
    11   hn     1   ADE    H9   11     0.435831      1.00800 ; qtot -1.165
    12   h5     1   ADE    H8   12     0.136268      1.00800 ; qtot -1.028
    13   hn     1   ADE   H61   13     0.463833      1.00800 ; qtot -0.565
    14   hn     1   ADE   H62   14     0.463833      1.00800 ; qtot -0.101
    15   h5     1   ADE    H2   15     0.100798      1.00800 ; qtot 0.000

[ bonds ]
;   ai     aj funct   r             k
     1      2  1    1.3802e-01    3.5631e+05 ;     N9 - C8
     1     10  1    1.3840e-01    3.5187e+05 ;     N9 - C4
     1     11  1    1.0100e-01    3.4175e+05 ;     N9 - H9
     2      3  1    1.3172e-01    4.3965e+05 ;     C8 - N7
     2     12  1    1.0819e-01    2.9430e+05 ;     C8 - H8
     3      4  1    1.3517e-01    3.9137e+05 ;     N7 - C5
     4      5  1    1.3984e-01    3.8585e+05 ;     C5 - C6
     4     10  1    1.3984e-01    3.8585e+05 ;     C5 - C4
     5      6  1    1.3859e-01    3.4970e+05 ;     C6 - N6
     5      7  1    1.3390e-01    4.0836e+05 ;     C6 - N1
     6     13  1    1.0121e-01    3.3857e+05 ;     N6 - H61
     6     14  1    1.0121e-01    3.3857e+05 ;     N6 - H62
     7      8  1    1.3390e-01    4.0836e+05 ;     N1 - C2
     8      9  1    1.3390e-01    4.0836e+05 ;     C2 - N3
     8     15  1    1.0878e-01    2.8719e+05 ;     C2 - H2
     9     10  1    1.3390e-01    4.0836e+05 ;     N3 - C4

[ pairs ]
;   ai     aj    funct
     1      5      1 ;     N9 - C6
     1      8      1 ;     N9 - C2
     2      5      1 ;     C8 - C6
     2      9      1 ;     C8 - N3
     3      6      1 ;     N7 - N6
     3      7      1 ;     N7 - N1
     3      9      1 ;     N7 - N3
     4      8      1 ;     C5 - C2
     4     12      1 ;     C5 - H8
     4     13      1 ;     C5 - H61
     4     14      1 ;     C5 - H62
     5      9      1 ;     C6 - N3
     5     15      1 ;     C6 - H2
     6      8      1 ;     N6 - C2
     6     10      1 ;     N6 - C4
```

```
     7     10      1 ;     N1 - C4
     7     13      1 ;     N1 - H61
     7     14      1 ;     N1 - H62
    10     12      1 ;     C4 - H8
    10     15      1 ;     C4 - H2
    11      3      1 ;     H9 - N7
    11      4      1 ;     H9 - C5
    11      9      1 ;     H9 - N3
    11     12      1 ;     H9 - H8
[ angles ]
;   ai     aj     ak    funct   theta         cth
     1      2      3      1    1.1222e+02    6.2676e+02 ;     N9 - C8     - N7
     1      2     12      1    1.2155e+02    4.1505e+02 ;     N9 - C8     - H8
     1     10      4      1    1.1834e+02    5.7823e+02 ;     N9 - C4     - C5
     1     10      9      1    1.2709e+02    5.8409e+02 ;     N9 - C4     - N3
     2      1     10      1    1.1315e+02    5.6400e+02 ;     C8 - N9     - C4
     2      1     11      1    1.2550e+02    3.9162e+02 ;     C8 - N9     - H9
     2      3      4      1    1.0488e+02    6.0668e+02 ;     C8 - N7     - C5
     3      2     12      1    1.2552e+02    4.2342e+02 ;     N7 - C8     - H8
     3      4      5      1    1.1972e+02    5.8158e+02 ;     N7 - C5     - C6
     3      4     10      1    1.1972e+02    5.8158e+02 ;     N7 - C5     - C4
     4      5      6      1    1.2095e+02    5.7153e+02 ;     C5 - C6     - N6
     4      5      7      1    1.2294e+02    5.7572e+02 ;     C5 - C6     - N1
     4     10      9      1    1.2294e+02    5.7572e+02 ;     C5 - C4     - N3
     5      4     10      1    1.2002e+02    5.5731e+02 ;     C6 - C5     - C4
     5      6     13      1    1.1607e+02    4.0501e+02 ;     C6 - N6     - H61
     5      6     14      1    1.1607e+02    4.0501e+02 ;     C6 - N6     - H62
     5      7      8      1    1.1722e+02    5.7153e+02 ;     C6 - N1     - C2
     6      5      7      1    1.1694e+02    6.0835e+02 ;     N6 - C6     - N1
     7      8      9      1    1.2726e+02    5.9329e+02 ;     N1 - C2     - N3
     7      8     15      1    1.1582e+02    4.3430e+02 ;     N1 - C2     - H2
     8      9     10      1    1.1722e+02    5.7153e+02 ;     C2 - N3     - C4
     9      8     15      1    1.1582e+02    4.3430e+02 ;     N3 - C2     - H2
    10      1     11      1    1.2554e+02    3.8995e+02 ;     C4 - N9     - H9
    13      6     14      1    1.1512e+02    3.3556e+02 ;    H61 - N6     - H62
[ dihedrals ] ; propers
; treated as RBs in GROMACS to use combine multiple AMBER torsions per quartet
;    i      j      k      l   func     C0         C1         C2         C3         C4         C5
     1      2      3      4      3   39.74800    0.00000  -39.74800    0.00000    0.00000    0.00000 ;     N9-    C8-    N7-    C5
     1     10      4      3      3   30.33400    0.00000  -30.33400    0.00000    0.00000    0.00000 ;     N9-    C4-    C5-    N7
     1     10      4      5      3   30.33400    0.00000  -30.33400    0.00000    0.00000    0.00000 ;     N9-    C4-    C5-    C6
     1     10      9      8      3   40.16640    0.00000  -40.16640    0.00000    0.00000    0.00000 ;     N9-    C4-    N3-    C2
     2      1     10      4      3    2.51040    0.00000   -2.51040    0.00000    0.00000    0.00000 ;     C8-    N9-    C4-    C5
     2      1     10      9      3    2.51040    0.00000   -2.51040    0.00000    0.00000    0.00000 ;     C8-    N9-    C4-    N3
     2      3      4      5      3   40.16640    0.00000  -40.16640    0.00000    0.00000    0.00000 ;     C8-    N7-    C5-    C6
     2      3      4     10      3   40.16640    0.00000  -40.16640    0.00000    0.00000    0.00000 ;     C8-    N7-    C5-    C4
     3      4      5      6      3   30.33400    0.00000  -30.33400    0.00000    0.00000    0.00000 ;     N7-    C5-    C6-    N6
     3      4      5      7      3   30.33400    0.00000  -30.33400    0.00000    0.00000    0.00000 ;     N7-    C5-    C6-    N1
     3      4     10      9      3   30.33400    0.00000  -30.33400    0.00000    0.00000    0.00000 ;     N7-    C5-    C4-    N3
     4      3      2     12      3   39.74800    0.00000  -39.74800    0.00000    0.00000    0.00000 ;     C5-    N7-    C8-    H8
     4      5      6     13      3    8.78640    0.00000   -8.78640    0.00000    0.00000    0.00000 ;     C5-    C6-    N6-   H61
     4      5      6     14      3    8.78640    0.00000   -8.78640    0.00000    0.00000    0.00000 ;     C5-    C6-    N6-   H62
     4      5      7      8      3   40.16640    0.00000  -40.16640    0.00000    0.00000    0.00000 ;     C5-    C6-    N1-    C2
     4     10      9      8      3   40.16640    0.00000  -40.16640    0.00000    0.00000    0.00000 ;     C5-    C4-    N3-    C2
     5      4     10      9      3   30.33400    0.00000  -30.33400    0.00000    0.00000    0.00000 ;     C6-    C5-    C4-    N3
     5      7      8      9      3   40.16640    0.00000  -40.16640    0.00000    0.00000    0.00000 ;     C6-    N1-    C2-    N3
     5      7      8     15      3   40.16640    0.00000  -40.16640    0.00000    0.00000    0.00000 ;     C6-    N1-    C2-    H2
     6      5      4     10      3   30.33400    0.00000  -30.33400    0.00000    0.00000    0.00000 ;     N6-    C6-    C5-    C4
     6      5      7      8      3   40.16640    0.00000  -40.16640    0.00000    0.00000    0.00000 ;     N6-    C6-    N1-    C2
     7      5      4     10      3   30.33400    0.00000  -30.33400    0.00000    0.00000    0.00000 ;     N1-    C6-    C5-    C4
     7      5      6     13      3    8.78640    0.00000   -8.78640    0.00000    0.00000    0.00000 ;     N1-    C6-    N6-   H61
     7      5      6     14      3    8.78640    0.00000   -8.78640    0.00000    0.00000    0.00000 ;     N1-    C6-    N6-   H62
     7      8      9     10      3   40.16640    0.00000  -40.16640    0.00000    0.00000    0.00000 ;     N1-    C2-    N3-    C4
    10      1      2      3      3   14.22560    0.00000  -14.22560    0.00000    0.00000    0.00000 ;     C4-    N9-    C8-    N7
    10      1      2     12      3   14.22560    0.00000  -14.22560    0.00000    0.00000    0.00000 ;     C4-    N9-    C8-    H8
    10      9      8     15      3   40.16640    0.00000  -40.16640    0.00000    0.00000    0.00000 ;     C4-    N3-    C2-    H2
    11      1      2      3      3   14.22560    0.00000  -14.22560    0.00000    0.00000    0.00000 ;     H9-    N9-    C8-    N7
    11      1      2     12      3   14.22560    0.00000  -14.22560    0.00000    0.00000    0.00000 ;     H9-    N9-    C8-    H8
    11      1     10      4      3    2.51040    0.00000   -2.51040    0.00000    0.00000    0.00000 ;     H9-    N9-    C4-    C5
    11      1     10      9      3    2.51040    0.00000   -2.51040    0.00000    0.00000    0.00000 ;     H9-    N9-    C4-    N3

[ dihedrals ] ; impropers
; treated as propers in GROMACS to use correct AMBER analytical function
;    i      j      k      l   func   phase     kd      pn
     4      1     10      9      1   180.00   4.60240   2 ;     C5-    N9-    C4-    N3
     4      7      5      6      1   180.00   4.60240   2 ;     C5-    N1-    C6-    N6
     5     10      4      3      1   180.00   4.60240   2 ;     C6-    C4-    C5-    N7
     5     13      6     14      1   180.00   4.60240   2 ;     C6-   H61-    N6-   H62
    11      1      2     10      1   180.00   4.60240   2 ;     H9-    N9-    C8-    C4
    12      1      2      3      1   180.00   4.60240   2 ;     H8-    N9-    C8-    N7
    15      7      8      9      1   180.00   4.60240   2 ;     H2-    N1-    C2-    N3
```

### Citrulline Parameterization

```
[ atomtypes ]
;name   bond_type     mass     charge   ptype   sigma         epsilon       Amb
 c2       c2          0.00000  0.00000   A     3.39967e-01   3.59824e-01 ; 1.91  0.0860
 o        o           0.00000  0.00000   A     2.95992e-01   8.78640e-01 ; 1.66  0.2100
 c3       c3          0.00000  0.00000   A     3.39967e-01   4.57730e-01 ; 1.91  0.1094
 n4       n4          0.00000  0.00000   A     3.25000e-01   7.11280e-01 ; 1.82  0.1700
 n        n           0.00000  0.00000   A     3.25000e-01   7.11280e-01 ; 1.82  0.1700
 c        c           0.00000  0.00000   A     3.39967e-01   3.59824e-01 ; 1.91  0.0860
 hx       hx          0.00000  0.00000   A     1.95998e-01   6.56888e-02 ; 1.10  0.0157
 hn       hn          0.00000  0.00000   A     1.06908e-01   6.56888e-02 ; 0.60  0.0157
 hc       hc          0.00000  0.00000   A     2.64953e-01   6.56888e-02 ; 1.49  0.0157
 h1       h1          0.00000  0.00000   A     2.47135e-01   6.56888e-02 ; 1.39  0.0157

[ moleculetype ]
;name            nrexcl
 CIR              3

[ atoms ]
;   nr  type  resi  res  atom  cgnr     charge      mass       ; qtot   bond_type
     1   c2     1   CIR    C1    1     0.744910     12.01000 ; qtot 0.745
     2    o     1   CIR    O1    2    -0.591837     16.00000 ; qtot 0.153
     3   c3     1   CIR    C2    3     0.387409     12.01000 ; qtot 0.540
     4   n4     1   CIR    N2    4    -1.051487     14.01000 ; qtot -0.511
     5    o     1   CIR    O2    5    -0.613828     16.00000 ; qtot -1.125
     6   c3     1   CIR    C3    6    -0.268176     12.01000 ; qtot -1.393
     7   c3     1   CIR    C4    7    -0.125750     12.01000 ; qtot -1.519
     8   c3     1   CIR    C5    8     0.496207     12.01000 ; qtot -1.023
     9    n     1   CIR    N6    9    -0.884485     14.01000 ; qtot -1.907
    10    c     1   CIR    C7   10     0.970314     12.01000 ; qtot -0.937
    11    o     1   CIR    O7   11    -0.663805     16.00000 ; qtot -1.601
    12    n     1   CIR    N8   12    -0.913355     14.01000 ; qtot -2.514
    13   hx     1   CIR    H2   13    -0.039685      1.00800 ; qtot -2.554
    14   hn     1   CIR    H6   14     0.382044      1.00800 ; qtot -2.172
    15   hn     1   CIR   H21   15     0.425394      1.00800 ; qtot -1.746
    16   hn     1   CIR   H22   16     0.390053      1.00800 ; qtot -1.356
    17   hn     1   CIR   H23   17     0.390053      1.00800 ; qtot -0.966
    18   hc     1   CIR   H31   18     0.089581      1.00800 ; qtot -0.876
    19   hc     1   CIR   H32   19     0.089581      1.00800 ; qtot -0.787
    20   hc     1   CIR   H41   20     0.066823      1.00800 ; qtot -0.720
    21   hc     1   CIR   H42   21     0.066823      1.00800 ; qtot -0.653
    22   h1     1   CIR   H51   22    -0.045110      1.00800 ; qtot -0.698
    23   h1     1   CIR   H52   23    -0.045110      1.00800 ; qtot -0.743
    24   hn     1   CIR   H81   24     0.371716      1.00800 ; qtot -0.372
    25   hn     1   CIR   H82   25     0.371716      1.00800 ; qtot 0.000

[ bonds ]
;   ai     aj funct   r             k
     1      2   1    1.2247e-01    5.2124e+05 ;     C1 - O1
     1      3   1    1.5095e-01    2.7347e+05 ;     C1 - C2
     1      5   1    1.2247e-01    5.2124e+05 ;     C1 - O2
     3      4   1    1.5110e-01    2.3707e+05 ;     C2 - N2
     3      6   1    1.5375e-01    2.5179e+05 ;     C2 - C3
     3     13   1    1.0910e-01    2.8342e+05 ;     C2 - H2
     4     15   1    1.0304e-01    3.1229e+05 ;     N2 - H21
     4     16   1    1.0304e-01    3.1229e+05 ;     N2 - H22
     4     17   1    1.0304e-01    3.1229e+05 ;     N2 - H23
     6      7   1    1.5375e-01    2.5179e+05 ;     C3 - C4
     6     18   1    1.0969e-01    2.7665e+05 ;     C3 - H31
     6     19   1    1.0969e-01    2.7665e+05 ;     C3 - H32
     7      8   1    1.5375e-01    2.5179e+05 ;     C4 - C5
     7     20   1    1.0969e-01    2.7665e+05 ;     C4 - H41
     7     21   1    1.0969e-01    2.7665e+05 ;     C4 - H42
     8      9   1    1.4619e-01    2.7506e+05 ;     C5 - N6
     8     22   1    1.0969e-01    2.7665e+05 ;     C5 - H51
     8     23   1    1.0969e-01    2.7665e+05 ;     C5 - H52
     9     10   1    1.3789e-01    3.5782e+05 ;     N6 - C7
     9     14   1    1.0129e-01    3.3740e+05 ;     N6 - H6
    10     11   1    1.2183e-01    5.3363e+05 ;     C7 - O7
    10     12   1    1.3789e-01    3.5782e+05 ;     C7 - N8
```

```
    12     24   1    1.0129e-01    3.3740e+05 ;     N8 - H81
    12     25   1    1.0129e-01    3.3740e+05 ;     N8 - H82

[ pairs ]
;   ai     aj    funct
     1      7      1 ;     C1 - C4
     1     15      1 ;     C1 - H21
     1     16      1 ;     C1 - H22
     1     17      1 ;     C1 - H23
     1     18      1 ;     C1 - H31
     1     19      1 ;     C1 - H32
     2      4      1 ;     O1 - N2
     2      6      1 ;     O1 - C3
     2     13      1 ;     O1 - H2
     3      8      1 ;     C2 - C5
     3     20      1 ;     C2 - H41
     3     21      1 ;     C2 - H42
     4      7      1 ;     N2 - C4
     4     18      1 ;     N2 - H31
     4     19      1 ;     N2 - H32
     5      4      1 ;     O2 - N2
     5      6      1 ;     O2 - C3
     5     13      1 ;     O2 - H2
     6      9      1 ;     C3 - N6
     6     15      1 ;     C3 - H21
     6     16      1 ;     C3 - H22
     6     17      1 ;     C3 - H23
     6     22      1 ;     C3 - H51
     6     23      1 ;     C3 - H52
     7     10      1 ;     C4 - C7
     7     13      1 ;     C4 - H2
     7     14      1 ;     C4 - H6
     8     11      1 ;     C5 - O7
     8     12      1 ;     C5 - N8
     8     18      1 ;     C5 - H31
     8     19      1 ;     C5 - H32
     9     20      1 ;     N6 - H41
     9     21      1 ;     N6 - H42
     9     24      1 ;     N6 - H81
     9     25      1 ;     N6 - H82
    10     22      1 ;     C7 - H51
    10     23      1 ;     C7 - H52
    11     14      1 ;     O7 - H6
    11     24      1 ;     O7 - H81
    11     25      1 ;     O7 - H82
    12     14      1 ;     N8 - H6
    13     15      1 ;     H2 - H21
    13     16      1 ;     H2 - H22
    13     17      1 ;     H2 - H23
    13     18      1 ;     H2 - H31
    13     19      1 ;     H2 - H32
    14     22      1 ;     H6 - H51
    14     23      1 ;     H6 - H52
    18     20      1 ;    H31 - H41
    18     21      1 ;    H31 - H42
    19     20      1 ;    H32 - H41
    19     21      1 ;    H32 - H42
    20     22      1 ;    H41 - H51
    20     23      1 ;    H41 - H52
    21     22      1 ;    H42 - H51
    21     23      1 ;    H42 - H52

[ angles ]
;   ai     aj     ak    funct   theta         cth
     1      3      4      1    1.1364e+02    5.4308e+02 ;     C1 - C2     - N2
     1      3      6      1    1.1156e+02    5.3053e+02 ;     C1 - C2     - C3
     1      3     13      1    1.1134e+02    3.9162e+02 ;     C1 - C2     - H2
     2      1      3      1    1.2282e+02    5.6819e+02 ;     O1 - C1     - C2
     2      1      5      1    1.2169e+02    6.7111e+02 ;     O1 - C1     - O2
     3      1      5      1    1.2282e+02    5.6819e+02 ;     C2 - C1     - O2
     3      4     15      1    1.1011e+02    3.8409e+02 ;     C2 - N2     - H21
```

```
    3      4     16      1    1.1011e+02    3.8409e+02 ;     C2 - N2     - H22
    3      4     17      1    1.1011e+02    3.8409e+02 ;     C2 - N2     - H23
    3      6      7      1    1.1151e+02    5.2635e+02 ;     C2 - C3     - C4
    3      6     18      1    1.0980e+02    3.8744e+02 ;     C2 - C3     - H31
    3      6     19      1    1.0980e+02    3.8744e+02 ;     C2 - C3     - H32
    4      3      6      1    1.1421e+02    5.3723e+02 ;     N2 - C2     - C3
    4      3     13      1    1.0801e+02    4.0668e+02 ;     N2 - C2     - H2
    6      3     13      1    1.1056e+02    3.8660e+02 ;     C3 - C2     - H2
    6      7      8      1    1.1151e+02    5.2635e+02 ;     C3 - C4     - C5
    6      7     20      1    1.0980e+02    3.8744e+02 ;     C3 - C4     - H41
    6      7     21      1    1.0980e+02    3.8744e+02 ;     C3 - C4     - H42
    7      6     18      1    1.0980e+02    3.8744e+02 ;     C4 - C3     - H31
    7      6     19      1    1.0980e+02    3.8744e+02 ;     C4 - C3     - H32
    7      8      9      1    1.1161e+02    5.5145e+02 ;     C4 - C5     - N6
    7      8     22      1    1.0956e+02    3.8828e+02 ;     C4 - C5     - H51
    7      8     23      1    1.0956e+02    3.8828e+02 ;     C4 - C5     - H52
    8      7     20      1    1.0980e+02    3.8744e+02 ;     C5 - C4     - H41
    8      7     21      1    1.0980e+02    3.8744e+02 ;     C5 - C4     - H42
    8      9     10      1    1.2069e+02    5.3053e+02 ;     C5 - N6     - C7
    8      9     14      1    1.1768e+02    3.8325e+02 ;     C5 - N6     - H6
    9      8     22      1    1.0888e+02    4.1673e+02 ;     N6 - C5     - H51
    9      8     23      1    1.0888e+02    4.1673e+02 ;     N6 - C5     - H52
    9     10     11      1    1.2305e+02    6.2091e+02 ;     N6 - C7     - O7
    9     10     12      1    1.1356e+02    6.1003e+02 ;     N6 - C7     - N8
   10      9     14      1    1.1755e+02    4.0417e+02 ;     C7 - N6     - H6
   10     12     24      1    1.1755e+02    4.0417e+02 ;     C7 - N8     - H81
   10     12     25      1    1.1755e+02    4.0417e+02 ;     C7 - N8     - H82
   11     10     12      1    1.2305e+02    6.2091e+02 ;     O7 - C7     - N8
   15      4     16      1    1.0830e+02    3.3974e+02 ;    H21 - N2     - H22
   15      4     17      1    1.0830e+02    3.3974e+02 ;    H21 - N2     - H23
   16      4     17      1    1.0830e+02    3.3974e+02 ;    H22 - N2     - H23
   18      6     19      1    1.0758e+02    3.2970e+02 ;    H31 - C3     - H32
   20      7     21      1    1.0758e+02    3.2970e+02 ;    H41 - C4     - H42
   22      8     23      1    1.0846e+02    3.2803e+02 ;    H51 - C5     - H52
   24     12     25      1    1.1795e+02    3.3137e+02 ;    H81 - N8     - H82

[ dihedrals ] ; propers
; treated as RBs in GROMACS to use combine multiple AMBER torsions per quartet
;    i      j      k      l   func     C0         C1         C2         C3         C4         C5
     1      3      4     15      3    0.65084    1.95253    0.00000   -2.60338    0.00000    0.00000 ;     C1-    C2-    N2-   H21
     1      3      4     16      3    0.65084    1.95253    0.00000   -2.60338    0.00000    0.00000 ;     C1-    C2-    N2-   H22
     1      3      4     17      3    0.65084    1.95253    0.00000   -2.60338    0.00000    0.00000 ;     C1-    C2-    N2-   H23
     1      3      6      7      3    0.65084    1.95253    0.00000   -2.60338    0.00000    0.00000 ;     C1-    C2-    C3-    C4
     1      3      6     18      3    0.65084    1.95253    0.00000   -2.60338    0.00000    0.00000 ;     C1-    C2-    C3-   H31
     1      3      6     19      3    0.65084    1.95253    0.00000   -2.60338    0.00000    0.00000 ;     C1-    C2-    C3-   H32
     2      1      3      4      3    0.00000    0.00000    0.00000    0.00000    0.00000    0.00000 ;     O1-    C1-    C2-    N2
     2      1      3      6      3    0.00000    0.00000    0.00000    0.00000    0.00000    0.00000 ;     O1-    C1-    C2-    C3
     2      1      3     13      3    0.00000    0.00000    0.00000    0.00000    0.00000    0.00000 ;     O1-    C1-    C2-    H2
     3      6      7      8      3    3.68192    3.09616   -2.09200   -3.01248    0.00000    0.00000 ;     C2-    C3-    C4-    C5
     3      6      7     20      3    0.66944    2.00832    0.00000   -2.67776    0.00000    0.00000 ;     C2-    C3-    C4-   H41
     3      6      7     21      3    0.66944    2.00832    0.00000   -2.67776    0.00000    0.00000 ;     C2-    C3-    C4-   H42
     4      3      6      7      3    0.65084    1.95253    0.00000   -2.60338    0.00000    0.00000 ;     N2-    C2-    C3-    C4
     4      3      6     18      3    0.65084    1.95253    0.00000   -2.60338    0.00000    0.00000 ;     N2-    C2-    C3-   H31
     4      3      6     19      3    0.65084    1.95253    0.00000   -2.60338    0.00000    0.00000 ;     N2-    C2-    C3-   H32
     5      1      3      4      3    0.00000    0.00000    0.00000    0.00000    0.00000    0.00000 ;     O2-    C1-    C2-    N2
     5      1      3      6      3    0.00000    0.00000    0.00000    0.00000    0.00000    0.00000 ;     O2-    C1-    C2-    C3
     5      1      3     13      3    0.00000    0.00000    0.00000    0.00000    0.00000    0.00000 ;     O2-    C1-    C2-    H2
     6      3      4     15      3    0.65084    1.95253    0.00000   -2.60338    0.00000    0.00000 ;     C3-    C2-    N2-   H21
     6      3      4     16      3    0.65084    1.95253    0.00000   -2.60338    0.00000    0.00000 ;     C3-    C2-    N2-   H22
     6      3      4     17      3    0.65084    1.95253    0.00000   -2.60338    0.00000    0.00000 ;     C3-    C2-    N2-   H23
     6      7      8      9      3    0.65084    1.95253    0.00000   -2.60338    0.00000    0.00000 ;     C3-    C4-    C5-    N6
     6      7      8     22      3    0.65084    1.95253    0.00000   -2.60338    0.00000    0.00000 ;     C3-    C4-    C5-   H51
     6      7      8     23      3    0.65084    1.95253    0.00000   -2.60338    0.00000    0.00000 ;     C3-    C4-    C5-   H52
     7      6      3     13      3    0.65084    1.95253    0.00000   -2.60338    0.00000    0.00000 ;     C4-    C3-    C2-    H2
     7      8      9     10      3    2.84512   -4.10032   16.73600    2.51040  -16.73600    0.00000 ;     C4-    C5-    N6-    C7
     7      8      9     14      3    0.00000    0.00000    0.00000    0.00000    0.00000    0.00000 ;     C4-    C5-    N6-    H6
     8      7      6     18      3    0.66944    2.00832    0.00000   -2.67776    0.00000    0.00000 ;     C5-    C4-    C3-   H31
     8      7      6     19      3    0.66944    2.00832    0.00000   -2.67776    0.00000    0.00000 ;     C5-    C4-    C3-   H32
     8      9     10     11      3   20.92000    0.00000  -20.92000    0.00000    0.00000    0.00000 ;     C5-    N6-    C7-    O7
     8      9     10     12      3   20.92000    0.00000  -20.92000    0.00000    0.00000    0.00000 ;     C5-    N6-    C7-    N8
     9      8      7     20      3    0.65084    1.95253    0.00000   -2.60338    0.00000    0.00000 ;     N6-    C5-    C4-   H41
     9      8      7     21      3    0.65084    1.95253    0.00000   -2.60338    0.00000    0.00000 ;     N6-    C5-    C4-   H42
     9     10     12     24      3   20.92000    0.00000  -20.92000    0.00000    0.00000    0.00000 ;     N6-    C7-    N8-   H81
     9     10     12     25      3   20.92000    0.00000  -20.92000    0.00000    0.00000    0.00000 ;     N6-    C7-    N8-   H82
    10      9      8     22      3    0.00000    0.00000    0.00000    0.00000    0.00000    0.00000 ;     C7-    N6-    C5-   H51
    10      9      8     23      3    0.00000    0.00000    0.00000    0.00000    0.00000    0.00000 ;     C7-    N6-    C5-   H52
    11     10      9     14      3   29.28800   -8.36800  -20.92000    0.00000    0.00000    0.00000 ;     O7-    C7-    N6-    H6
    11     10     12     24      3   29.28800   -8.36800  -20.92000    0.00000    0.00000    0.00000 ;     O7-    C7-    N8-   H81
    11     10     12     25      3   29.28800   -8.36800  -20.92000    0.00000    0.00000    0.00000 ;     O7-    C7-    N8-   H82
    12     10      9     14      3   20.92000    0.00000  -20.92000    0.00000    0.00000    0.00000 ;     N8-    C7-    N6-    H6
    13      3      4     15      3    0.65084    1.95253    0.00000   -2.60338    0.00000    0.00000 ;     H2-    C2-    N2-   H21
    13      3      4     16      3    0.65084    1.95253    0.00000   -2.60338    0.00000    0.00000 ;     H2-    C2-    N2-   H22
    13      3      4     17      3    0.65084    1.95253    0.00000   -2.60338    0.00000    0.00000 ;     H2-    C2-    N2-   H23
    13      3      6     18      3    0.65084    1.95253    0.00000   -2.60338    0.00000    0.00000 ;     H2-    C2-    C3-   H31
    13      3      6     19      3    0.65084    1.95253    0.00000   -2.60338    0.00000    0.00000 ;     H2-    C2-    C3-   H32
    14      9      8     22      3    0.00000    0.00000    0.00000    0.00000    0.00000    0.00000 ;     H6-    N6-    C5-   H51
```

```
    14      9      8     23      3    0.00000    0.00000    0.00000    0.00000    0.00000    0.00000 ;     H6-    N6-    C5-   H52
    18      6      7     20      3    0.62760    1.88280    0.00000   -2.51040    0.00000    0.00000 ;    H31-    C3-    C4-   H41
    18      6      7     21      3    0.62760    1.88280    0.00000   -2.51040    0.00000    0.00000 ;    H31-    C3-    C4-   H42
    19      6      7     20      3    0.62760    1.88280    0.00000   -2.51040    0.00000    0.00000 ;    H32-    C3-    C4-   H41
    19      6      7     21      3    0.62760    1.88280    0.00000   -2.51040    0.00000    0.00000 ;    H32-    C3-    C4-   H42
    20      7      8     22      3    0.65084    1.95253    0.00000   -2.60338    0.00000    0.00000 ;    H41-    C4-    C5-   H51
    20      7      8     23      3    0.65084    1.95253    0.00000   -2.60338    0.00000    0.00000 ;    H41-    C4-    C5-   H52
    21      7      8     22      3    0.65084    1.95253    0.00000   -2.60338    0.00000    0.00000 ;    H42-    C4-    C5-   H51
    21      7      8     23      3    0.65084    1.95253    0.00000   -2.60338    0.00000    0.00000 ;    H42-    C4-    C5-   H52

[ dihedrals ] ; impropers
; treated as propers in GROMACS to use correct AMBER analytical function
;    i      j       k      l   func   phase     kd      pn
     5      1       2      3      1   180.00   4.60240   2 ;     O2-    C1-    O1-    C2
     9     12      10     11      1   180.00  43.93200   2 ;     N6-    N8-    C7-    O7
    10      8       9     14      1   180.00   4.60240   2 ;     C7-    C5-    N6-    H6
    10     24      12     25      1   180.00   4.60240   2 ;     C7-   H81-    N8-   H82
```

### Arginine Parameterization

```
[ atomtypes ]
;name   bond_type     mass     charge   ptype   sigma         epsilon       Amb
 n4       n4          0.00000  0.00000   A     3.25000e-01   7.11280e-01 ; 1.82  0.1700
 c3       c3          0.00000  0.00000   A     3.39967e-01   4.57730e-01 ; 1.91  0.1094
 c2       c2          0.00000  0.00000   A     3.39967e-01   3.59824e-01 ; 1.91  0.0860
 o        o           0.00000  0.00000   A     2.95992e-01   8.78640e-01 ; 1.66  0.2100
 n3       n3          0.00000  0.00000   A     3.25000e-01   7.11280e-01 ; 1.82  0.1700
 cz       cz          0.00000  0.00000   A     3.39967e-01   3.59824e-01 ; 1.91  0.0860
 hn       hn          0.00000  0.00000   A     1.06908e-01   6.56888e-02 ; 0.60  0.0157
 hx       hx          0.00000  0.00000   A     1.95998e-01   6.56888e-02 ; 1.10  0.0157
 hc       hc          0.00000  0.00000   A     2.64953e-01   6.56888e-02 ; 1.49  0.0157
 h1       h1          0.00000  0.00000   A     2.47135e-01   6.56888e-02 ; 1.39  0.0157

[ moleculetype ]
;name            nrexcl
 ARG              3

[ atoms ]
;   nr  type  resi  res  atom  cgnr     charge      mass       ; qtot   bond_type
     1   n4     1   ARG     N    1    -1.031055     14.01000 ; qtot -1.031
     2   c3     1   ARG    CA    2     0.413143     12.01000 ; qtot -0.618
     3   c2     1   ARG     C    3     0.726624     12.01000 ; qtot 0.109
     4    o     1   ARG     O    4    -0.589459     16.00000 ; qtot -0.481
     5   c3     1   ARG    CB    5    -0.379750     12.01000 ; qtot -0.860
     6   c3     1   ARG    CG    6    -0.195282     12.01000 ; qtot -1.056
     7   c3     1   ARG    CD    7     0.403002     12.01000 ; qtot -0.653
     8   n3     1   ARG    NE    8    -0.914270     14.01000 ; qtot -1.567
     9   cz     1   ARG    CZ    9     1.112648     12.01000 ; qtot -0.454
    10   n3     1   ARG   NH1   10    -1.040737     14.01000 ; qtot -1.495
    11   n3     1   ARG   NH2   11    -1.040737     14.01000 ; qtot -2.536
    12    o     1   ARG   OXT   12    -0.577815     16.00000 ; qtot -3.114
    13   hn     1   ARG    H1   13     0.444131      1.00800 ; qtot -2.670
    14   hn     1   ARG    H2   14     0.400624      1.00800 ; qtot -2.269
    15   hn     1   ARG    H3   15     0.400624      1.00800 ; qtot -1.868
    16   hx     1   ARG    HA   16    -0.023402      1.00800 ; qtot -1.892
    17   hc     1   ARG   HB2   17     0.135922      1.00800 ; qtot -1.756
    18   hc     1   ARG   HB3   18     0.135922      1.00800 ; qtot -1.620
    19   hc     1   ARG   HG2   19     0.102761      1.00800 ; qtot -1.517
    20   hc     1   ARG   HG3   20     0.102761      1.00800 ; qtot -1.414
    21   h1     1   ARG   HD2   21     0.026011      1.00800 ; qtot -1.388
    22   h1     1   ARG   HD3   22     0.026011      1.00800 ; qtot -1.362
    23   hn     1   ARG    HE   23     0.450098      1.00800 ; qtot -0.912
    24   hn     1   ARG  HH11   24     0.478055      1.00800 ; qtot -0.434
    25   hn     1   ARG  HH12   25     0.478055      1.00800 ; qtot 0.044
    26   hn     1   ARG  HH21   26     0.478055      1.00800 ; qtot 0.522
    27   hn     1   ARG  HH22   27     0.478055      1.00800 ; qtot 1.000

[ bonds ]
;   ai     aj  funct   r             k
     1      2   1    1.5110e-01    2.3707e+05 ;      N - CA
     1     13   1    1.0304e-01    3.1229e+05 ;      N - H1
     1     14   1    1.0304e-01    3.1229e+05 ;      N - H2
     1     15   1    1.0304e-01    3.1229e+05 ;      N - H3
     2      3   1    1.5095e-01    2.7347e+05 ;     CA - C
     2      5   1    1.5375e-01    2.5179e+05 ;     CA - CB
     2     16   1    1.0910e-01    2.8342e+05 ;     CA - HA
     3      4   1    1.2247e-01    5.2124e+05 ;      C - O
     3     12   1    1.2247e-01    5.2124e+05 ;      C - OXT
     5      6   1    1.5375e-01    2.5179e+05 ;     CB - CG
     5     17   1    1.0969e-01    2.7665e+05 ;     CB - HB2
     5     18   1    1.0969e-01    2.7665e+05 ;     CB - HB3
     6      7   1    1.5375e-01    2.5179e+05 ;     CG - CD
     6     19   1    1.0969e-01    2.7665e+05 ;     CG - HG2
     6     20   1    1.0969e-01    2.7665e+05 ;     CG - HG3
     7      8   1    1.4647e-01    2.7271e+05 ;     CD - NE
     7     21   1    1.0969e-01    2.7665e+05 ;     CD - HD2
     7     22   1    1.0969e-01    2.7665e+05 ;     CD - HD3
     8      9   1    1.3400e-01    4.0694e+05 ;     NE - CZ
     8     23   1    1.0190e-01    3.2836e+05 ;     NE - HE
```

```
     9     10   1    1.3400e-01    4.0694e+05 ;      CZ - NH1
     9     11   1    1.3400e-01    4.0694e+05 ;      CZ - NH2
    10     24   1    1.0190e-01    3.2836e+05 ;    NH1 - HH11
    10     25   1    1.0190e-01    3.2836e+05 ;    NH1 - HH12
    11     26   1    1.0190e-01    3.2836e+05 ;    NH2 - HH21
    11     27   1    1.0190e-01    3.2836e+05 ;    NH2 - HH22

[ pairs ]
;   ai     aj    funct
     1      4      1 ;      N - O
     1      6      1 ;      N - CG
     1     12      1 ;      N - OXT
     1     17      1 ;      N - HB2
     1     18      1 ;      N - HB3
     2      7      1 ;     CA - CD
     2     19      1 ;     CA - HG2
     2     20      1 ;     CA - HG3
     3      6      1 ;      C - CG
     3     17      1 ;      C - HB2
     3     18      1 ;      C - HB3
     4      5      1 ;      O - CB
     4     16      1 ;      O - HA
     5      8      1 ;     CB - NE
     5     12      1 ;     CB - OXT
     5     21      1 ;     CB - HD2
     5     22      1 ;     CB - HD3
     6      9      1 ;     CG - CZ
     6     16      1 ;     CG - HA
     6     23      1 ;     CG - HE
     7     10      1 ;     CD - NH1
     7     11      1 ;     CD - NH2
     7     17      1 ;     CD - HB2
     7     18      1 ;     CD - HB3
     8     19      1 ;     NE - HG2
     8     20      1 ;     NE - HG3
     8     24      1 ;     NE - HH11
     8     25      1 ;     NE - HH12
     8     26      1 ;     NE - HH21
     8     27      1 ;     NE - HH22
     9     21      1 ;     CZ - HD2
     9     22      1 ;     CZ - HD3
    10     23      1 ;    NH1 - HE
    10     26      1 ;    NH1 - HH21
    10     27      1 ;    NH1 - HH22
    11     23      1 ;    NH2 - HE
    11     24      1 ;    NH2 - HH11
    11     25      1 ;    NH2 - HH12
    12     16      1 ;    OXT - HA
    13      3      1 ;     H1 - C
    13      5      1 ;     H1 - CB
    13     16      1 ;     H1 - HA
    14      3      1 ;     H2 - C
    14      5      1 ;     H2 - CB
    14     16      1 ;     H2 - HA
    15      3      1 ;     H3 - C
    15      5      1 ;     H3 - CB
    15     16      1 ;     H3 - HA
    16     17      1 ;     HA - HB2
    16     18      1 ;     HA - HB3
    17     19      1 ;    HB2 - HG2
    17     20      1 ;    HB2 - HG3
    18     19      1 ;    HB3 - HG2
    18     20      1 ;    HB3 - HG3
    19     21      1 ;    HG2 - HD2
    19     22      1 ;    HG2 - HD3
    20     21      1 ;    HG3 - HD2
    20     22      1 ;    HG3 - HD3
    21     23      1 ;    HD2 - HE
    22     23      1 ;    HD3 - HE

[ angles ]
```

```
;   ai     aj     ak    funct   theta         cth
     1      2      3      1    1.1364e+02    5.4308e+02 ;      N - CA     - C
     1      2      5      1    1.1421e+02    5.3723e+02 ;      N - CA     - CB
     1      2     16      1    1.0801e+02    4.0668e+02 ;      N - CA     - HA
     2      1     13      1    1.1011e+02    3.8409e+02 ;     CA - N      - H1
     2      1     14      1    1.1011e+02    3.8409e+02 ;     CA - N      - H2
     2      1     15      1    1.1011e+02    3.8409e+02 ;     CA - N      - H3
     2      3      4      1    1.2282e+02    5.6819e+02 ;     CA - C      - O
     2      3     12      1    1.2282e+02    5.6819e+02 ;     CA - C      - OXT
     2      5      6      1    1.1151e+02    5.2635e+02 ;     CA - CB     - CG
     2      5     17      1    1.0980e+02    3.8744e+02 ;     CA - CB     - HB2
     2      5     18      1    1.0980e+02    3.8744e+02 ;     CA - CB     - HB3
     3      2      5      1    1.1156e+02    5.3053e+02 ;      C - CA     - CB
     3      2     16      1    1.1134e+02    3.9162e+02 ;      C - CA     - HA
     4      3     12      1    1.2169e+02    6.7111e+02 ;      O - C      - OXT
     5      2     16      1    1.1056e+02    3.8660e+02 ;     CB - CA     - HA
     5      6      7      1    1.1151e+02    5.2635e+02 ;     CB - CG     - CD
     5      6     19      1    1.0980e+02    3.8744e+02 ;     CB - CG     - HG2
     5      6     20      1    1.0980e+02    3.8744e+02 ;     CB - CG     - HG3
     6      5     17      1    1.0980e+02    3.8744e+02 ;     CG - CB     - HB2
     6      5     18      1    1.0980e+02    3.8744e+02 ;     CG - CB     - HB3
     6      7      8      1    1.1104e+02    5.5229e+02 ;     CG - CD     - NE
     6      7     21      1    1.0956e+02    3.8828e+02 ;     CG - CD     - HD2
     6      7     22      1    1.0956e+02    3.8828e+02 ;     CG - CD     - HD3
     7      6     19      1    1.0980e+02    3.8744e+02 ;     CD - CG     - HG2
     7      6     20      1    1.0980e+02    3.8744e+02 ;     CD - CG     - HG3
     7      8      9      1    1.1852e+02    5.4096e+02 ;     CD - NE     - CZ
     7      8     23      1    1.0929e+02    3.9664e+02 ;     CD - NE     - HE
     8      7     21      1    1.0988e+02    4.1422e+02 ;     NE - CD     - HD2
     8      7     22      1    1.0988e+02    4.1422e+02 ;     NE - CD     - HD3
     8      9     10      1    1.2014e+02    6.1086e+02 ;     NE - CZ     - NH1
     8      9     11      1    1.2014e+02    6.1086e+02 ;     NE - CZ     - NH2
     9      8     23      1    1.1938e+02    4.1087e+02 ;     CZ - NE     - HE
     9     10     24      1    1.1938e+02    4.1087e+02 ;     CZ - NH1    - HH11
     9     10     25      1    1.1938e+02    4.1087e+02 ;     CZ - NH1    - HH12
     9     11     26      1    1.1938e+02    4.1087e+02 ;     CZ - NH2    - HH21
     9     11     27      1    1.1938e+02    4.1087e+02 ;     CZ - NH2    - HH22
    10      9     11      1    1.2014e+02    6.1086e+02 ;    NH1 - CZ     - NH2
    13      1     14      1    1.0830e+02    3.3974e+02 ;     H1 - N      - H2
    13      1     15      1    1.0830e+02    3.3974e+02 ;     H1 - N      - H3
    14      1     15      1    1.0830e+02    3.3974e+02 ;     H2 - N      - H3
    17      5     18      1    1.0758e+02    3.2970e+02 ;    HB2 - CB     - HB3
    19      6     20      1    1.0758e+02    3.2970e+02 ;    HG2 - CG     - HG3
    21      7     22      1    1.0846e+02    3.2803e+02 ;    HD2 - CD     - HD3
    24     10     25      1    1.0640e+02    3.4644e+02 ;   HH11 - NH1    - HH12
    26     11     27      1    1.0640e+02    3.4644e+02 ;   HH21 - NH2    - HH22

[ dihedrals ] ; propers
; treated as RBs in GROMACS to use combine multiple AMBER torsions per quartet
;    i      j      k      l   func     C0         C1         C2         C3         C4         C5
     1      2      3      4      3    0.00000    0.00000    0.00000    0.00000    0.00000    0.00000 ;      N-    CA-     C-     O
     1      2      3     12      3    0.00000    0.00000    0.00000    0.00000    0.00000    0.00000 ;      N-    CA-     C-   OXT
     1      2      5      6      3    0.65084    1.95253    0.00000   -2.60338    0.00000    0.00000 ;      N-    CA-    CB-    CG
     1      2      5     17      3    0.65084    1.95253    0.00000   -2.60338    0.00000    0.00000 ;      N-    CA-    CB-   HB2
     1      2      5     18      3    0.65084    1.95253    0.00000   -2.60338    0.00000    0.00000 ;      N-    CA-    CB-   HB3
     2      5      6      7      3    3.68192    3.09616   -2.09200   -3.01248    0.00000    0.00000 ;    CA-    CB-    CG-    CD
     2      5      6     19      3    0.66944    2.00832    0.00000   -2.67776    0.00000    0.00000 ;    CA-    CB-    CG-   HG2
     2      5      6     20      3    0.66944    2.00832    0.00000   -2.67776    0.00000    0.00000 ;    CA-    CB-    CG-   HG3
     3      2      5      6      3    0.65084    1.95253    0.00000   -2.60338    0.00000    0.00000 ;     C-    CA-    CB-    CG
     3      2      5     17      3    0.65084    1.95253    0.00000   -2.60338    0.00000    0.00000 ;     C-    CA-    CB-   HB2
     3      2      5     18      3    0.65084    1.95253    0.00000   -2.60338    0.00000    0.00000 ;     C-    CA-    CB-   HB3
     4      3      2      5      3    0.00000    0.00000    0.00000    0.00000    0.00000    0.00000 ;     O-     C-    CA-    CB
     4      3      2     16      3    0.00000    0.00000    0.00000    0.00000    0.00000    0.00000 ;     O-     C-    CA-    HA
     5      2      3     12      3    0.00000    0.00000    0.00000    0.00000    0.00000    0.00000 ;    CB-    CA-     C-   OXT
     5      6      7      8      3    0.65084    1.95253    0.00000   -2.60338    0.00000    0.00000 ;    CB-    CG-    CD-    NE
     5      6      7     21      3    0.65084    1.95253    0.00000   -2.60338    0.00000    0.00000 ;    CB-    CG-    CD-   HD2
     5      6      7     22      3    0.65084    1.95253    0.00000   -2.60338    0.00000    0.00000 ;    CB-    CG-    CD-   HD3
     6      5      2     16      3    0.65084    1.95253    0.00000   -2.60338    0.00000    0.00000 ;    CG-    CB-    CA-    HA
     6      7      8      9      3    1.25520    3.76560    0.00000   -5.02080    0.00000    0.00000 ;    CG-    CD-    NE-    CZ
     6      7      8     23      3    1.25520    3.76560    0.00000   -5.02080    0.00000    0.00000 ;    CG-    CD-    NE-    HE
     7      6      5     17      3    0.66944    2.00832    0.00000   -2.67776    0.00000    0.00000 ;    CD-    CG-    CB-   HB2
     7      6      5     18      3    0.66944    2.00832    0.00000   -2.67776    0.00000    0.00000 ;    CD-    CG-    CB-   HB3
     7      8      9     10      3    2.51040    0.00000   -2.51040    0.00000    0.00000    0.00000 ;    CD-    NE-    CZ-   NH1
     7      8      9     11      3    2.51040    0.00000   -2.51040    0.00000    0.00000    0.00000 ;    CD-    NE-    CZ-   NH2
     8      7      6     19      3    0.65084    1.95253    0.00000   -2.60338    0.00000    0.00000 ;    NE-    CD-    CG-   HG2
     8      7      6     20      3    0.65084    1.95253    0.00000   -2.60338    0.00000    0.00000 ;    NE-    CD-    CG-   HG3
     8      9     10     24      3    2.51040    0.00000   -2.51040    0.00000    0.00000    0.00000 ;    NE-    CZ-   NH1-  HH11
     8      9     10     25      3    2.51040    0.00000   -2.51040    0.00000    0.00000    0.00000 ;    NE-    CZ-   NH1-  HH12
     8      9     11     26      3    2.51040    0.00000   -2.51040    0.00000    0.00000    0.00000 ;    NE-    CZ-   NH2-  HH21
     8      9     11     27      3    2.51040    0.00000   -2.51040    0.00000    0.00000    0.00000 ;    NE-    CZ-   NH2-  HH22
     9      8      7     21      3    1.25520    3.76560    0.00000   -5.02080    0.00000    0.00000 ;    CZ-    NE-    CD-   HD2
```

```
    9      8      7     22      3    1.25520    3.76560    0.00000   -5.02080    0.00000    0.00000 ;     CZ-    NE-    CD-   HD3
   10      9      8     23      3    2.51040    0.00000   -2.51040    0.00000    0.00000    0.00000 ;    NH1-    CZ-    NE-    HE
   10      9     11     26      3    2.51040    0.00000   -2.51040    0.00000    0.00000    0.00000 ;    NH1-    CZ-   NH2-  HH21
   10      9     11     27      3    2.51040    0.00000   -2.51040    0.00000    0.00000    0.00000 ;    NH1-    CZ-   NH2-  HH22
   11      9      8     23      3    2.51040    0.00000   -2.51040    0.00000    0.00000    0.00000 ;    NH2-    CZ-    NE-    HE
   11      9     10     24      3    2.51040    0.00000   -2.51040    0.00000    0.00000    0.00000 ;    NH2-    CZ-   NH1-  HH11
   11      9     10     25      3    2.51040    0.00000   -2.51040    0.00000    0.00000    0.00000 ;    NH2-    CZ-   NH1-  HH12
   12      3      2     16      3    0.00000    0.00000    0.00000    0.00000    0.00000    0.00000 ;    OXT-     C-    CA-    HA
   13      1      2      3      3    0.65084    1.95253    0.00000   -2.60338    0.00000    0.00000 ;     H1-     N-    CA-     C
   13      1      2      5      3    0.65084    1.95253    0.00000   -2.60338    0.00000    0.00000 ;     H1-     N-    CA-    CB
   13      1      2     16      3    0.65084    1.95253    0.00000   -2.60338    0.00000    0.00000 ;     H1-     N-    CA-    HA
   14      1      2      3      3    0.65084    1.95253    0.00000   -2.60338    0.00000    0.00000 ;     H2-     N-    CA-     C
   14      1      2      5      3    0.65084    1.95253    0.00000   -2.60338    0.00000    0.00000 ;     H2-     N-    CA-    CB
   14      1      2     16      3    0.65084    1.95253    0.00000   -2.60338    0.00000    0.00000 ;     H2-     N-    CA-    HA
   15      1      2      3      3    0.65084    1.95253    0.00000   -2.60338    0.00000    0.00000 ;     H3-     N-    CA-     C
   15      1      2      5      3    0.65084    1.95253    0.00000   -2.60338    0.00000    0.00000 ;     H3-     N-    CA-    CB
   15      1      2     16      3    0.65084    1.95253    0.00000   -2.60338    0.00000    0.00000 ;     H3-     N-    CA-    HA
   16      2      5     17      3    0.65084    1.95253    0.00000   -2.60338    0.00000    0.00000 ;     HA-    CA-    CB-   HB2
   16      2      5     18      3    0.65084    1.95253    0.00000   -2.60338    0.00000    0.00000 ;     HA-    CA-    CB-   HB3
   17      5      6     19      3    0.62760    1.88280    0.00000   -2.51040    0.00000    0.00000 ;    HB2-    CB-    CG-   HG2
   17      5      6     20      3    0.62760    1.88280    0.00000   -2.51040    0.00000    0.00000 ;    HB2-    CB-    CG-   HG3
   18      5      6     19      3    0.62760    1.88280    0.00000   -2.51040    0.00000    0.00000 ;    HB3-    CB-    CG-   HG2
   18      5      6     20      3    0.62760    1.88280    0.00000   -2.51040    0.00000    0.00000 ;    HB3-    CB-    CG-   HG3
   19      6      7     21      3    0.65084    1.95253    0.00000   -2.60338    0.00000    0.00000 ;    HG2-    CG-    CD-   HD2
   19      6      7     22      3    0.65084    1.95253    0.00000   -2.60338    0.00000    0.00000 ;    HG2-    CG-    CD-   HD3
   20      6      7     21      3    0.65084    1.95253    0.00000   -2.60338    0.00000    0.00000 ;    HG3-    CG-    CD-   HD2
   20      6      7     22      3    0.65084    1.95253    0.00000   -2.60338    0.00000    0.00000 ;    HG3-    CG-    CD-   HD3
   21      7      8     23      3    1.25520    3.76560    0.00000   -5.02080    0.00000    0.00000 ;    HD2-    CD-    NE-    HE
   22      7      8     23      3    1.25520    3.76560    0.00000   -5.02080    0.00000    0.00000 ;    HD3-    CD-    NE-    HE

[ dihedrals ] ; impropers
; treated as propers in GROMACS to use correct AMBER analytical function
;    i      j      k      l   func   phase     kd      pn
     2      4      3     12      1   180.00   4.60240   2 ;     CA-     O-     C-   OXT
     8     10      9     11      1   180.00   4.60240   2 ;     NE-   NH1-    CZ-   NH2
```
